# Biotechnological potential of aromatic compounds–utilizing bacteria from Brazilian caves, including a novel cave *Nocardioides sp*. SF1

**DOI:** 10.64898/2026.06.23.734003

**Authors:** Eric de Lima Silva Marques, Eduardo Gross, Isis Caroline de Amorim Jambeiro, Maria Clara Bessa Souza, João Carlos Teixeira Dias, Rachel Passos Rezende

## Abstract

From Brazilian limestone caves, we isolated 29 bacteria utilizing phenol (23 bacteria), toluene (all bacteria), and/or benzene (all bacteria) as sole carbon sources. One isolate showed phosphate solubilization, while lipase/esterase activity occurred in two isolates; no amylase activity was detected, but 16 isolates (∼55%) exhibited protease activity. Among them, *Nocardioides* sp. SF1 was selected for whole-genome sequencing due to its aromatic compound tolerance and protease activity. Additionally, catechol cleavage assays yielded unexpected purple pigmentation, suggesting non-canonical aromatic metabolism. Its high-quality draft genome (4.25 Mbp, 16 contigs, N50 of 887 kb) lacks canonical phenol hydroxylase but encodes alternative oxidation systems, phenylacetyl-CoA pathway, besides, desferrioxamine siderophore, biosurfactants, and phosphate solubilization, key adaptations for oligotrophic caves and biotechnologically interesting activities. Whole-genome comparisons (TYGS/GGDC, OrthoANI and k-mer) suggest potential new species. Lacks acquired antimicrobial resistance genes (ResFinder) and pathogenicity potential (PathogenFinder). *Nocardioides* sp. SF1 emerges as a non-pathogenic candidate for aromatic bioremediation and plant growth promotion in contaminated, nutrient-poor environments, highlighting cave actinobacteria’s unexplored biotechnological potential.

## Introduction

Caves are extreme oligotrophic environments characterized by low nutrient availability, absence of light, and high physicochemical stability, conditions that select highly specialized and often underexplored microbial communities (Turrini et al. 2024). These microorganisms can exhibit remarkable metabolic versatility, synthesizing enzymes and secondary metabolites of biotechnological interest, including degraders of recalcitrant compounds, siderophores, and molecules with potential applications in environmental remediation and plant growth promotion (Marques et al. 2019b; Duncan et al. 2021; Lemes et al. 2021; Zada et al. 2022). Besides, aromatic compounds can serve as sole carbon and energy sources for specialized bacteria, while simultaneously acting as selective agents during enrichment and isolation, thereby favoring the growth of otherwise low-abundance microorganisms (Spini et al. 2018).

The family Nocardiaceae, and specifically the genus *Nocardioides*, comprises actinobacteria known for their tolerance to nutrient-poor conditions and their ability to degrade recalcitrant pollutants (Ma et al. 2023), making them promising targets for the prospection of novel oxidative enzymes and non-canonical catabolic pathways. However, representatives of *Nocardioides* from cave environments remain poorly characterized (Han et al. 2017; Lee et al. 2021) particularly regarding their potential for hydrocarbon transformation, production of biosurfactants and siderophores, and interaction with nutrient cycling. In this context, the isolation and integrated characterization (phenotypic, enzymatic, and genomic) of cave-dwelling strains capable of growing on aromatic compounds represent an opportunity to reveal novel metabolic capabilities and evaluate their biotechnological potential.

In this study, the central objective was to isolate and characterize cave bacteria capable of utilizing toxic aromatic compounds as their sole carbon source, evaluating their tolerance to aromatic compounds, biotechnological potential, and taxonomic position. To this end, bacteria were isolated from three limestone caves in the state of Bahia, Brazil, and their growth profiles on phenol, toluene, and benzene were determined, alongside their extracellular enzymatic activities and phosphate solubilization capacity. Based on these results, a representative *Nocardioides* strain was selected for whole-genome sequencing and phylogenomic and functional analyses, aiming to elucidate the genetic mechanisms associated with aromatic degradation, the production of metabolites of interest, and its potential classification as a novel species within the genus.

## Methodology

### Sample origin

Samples were collected from three limestone caves located in Bahia state, Brazil. These caves are: Buraco da Sopradeira (BS, 12°26′55.97′′S, 44°57′57.32′′W) in São Desidério (Western Bahia), Gruta de Manuel Ioiô (GMI, 12°20′9.96′′S, 41°33′50.04′′ W) in Iraquara (Central Bahia), and Toca da Salamanta (TS, 10°36′16′′ S, 37°53′9′′ W) in Paripiranga (Eastern Bahia). Sampling procedures for BS and GMI were previously described (Marques et al. 2018, 2019a). A rock sample was used from BS, and the L4 sample (approximately 700 m from the entrance) was used from GMI. The TS sample was taken from the superficial sediment layer. All samples were collected in the aphotic zone of the respective caves and authorized by SISBIO/IBAMA/MMA (permit no. 38453).

### Isolation of microorganism

A 5 g sample of cave soil/rock was individually added to an Erlenmeyer containing 50 mL of mineral medium. The medium composition per liter was: 0.25 g KH_2_PO_4_, 2.0 g K_2_HPO_4_, 0.50 g NH_4_Cl, 0.1 g CaCl_2_, 0.5 g of NaCl, 2.0 g of MgCl_2_.6H_2_O (Matos Neto et al. 2023), and supplemented with 10mM of phenol, toluene or benzene as the sole carbon source. After 15 days of incubation at 28°C and 150 RPM, an aliquot of the culture was diluted and plated on agar containing the same mineral medium and the corresponding carbon source. Plates were incubated at 28°C for up to 5 days and resulting morphotypes were isolated in pure culture using the same medium.

### Growth test

The growth of each isolated morphotype was evaluated using the three aromatic compounds (phenol, toluene, and benzene) at the same isolation concentrations (10 mM) to verify their capacity for growth on other aromatic compounds. Growth was monitored up to 120 h with observations taken every 24 h. Growth of each morphotype was also evaluated using the mineral medium supplemented with 0.5% glucose and in BHI (Brain Heart Infusion) medium at 1/10th of the recommended concentration, which was also used for storage in glycerol 20%.

For all growth assays, *Escherichia coli* was included as a negative control, as it is not adapted to grow on phenol, toluene, or benzene as sole carbon sources under the conditions tested.

### Enzymatic assay

Enzymatic assays for protease, amylase, lipase/esterase, and cellulase were conducted on agar plates (2 g/L) using BHI 1/10th supplemented with the respective substrate. The specific substrates utilized were: 1% skim milk for protease activity and 1% tributyrin for lipase/esterase activity. Degradation halos were observed every 24 h up to 120 h. Amylase activity was tested with 1% starch, and the formation of clearance halos was visualized after staining with iodine solution. The phosphate solubilization assay was conducted in NBRIP (National Botanical Research Institute’s Phosphate) medium, and solubilization halo was visualized up to 120 h.

### DNA extraction and 16S rDNA PCR

Microorganisms showing biotechnological potential were selected for DNA extraction. Genomic DNA was extracted after growth in BHI 1/10 using the EasyPure bacterial genomic DNA kit (TransGen Biotech co., China) according to the manufacturer’s instructions. The quality and quantity of the extracted DNA were evaluated spectrophotometrically using a nanodrop ND-1000 (Thermo Fischer Scientific, Minnesota, USA). DNA quality was considered acceptable when the A260/A280 ratio was between 1.8 and 2.0. The 16S rDNA gene was amplified by PCR and sequenced using a standardized methodology (Matos Neto et al. 2023) employing the universal primers F27 and R1525.

### Ortho and meta cleavage assay

For the determination of the catechol cleavage pathway of selected microorganisms, the microorganism was inoculated into 10 mL of liquid mineral medium containing 5 µL of toluene as an inducer. The flasks were incubated under shaking at room temperature for 24 h. Subsequently, the medium was centrifuged at 5,000 g for 5 min to separate the cells from the supernatant. The supernatant was discarded, and the cells were washed three times with 50 mM Tris buffer, pH 8.0. After washing, the cells were resuspended in a total volume of 3 mL containing 1.5 mL of 50 mM Tris buffer, pH 8.0, 1.5 mL of 9 mM catechol, and 0.3 µL of toluene. The tube was shaken and incubated at 28°C. Within 60 min, the formation of a yellow color was monitored, indicating the accumulation of 2-hydroxymuconic semialdehyde (meta-cleavage pathway). If no yellow color developed, the cells were tested after 12 h for the formation of β-ketoadipate using the Rothera reaction (MacLean et al. 2006), which indicates the ortho-cleavage pathway.

### Genome sequence

Based on the identification and the functional assays, one representative isolate was selected for whole-genome sequencing. The genome was sequenced using paired-end sequencing technology on an Illumina Novaseq 6000 platform. Raw sequencing data were assembled using SPAdes 3.15.5 (Prjibelski et al. 2020) Open Reading Frames (ORF) were annotated using RAST (Aziz et al. 2008) and BV-BRC platform (version 3.54.6) (Wattam et al. 2017; Olson et al. 2023). Antimicrobial resistance genes were predicted using ResFinder 4.5.0 (Bortolaia et al. 2020) and potential pathogenicity was predicted with PathogenFinder 1.1 (Cosentino et al. 2013) online tools. Sequence comparisons were performed using the BLASTp and BLASTn algorithms (Altschul et al. 1990). Predicted proteins were searched against the non-redundant protein database, while nucleotide sequences were aligned with nucleotide collection except 16S rDNA that align with rDNA database.

A whole genome-based taxonomic analysis was performed using the Type (Strain) Genome Server (TYGS) platform (Meier-Kolthoff and Göker 2019), which automatically selected ≥15 reference genomes of the genus *Nocardioides* for comparative analysis and generated a SSU tree and a Genome BLAST Distance Phylogeny approach tree (GBPD tree) with FastME 2.1.6.1. Genome-to-genome distance (GGDC) estimation was performed using the Genome-to-Genome Distance Calculator (GGDC) version 3.0 (Meier-Kolthoff et al. 2022) via the DSMZ web service (http://ggdc.dsmz.de). Due to the draft status of the assembly (50 contigs), Formula 2 was prioritized as recommended for incomplete genomes, as it accounts for gene content differences via high-scoring segment pairs with greedy-with-trimming algorithm (Meier-Kolthoff et al. 2022). GGDC values, confidence intervals, and probabilities of DDH ≥70% were calculated against 17 reference *Nocardioides* genomes available at NCBI.

A genome phylogeny tree was constructed with BV-BRC platform’s CodonTree service (Olson et al. 2023) setting of 1000 Single-copy genes (Tables S1 and S2). Sequence alignments were generated with MUSCLE and Biopython (Edgar 2004; Cock et al. 2009), followed by Maximum Likelihood inference using RAxML (Stamatakis 2014) to establish the final phylogenetic relationships among the analyzed genomes and reference sequences.

The proteome comparison analysis was performed in BV-BRC version 3.54.6 using the Proteome Comparison service, which identifies orthologous proteins by bidirectional BLASTP best hits between the reference genome and the selected comparison of representative genomes closely related. A Similar Genome Finder analysis was also conducted in BV-BRC version 3.54.6 to identify the closest related genomes. OrthoANI values (Yoon et al. 2017) were calculated between *Nocardioides* sp. SF1 and the closely related and selected genomes. Heatmaps and hierarchical dendrograms were generated using custom Python scripts with Seaborn v0.13.2 (Waskom 2021) and SciPy v1.13.1 (Virtanen et al. 2020).

Secondary metabolite biosynthetic gene clusters (BGCs) were predicted using antiSMASH (version 7.0.1) (Blin et al. 2023) with default bacterial genome mining parameters. This analysis included the detection of classic BGC types and a comparison against the MiBIG database (Zdouc et al. 2025) to identify similar, previously described clusters. The pipeline provided putative cluster boundaries, predicted product classes, and similarity scores relative to known biosynthetic pathways.

The presence of genes and pathways related to biosurfactant production and hydrocarbon degradation was investigated using BioSurfDB (Oliveira et al. 2015), a curated database and analysis platform dedicated to biosurfactants and biodegradation. Hits corresponding to biosurfactant synthesis enzymes or biosurfactant-associated membrane proteins were annotated as putative biosurfactant-related functions.

## Results

A total of 29 bacterial strains were isolated from the three cave samples (Table 1). Growth assays showed that all isolates were able to use benzene, toluene, and glucose as sole carbon sources, and all strains also grew in BHI medium diluted to 1/10 of the recommended concentration. In contrast, seven strains (TST1, TST2, TST3, TST6, TST7, TSB3, BSB1) did not grow in the presence of phenol as the sole carbon source. Growth on all tested aromatic compounds was detected within 72 h.

**Table 1.** Number of bacterial morphotype isolated per sample using phenol, toluene or benzene as sole carbon source.

| Cave | Number of morphotypes, isolates |  |  |
| --- | --- | --- | --- |
|  | Phenol | Toluene | Benzene |
| Sopradeira | 5, BSF1-BSF5 | 2, BST1-BST2 | 2, BSB1-BSB2 |
| Manuel Ioiô | 6, MIF1-MIF6 | 0 | 2, MIB1-MIB2 |
| Salamanta | 1, TSF1 | 7, TST1-TST7 | 4, TSB1-TSB4 |

Only one isolate (TSB1) grew on NBRIP medium and showed phosphate solubilization capability. None of the isolates exhibited amylase activity under the tested conditions. Lipase/esterase activity was detected in two isolates: TSB4, which produced a visible halo within 24 h, and MIB1, in which halo formation was observed after 72 h. Protease activity was detected in 16 isolates (∼55% of the total), which were grouped according to the time required for halo detection: six isolates (BSF2, BSF4, MIF1, TSB2, MIB1, MIB2) showed halos at 24 h, four (TST6, MIF2, MIF6, BSB1) at 48 h, and the remaining six (BSF1, TST2, MIF4, TSB1, TSB3, TSB4) at 72 h.

Based on growth and enzymatic profiles, selected isolates were subjected to 16S rDNA sequencing. Most isolates were affiliated with the phylum Bacillota, with one Actinomycetota (genus *Nocardioides*) and one Pseudomonadota (genus *Brevundimonas*) also detected (Table 2).

**Table 2.**
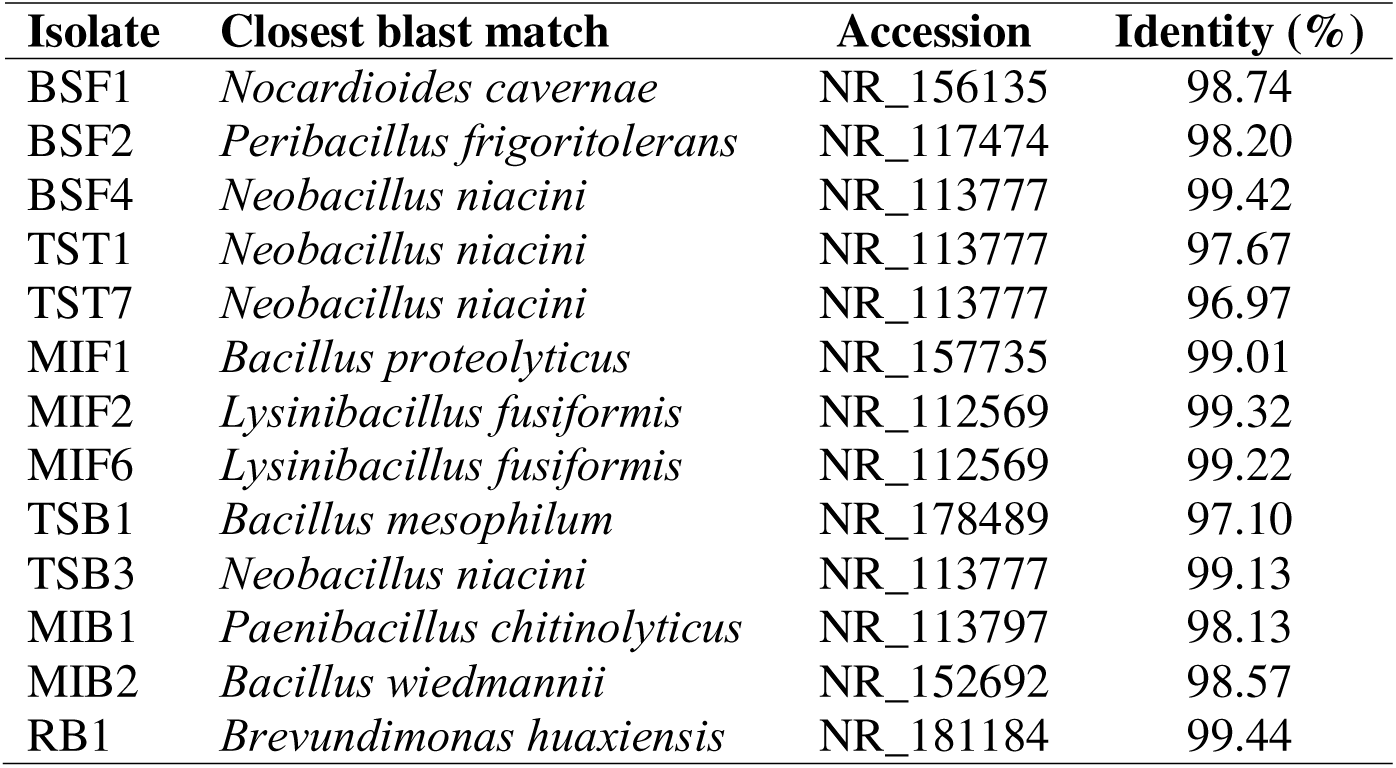
Identification of bacterial isolates based on the nBLAST of 16S rDNA sequencing against ribosomal reference database.

Ortho and meta cleavage assays were inconclusive for two of the tested isolates. Within the first 60 min of incubation with catechol and toluene, both cultures developed an unexpected purple coloration, instead of the characteristic yellow associated with 2-hydroxymuconic semialdehyde formation via the meta-cleavage pathway, and this color change occurred before the addition of the Rothera reagents required for the ortho-cleavage test (Figure 1). This early purple coloration suggests the formation of alternative oxidation products or pigmented intermediates, preventing a reliable assignment of the catechol cleavage pathway for these isolates.

**Figure 1.**
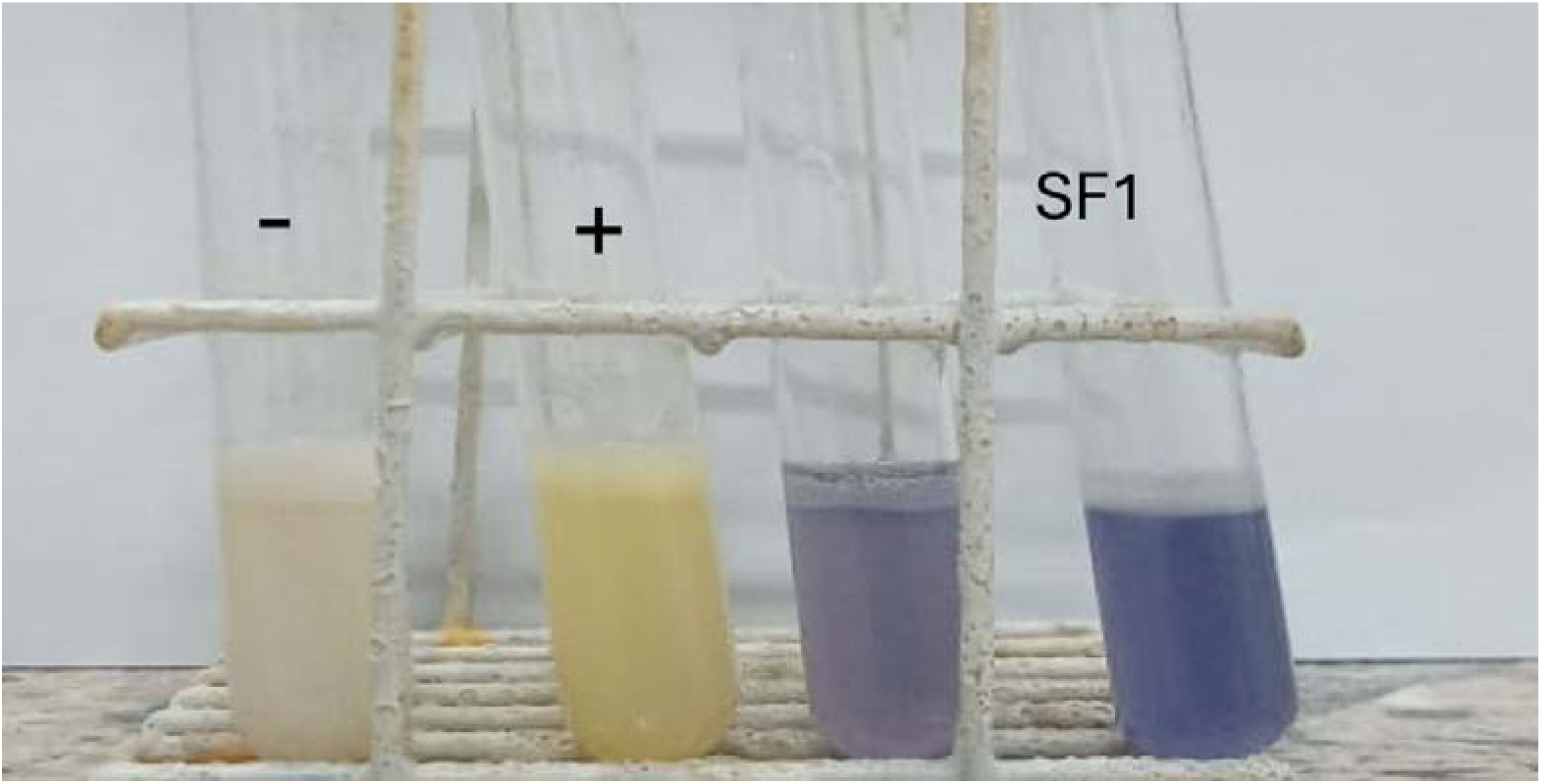
Detection of catechol 2,3-dioxygenase activity (meta-cleavage pathway) in isolate SF1. A positive reaction (+) is indicated by the accumulation of the yellow product, compared to the negative control (–).

We selected the Actinomycetota isolate for whole-genome sequencing. The draft genome of *Nocardioides* sp. SF1 (isolate BSF1) is 4,247,678 bp in size, assembled into 16 contigs, with a GC content of 72.88% and 47 identified RNA genes. The genome showed 0.2% contamination according to CheckM. The N50 of this draft genome is 886,820 bp, and the L50 is 2. A total of 4,107 (BV-BRC) and 4,109 (RAST) coding sequences (CDS), 46 transfer RNA (tRNA) genes, and 2 ribosomal RNA (rRNA) genes were identified. Of these CDS, 965 (24%) were classified into 287 subsystems using RAST. In BV-BRC/PATRIC, the annotation included 1,549 hypothetical proteins (37.7% of the total) and 2,558 proteins with functional assignments. The latter included 917 proteins with Enzyme Commission (EC) numbers, 772 with Gene Ontology (GO) terms, and 685 proteins mapped to KEGG pathways. PATRIC annotation also assigned 3,132 proteins to genus-specific protein families (PLFams) and 3,238 proteins to cross-genus protein families (PGFams). The distribution of the genome annotation is presented in Figure 2.

**Figure 2.**
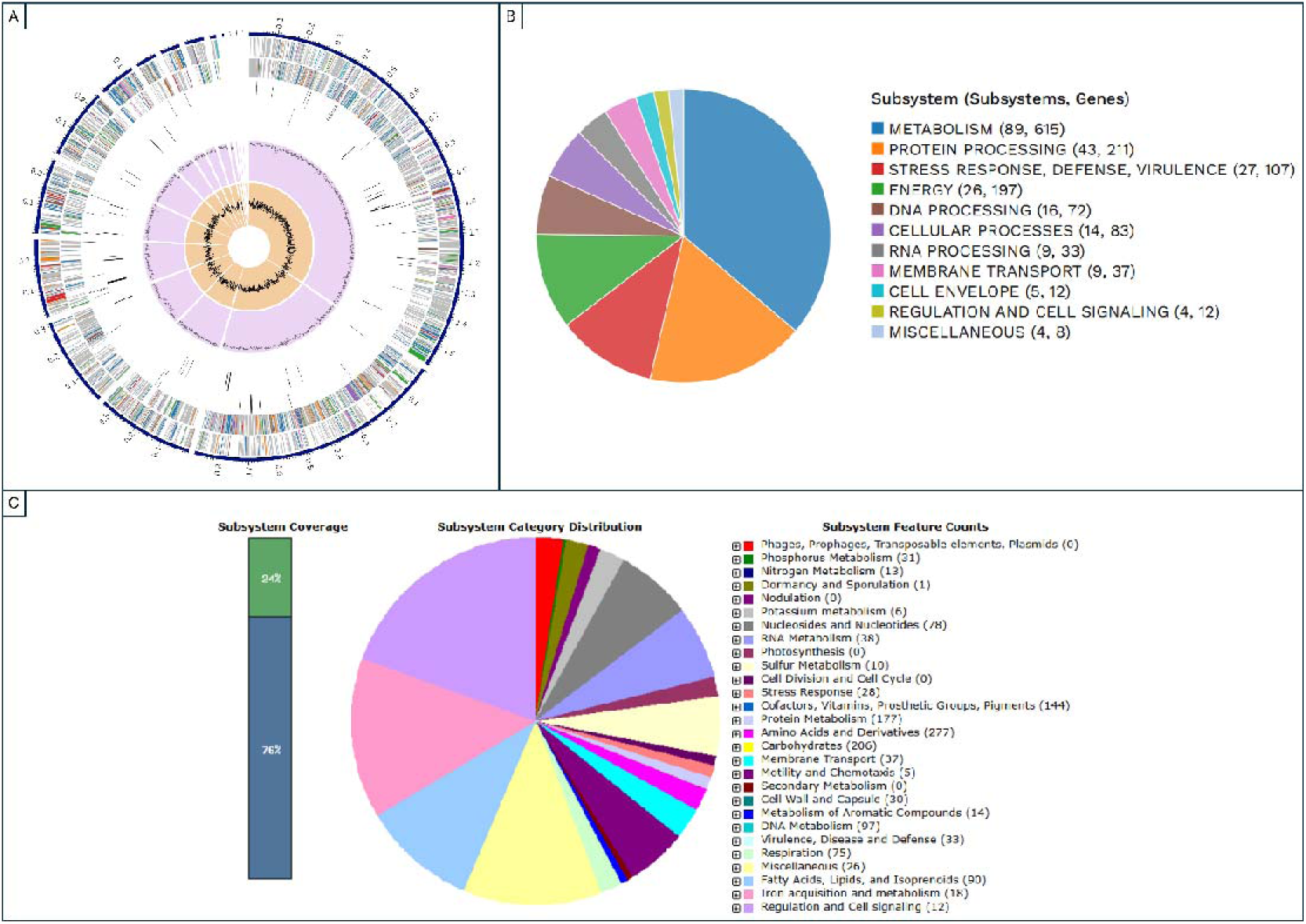
Genome annotation overview of isolate SF1 using BV-BRC and RAST. (A) Circular genomic map and (B) distribution of subsystems generated by BV-BRC; (C) subsystem distribution from RAST annotation. For the circular map (A), rings are displayed from outer to inner as follows: contigs, CDS on the forward strand, CDS on the reverse strand, RNA genes, CDS with homology to known antimicrobial resistance (AMR) genes, CDS with homology to known virulence factors, GC content, and GC skew. CDS colors on the forward and reverse strands correspond to their respective functional subsystems.

Some genes of biotechnological interest were identified in the RAST and BV-BRC annotations, including an alkaline phosphatase and an exopolyphosphatase. In addition, an alkane monooxygenase involved in alkane metabolism was detected, highlighting a potential for bioremediation. In contrast, no canonical phenol-degradation genes, such as phenol hydroxylase or aromatic ring dioxygenases, were identified. Nonetheless, the genome harbors a multicopper polyphenol oxidase, a cyclohexanone monooxygenase, a nitrilotriacetate monooxygenase, genes for the aerobic phenylacetyl-CoA pathway and arsenate reductase (Table S3).

No acquired antimicrobial resistance genes were detected using ResFinder, and no evidence for human pathogenic potential was found with PathogenFinder. In contrast, the BV-BRC analysis identified several genes associated with intrinsic antibiotic targets and resistance-related functions. These included katG, proteins corresponding to known antibiotic targets in susceptible species (Alr, Ddl, dxr, EF-G, EF-Tu, folA/Dfr, folP, gyrA, gyrB, inhA, fabI, iso-tRNA, kasA, rho, rpoB, rpoC, S10p, S12p), target replacement proteins (FabG, HtdX), the gene gidB (conferring resistance via loss-of-function), proteins that modulate cell wall charge (GdpD, PgsA), and the two-component regulators MtrA and MtrB, which can influence the expression of resistance-associated genes.

The antiSMASH analysis predicted three secondary metabolite biosynthetic gene clusters of potential biotechnological interest: a siderophore cluster, a terpene-associated type III polyketide synthase cluster, and a β-lactone cluster. PRISM identified two additional clusters: an NRPS-independent siderophore synthase pathway and a polyketide-associated nonribosomal peptide synthetase (NRPS) pathway. The former corresponds to the siderophore cluster detected by antiSMASH, whereas the latter was not recovered by antiSMASH. The siderophore cluster showed a MiBIG similarity score of 0.8 to the desferrioxamine E biosynthetic gene cluster from *Streptomyces sp*. ID38640 (BGC0001478/MG459167), suggesting that the compound produced by *Nocardioides sp*. SF1 is chemically closely related. The polyketide cluster displayed a MiBIG similarity of 0.68 to the pathways producing 2-methoxy-5-methyl-6-(13-methyltetradecyl)-1,4-benzoquinone and 2-methoxy-5-methyl-6-(13-methyltetradecyl)phenol, which are likely precursors of other natural products. It is important to mention that the BV-BRC also detected polyketide genes (Figure S1).

Analysis with BioSurfDB, using both DNA and protein profiles, revealed the presence of an enzyme related to biosurfactant synthesis and a putisolvin-like integral membrane protein, indicating the potential for biosurfactant production. Although no complete aromatic-compound degradation pathway was detected, individual genes associated with naphthalene and benzoate degradation were present.

To resolve the taxonomic position of strain SF1, several conserved and functional genes were analyzed using BLASTp and BLASTn (Table 3). The 16S rRNA gene sequence showed 98.74% identity with *Nocardioides cavernae* (NR_156135). Analysis of key housekeeping genes provided stronger evidence for the distinctness of the strain. The DNA gyrase subunit B (gyrB), recognized as robust phylogenetic markers in Actinobacteria, shared only 94.22% amino acid identity with their closest matches in the genus *Nocardioides*. Furthermore, the ATP synthase beta chain and glucose-6-phosphate isomerase sequences exhibited identities below 94% with *Nocardioides*. Notably, a multicopper polyphenol oxidase, potentially involved in the degradation of aromatic compounds, showed only 82.25% identity to its closest ortholog. These consistently low identity values across multiple independent markers support the classification of strain SF1 as a novel species within the genus *Nocardioides*.

**Table 3.** Sequence identity (%) of taxonomic markers and selected functional proteins of strain SF1 compared to their closest relatives within the genus *Nocardioides*.

| Gene | BLASTp<br>(identity in %) | Closest Blast<br>match | BLASTn<br>(identity in %) | Closest Blast<br>match |
| --- | --- | --- | --- | --- |
| 16S rDNA | - | - | 98,74% | *Nocardioide*s<br>*cavernae*<br>NR_156135 |
| RecA protein | 99.71 | *Nocardioide*s<br>*okcheonensis*<br>WP_230442827 | 92.65 | *Nocardioide*s<br>*okcheonensis*<br>CP087710 |
| ATP synthase<br>beta chain | 97.73 | *Nocardioide*s<br>*pini*<br>WP_268110515 | 94.66 | *Nocardioide*s sp.<br>CP070961 |
| DNA gyrase<br>subunit B | 94.22 | *Nocardioide*s<br>*palaemonis*<br>WP_211735089 | 90.50 | *Nocardioide*s<br>*okcheonensis*<br>CP087710 |
| Glucose-6-phosphate<br>isomerase | 93.42 | *Nocardioide*s<br>*okcheonensis*<br>WP_230443404 | 81.28 | *Nocardioide*s<br>*okcheonensis*<br>CP087710 |
| Multicopper<br>polyphenol<br>oxidase | 82.25 | *Nocardioide*s sp.<br>WP_374456656 | 91.73 | *Nocardioide*s<br>*okcheonensis*<br>CP087710 |

The TYGS platform automatically selected 16 reference genomes of *Nocardioides* spp. for comparative analysis. Genome-to-genome distance (GGDC) estimation revealed a maximum dDDH (Digital DNA-DNA hybridization) value of 38.9% with *Nocardioides zeicaulis* (GCA_042432345) (Table S4). All other reference genomes yielded dDDH values ranging from 20.3–30.9%, confirming genomic distinctiveness from known species.

Phylogenomic analysis (Figure 3) placed *Nocardioides sp*. SF1 close to *N. zeicaulis*, *N. palaemonis*, *N. okcheonensis* and *Nocardioides sp*. CF479. Initial analyses involving 60 and 16 genomes yielded only 332 and 733 single-copy core genes, respectively. The last one made to enhance phylogenetic resolution and prove more robust topological support. A similar result was found in GBPD tree with *N. zeicaulis*, *N. okcheonensis* and *N. palaemonis* (Figure 4) phylogenetically close to *Nocardioides* sp. SF1.

**Figure 3.**
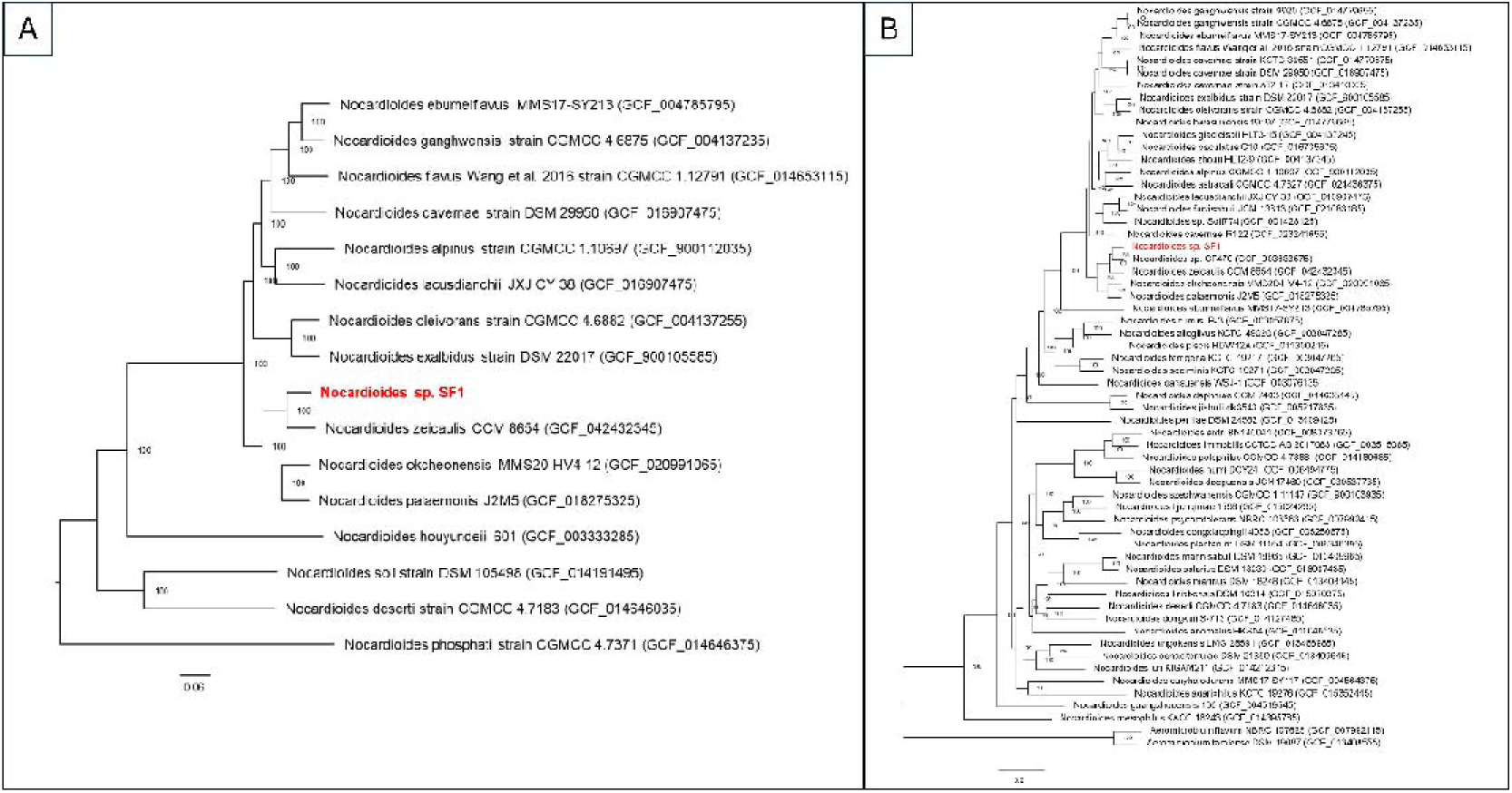
Phylogenomic placement of *Nocardioides sp.* SF1 within the genus using BV-BRC data. The trees were reconstructed using the Maximum Likelihood method based on single-copy core genes. (A) Phylogeny with 16 reference genomes of *Nocardioides* (733 core genes); (B) broad-scale phylogeny including 60 representative genomes (332 core genes). *Aeromicrobium* spp. from Nocardioidaceae family was used as outgroup. Values at the nodes represent bootstrap support. *Nocardioides sp.* SF1 is in bold red font.

**Figure 4.**
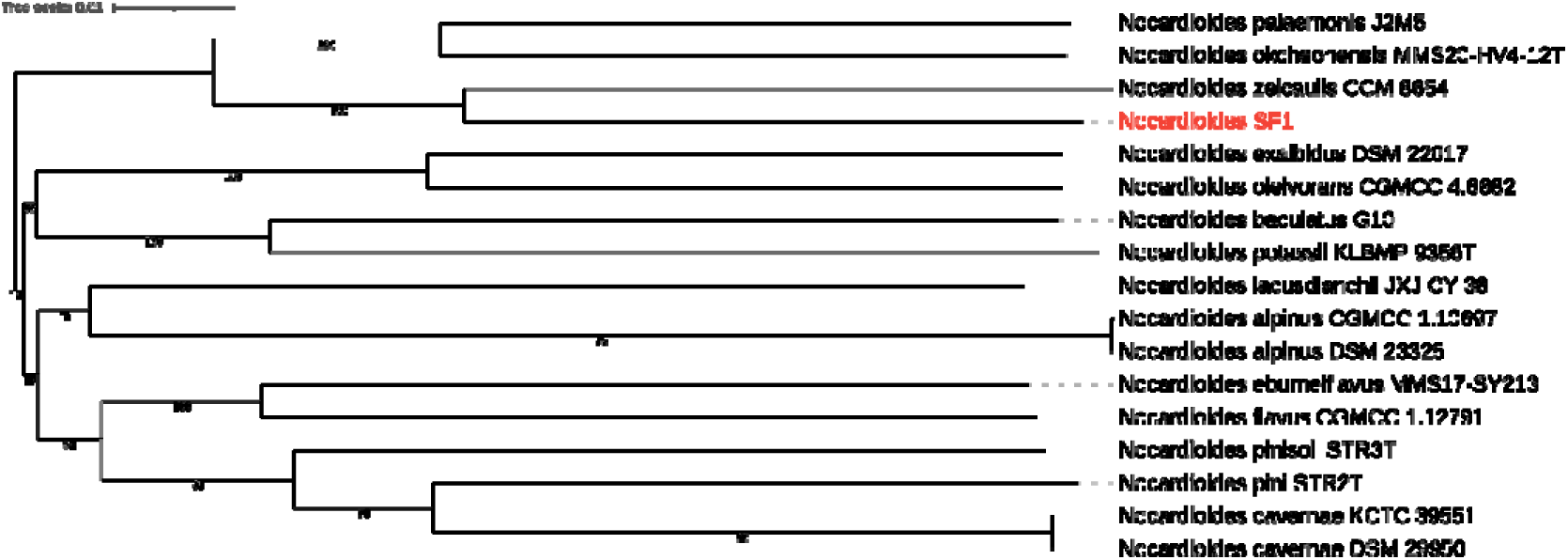
Tree inferred with FastME from Genome BLAST Distance Phylogeny approach (GBPD). The numbers above branches are GBDP pseudo-bootstrap support values > 60 % from 100 replications (average branch support of 85.3 %).

Similar Genome Finder in BV-BRC retrieved *N. okcheonensis, N. ganghwensis* and other *Nocardioides* genomes as the closest relatives, with k-mer distances ranging from 0.075 to 0.123. Consistently, OrthoANI analysis including these genomes showed values between 76.26% and 91.25%, with the highest value obtained between *Nocardioides sp.* SF1 and *Nocardioides* sp. CF479 (isolated from *Populus* sp. roots). The close related described species in OrthoANI is *N. zeicaulis* with a value of 89.88% (Figure 5). These values are well below the commonly accepted species delineation threshold of 95–96%, supporting the placement of strain SF1 within the genus *Nocardioides* while indicating that it likely represents a distinct, previously undescribed species.

**Figure 5.**
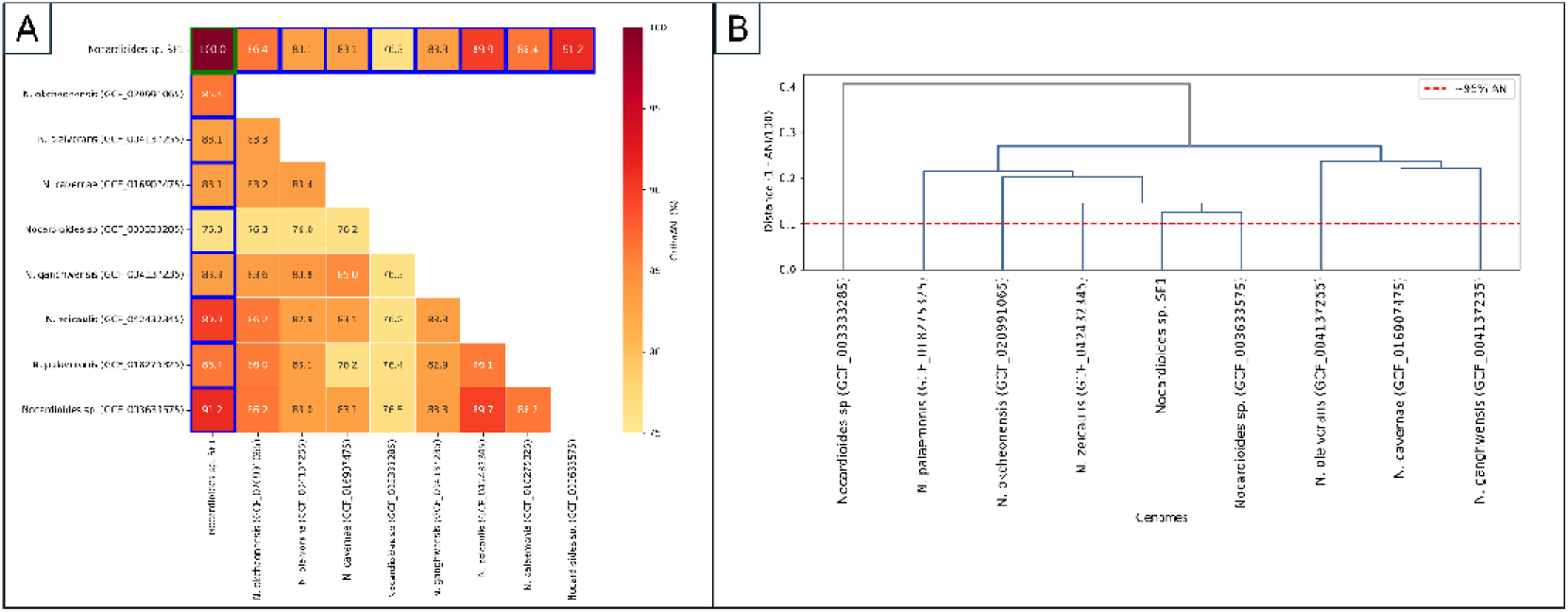
OrthoANI-based genomic relatedness among *Nocardioides* sp. SF1 and related taxa. (A) Heatmap of OrthoANI values in % (red: high similarity; yellow: low similarity); blue lines indicate SF1 genome; (B) Hierarchical clustering dendrogram based on OrthoANI distances, with *Nocardioides* sp. SF1 forming distinct cluster from type strains at 95% threshold

## Discussion

We chose to sequence the *Nocardioides* sp. SF1 isolate because members of this genus are known to tolerate low-nutrient conditions and to degrade recalcitrant pollutants, making them attractive targets for biotechnological applications (Ma et al. 2023). This choice is particularly interesting given that, in a previous study of the same sample (Marques et al. 2019a), the family Nocardiaceae represented only 0.0014% of the community, indicating that enrichment with aromatic compounds as the sole carbon source created a favorable environment for low-abundance microorganisms.

Phenol is metabolically related to toluene and benzene, but its hydroxylated nature confers substantially higher cellular toxicity, requiring robust detoxification and/or efflux mechanisms. The growth failure observed in seven isolates suggests that the morphotypes selected directly on toluene and benzene are more susceptible to phenol toxicity than those initially isolated on phenol, supporting the idea that phenol acted as a strong selective agent during the isolation phase. A concentration range of 1–10 mM for phenol and BTEX is widely used in degradation assays (Santos and Linardi 2004), as it is low enough to allow growth of adapted strains but high enough to inhibit non-adapted lineages. The ability of *Nocardioides* sp. SF1 (and the other 21 isolates) to grow on phenol at this concentration underscores their high tolerance and bioremediation potential.

It is notable that the genome of *Nocardioides* sp. SF1 does not encode a canonical phenol hydroxylase, the main entry enzyme in the classical aerobic phenol degradation pathway. This suggests that SF1’s survival and degradation capacity may rely on alternative genomic systems, such as the multicopper polyphenol oxidase (PPO), which can oxidize phenolic substrates, together with an expanded repertoire of efflux transporters that likely contributes to the high tolerance required to withstand phenol and related aromatic compounds.

Although no canonical phenol hydroxylase genes were detected, the genome encodes several flavin[dependent monooxygenases and a multicopper polyphenol oxidase, which are known to oxidize phenolic substrates and could function as alternative entry points into aromatic degradation pathways in strain SF1. In addition, the presence of the aerobic phenylacetyl[CoA pathway and other ring[cleavage genes suggests that phenol-derived intermediates might be funneled into non[classical routes, a hypothesis that warrants future validation.

The draft genome of *Nocardioides* sp. SF1 (4.24 Mbp, GC content of 72.88%) exhibits good assembly quality, as evidenced by its contiguity metrics. The N50 of 886,820 bp and L50 of 2 indicate that the SPAdes assembly produced long, highly continuous contigs (only 16 contigs in total). This genomic robustness is further supported by the 0.2% contamination estimated by CheckM, ensuring that the gene set used in downstream analyses is representative of the complete genome of the isolate.

GGDC analysis conducted via TYGS confirmed this taxonomic distinctiveness with a dDDH value below the 70% species threshold recommended for draft assemblies (Meier-Kolthoff et al. 2013). Interestingly, the closest genome was different from k-mer analysis, though all values remain below the species thresholds. Together, GGDC distances, OrthoANI and k[mer values between *Nocardioides* sp. SF1 and the closest related genomes, combined with the high quality and completeness of its genome, strongly indicate that this strain represents a new undescribed *Nocardioides* species. Furthermore, the discrepancies between high protein conservation and low nucleotide identity on BLAST of selected genes provide additional evidence that strain SF1 represents a novel species within the *Nocardioides* genus and related to *N. zeicaulis, N. okcheonensis* and *N. palaemonis*.

The presence of phenylacetyl[CoA degradation operons and noncanonical monooxygenases in such a nearly complete genome suggests that these pathways are genuinely encoded and likely central to the ecology of the strain, rather than assembly artifacts. Phosphatases encoded in the genome, together with the phosphate-solubilizing phenotype observed for TSB1, suggest that *Nocardioides* sp. SF1 and related isolates may contribute to phosphate mobilization in cave soil and could act as plant growth-promoting bacteria, a role already reported for this species complex (Soumare et al. 2021). In addition, the desferrioxamine E–like siderophore biosynthetic cluster identified in SF1 is associated with molecules that have recognized biotechnological value, including as plant growth promoters and metal chelators (Mahajan et al. 2021). Finally, the presence of a cyclohexanone monooxygenase, a well-known Baeyer–Villiger monooxygenase (Torres Pazmiño et al. 2010), further expands the oxidative repertoire of SF1 toward hydrophobic substrates and supports its potential application in biocatalysis and bioremediation.

Notably, ResFinder did not detect any acquired antimicrobial resistance genes in *Nocardioides* sp. SF1, whereas BV-BRC annotation identified multiple loci corresponding to known antibiotic targets and resistance-associated functions. This apparent discrepancy reflects the different scopes of the tools, ResFinder is designed to detect horizontally acquired or clinically validated resistance determinants, while the BV-BRC AMR annotation also flags housekeeping genes that can confer resistance only when specific mutations are present. In SF1, these genes show no known resistance-associated variants, suggesting that they represent intrinsic antibiotic targets rather than acquired AMR determinants, and are therefore unlikely to pose a significant risk for environmental or biotechnological applications. This interpretation is further supported by PathogenFinder, which did not predict any potential for human pathogenicity in *Nocardioides* sp. SF1.

From an ecological and biogeochemical perspective, these findings highlight the importance of the rare biosphere as an enzymatic reservoir in cave ecosystems. Under natural oligotrophic conditions, these low-abundance taxa may remain stable or latent. However, when specific carbon inputs, such as natural or anthropogenic aromatic hydrocarbons, become available, these specialized microorganisms rapidly proliferate, actively driving the subterranean carbon cycle through alternative oxidative and non-canonical pathways, such as the phenylacetyl-CoA route identified here. Furthermore, the functional genome of *Nocardioides* sp. SF1 demonstrates that this biogeochemical role extends beyond carbon turnover. The presence of genes for alkaline phosphatase, exopolyphosphatase, and siderophore biosynthesis suggests that once these rare taxa are present, they can influence the phosphorus and iron cycles within the cave matrix. By mobilizing inorganic phosphate from limestone rocks and chelating metals, these bacteria act as a biogeochemical node, transforming insoluble minerals into bioavailable forms that sustain the local subterranean food web.

## Conclusion

*Nocardioides* sp. SF1 is a non-pathogenic cave actinobacterium that combines high tolerance to aromatic compounds with genomic features supporting bioremediation and plant growth promotion, including alternative phenol-oxidizing systems, siderophore and biosurfactant biosynthesis, and phosphate mobilization. Genomic distances (GGDC Formula 2: 38.9% maximum dDDH; OrthoANI <95%) to the closest described *Nocardioides* taxa, combined with polyphasic evidence (16S rRNA, rpoB, gyrB; and phylogeny), suggest SF1 represents a novel species with relevant biotechnological potential in contaminated and nutrient-poor environments.

## Acknowledgment

We would like to thank the National Council for Scientific and Technological Development (CNPq), Grant Number 435702/2018–1, for funding the research

