## Supplementary material for "Biotechnological potential of aromatic compounds–utilizing bacteria from Brazilian caves, including a novel cave *Nocardioides sp*. SF1": Table S1

**Table S1 – Single-copy genes used to phylogenomic of 16 *Nocardioides* species**

| **PGFam** | **Align. Score** | **Align. Length** | **Num Seqs** | **Mean Sqr Freq** | **Prop Gaps** | **Used In Analysis** | **Product** |
| --- | --- | --- | --- | --- | --- | --- | --- |
| PGF_00008115 | 34.63 | 1551 | 16 | 0.879 | 0.021 | True | Glutamate synthase [NADPH] large chain (EC 1.4.1.13) |
| PGF_02797402 | 31.33 | 1234 | 16 | 0.892 | 0.040 | True | DNA-directed RNA polymerase beta subunit (EC 2.7.7.6) |
| PGF_00423351 | 30.53 | 1322 | 16 | 0.840 | 0.042 | True | Dihydrolipoamide succinyltransferase component (E2) of 2-oxoglutarate dehydrogenase complex (EC 2.3.1.61) / 2-oxoglutarate dehydrogenase E1 component (EC 1.2.4.2) @ 2-oxoglutarate decarboxylase (EC 4.1.1.71) @ 2-hydroxy-3-oxoadipate synthase (EC 2.2.1.5) |
| PGF_10406880 | 29.81 | 1197 | 16 | 0.862 | 0.009 | True | DNA polymerase III alpha subunit (EC 2.7.7.7) |
| PGF_00006434 | 29.53 | 1359 | 16 | 0.801 | 0.046 | True | FtsK/SpoIIIE family protein, putative EssC/YukB component of Type VII secretion system |
| PGF_00416410 | 29.52 | 1135 | 16 | 0.876 | 0.021 | True | Carbamoyl-phosphate synthase large chain (EC 6.3.5.5) |
| PGF_08675943 | 29.39 | 1232 | 16 | 0.837 | 0.031 | True | Transcription-repair coupling factor |
| PGF_09950730 | 29.21 | 1219 | 16 | 0.837 | 0.027 | True | Chromosome partition protein smc |
| PGF_08152874 | 28.32 | 1442 | 16 | 0.746 | 0.106 | True | ATP-dependent helicase HrpA |
| PGF_08854233 | 27.82 | 995 | 16 | 0.882 | 0.033 | True | Ribonucleotide reductase of class II (coenzyme B12-dependent) (EC 1.17.4.1) |
| PGF_05171623 | 27.62 | 1138 | 16 | 0.819 | 0.059 | True | Isoleucyl-tRNA synthetase (EC 6.1.1.5) |
| PGF_03999196 | 27.56 | 871 | 16 | 0.934 | 0.013 | True | ATP-dependent Clp protease, ATP-binding subunit ClpC |
| PGF_10049811 | 26.46 | 970 | 16 | 0.850 | 0.026 | True | Protein translocase subunit SecA |
| PGF_00008773 | 26.32 | 997 | 16 | 0.834 | 0.038 | True | Glycine dehydrogenase [decarboxylating] (glycine cleavage system P protein) (EC 1.4.4.2) |
| PGF_10300474 | 26.23 | 947 | 16 | 0.852 | 0.042 | True | DNA topoisomerase I (EC 5.99.1.2) |
| PGF_00529632 | 26.01 | 886 | 16 | 0.874 | 0.026 | True | Chaperone protein ClpB (ATP-dependent unfoldase) |
| PGF_04243787 | 26.00 | 991 | 16 | 0.826 | 0.070 | True | Aconitate hydratase (EC 4.2.1.3) |
| PGF_02409931 | 25.78 | 1130 | 16 | 0.767 | 0.045 | True | ATP-dependent DNA helicase SCO5184 |
| PGF_00426726 | 25.77 | 961 | 16 | 0.831 | 0.027 | True | FIG005666: putative helicase |
| PGF_07830674 | 25.65 | 922 | 16 | 0.845 | 0.024 | True | Alanyl-tRNA synthetase (EC 6.1.1.7) |
| PGF_01053024 | 25.64 | 1005 | 16 | 0.809 | 0.011 | True | Glutamine synthetase adenylyl-L-tyrosine phosphorylase (EC 2.7.7.89) / Glutamate-ammonia-ligase adenylyltransferase (EC 2.7.7.42) |
| PGF_10525419 | 25.59 | 1043 | 16 | 0.792 | 0.111 | True | Translation initiation factor 2 |
| PGF_07517180 | 25.52 | 1142 | 16 | 0.755 | 0.051 | True | ATP-dependent DNA helicase SCO5183 |
| PGF_05500127 | 25.50 | 898 | 16 | 0.851 | 0.025 | True | Valyl-tRNA synthetase (EC 6.1.1.9) |
| PGF_03272313 | 25.27 | 974 | 16 | 0.810 | 0.071 | True | DNA gyrase subunit A (EC 5.99.1.3) |
| PGF_03104485 | 24.74 | 757 | 16 | 0.899 | 0.017 | True | Polyribonucleotide nucleotidyltransferase (EC 2.7.7.8) |
| PGF_00041217 | 24.61 | 890 | 16 | 0.825 | 0.027 | True | Putative helicase |
| PGF_00010349 | 24.60 | 780 | 16 | 0.881 | 0.024 | True | Guanosine-3',5'-bis(diphosphate) 3'-pyrophosphohydrolase (EC 3.1.7.2) / GTP pyrophosphokinase (EC 2.7.6.5), (p)ppGpp synthetase II |
| PGF_00950554 | 24.35 | 725 | 16 | 0.904 | 0.022 | True | Excinuclease ABC subunit B |
| PGF_00033961 | 24.31 | 793 | 16 | 0.863 | 0.030 | True | Phosphoribosylformylglycinamidine synthase, synthetase subunit (EC 6.3.5.3) |
| PGF_00060409 | 24.19 | 705 | 16 | 0.911 | 0.001 | True | Translation elongation factor G |
| PGF_00021022 | 24.14 | 748 | 16 | 0.883 | 0.033 | True | Methylmalonyl-CoA mutase large subunit, MutB (EC 5.4.99.2) |
| PGF_10471233 | 23.96 | 855 | 16 | 0.820 | 0.064 | True | ATP-dependent DNA helicase UvrD/PcrA (EC 3.6.4.12) |
| PGF_08155727 | 23.96 | 844 | 16 | 0.825 | 0.043 | True | DNA translocase FtsK |
| PGF_06005188 | 23.62 | 702 | 16 | 0.892 | 0.018 | True | DNA topoisomerase IV subunit B (EC 5.99.1.3) |
| PGF_00425021 | 23.39 | 1183 | 16 | 0.680 | 0.054 | True | Exodeoxyribonuclease V gamma chain (EC 3.1.11.5) |
| PGF_00015703 | 23.37 | 746 | 16 | 0.856 | 0.014 | True | Isocitrate dehydrogenase [NADP] (EC 1.1.1.42); Monomeric isocitrate dehydrogenase [NADP] (EC 1.1.1.42) |
| PGF_00041234 | 23.28 | 800 | 16 | 0.823 | 0.025 | True | Putative helicase |
| PGF_06812369 | 23.27 | 873 | 16 | 0.787 | 0.046 | True | Leucyl-tRNA synthetase (EC 6.1.1.4) |
| PGF_01058419 | 23.26 | 1210 | 16 | 0.669 | 0.077 | True | Exodeoxyribonuclease V beta chain (EC 3.1.11.5) |
| PGF_10357457 | 23.08 | 639 | 16 | 0.913 | 0.027 | True | Chaperone protein DnaK |
| PGF_00007041 | 23.05 | 634 | 16 | 0.915 | 0.007 | True | GTP-binding protein TypA/BipA |
| PGF_00033095 | 23.04 | 845 | 16 | 0.793 | 0.019 | True | Phenylalanyl-tRNA synthetase beta chain (EC 6.1.1.20) |
| PGF_06703483 | 22.97 | 752 | 16 | 0.838 | 0.034 | True | DNA gyrase subunit B (EC 5.99.1.3) |
| PGF_05195470 | 22.88 | 735 | 16 | 0.844 | 0.014 | True | Polyphosphate kinase (EC 2.7.4.1) |
| PGF_00045941 | 22.70 | 845 | 16 | 0.781 | 0.080 | True | Pyrophosphate-energized proton pump (EC 3.6.1.1) |
| PGF_00424921 | 22.61 | 695 | 16 | 0.858 | 0.035 | True | Ethylmalonyl-CoA mutase, methylsuccinyl-CoA-forming |
| PGF_00060414 | 22.52 | 665 | 16 | 0.873 | 0.054 | True | Translation elongation factor LepA |
| PGF_10429292 | 22.48 | 1050 | 16 | 0.694 | 0.131 | True | Helicase, SNF2/RAD54 family |
| PGF_03752158 | 22.45 | 718 | 16 | 0.838 | 0.081 | True | FIG092679: Fe-S oxidoreductase |
| PGF_10467689 | 22.34 | 1032 | 16 | 0.695 | 0.066 | True | Glycosyl transferase, family 2 |
| PGF_01175575 | 22.30 | 672 | 16 | 0.860 | 0.015 | True | Threonyl-tRNA synthetase (EC 6.1.1.3) |
| PGF_00426115 | 22.10 | 643 | 16 | 0.871 | 0.014 | True | 1-deoxy-D-xylulose 5-phosphate synthase (EC 2.2.1.7) |
| PGF_06525160 | 22.07 | 706 | 16 | 0.830 | 0.035 | True | NAD synthetase (EC 6.3.1.5) / Glutamine amidotransferase chain of NAD synthetase |
| PGF_00424971 | 22.03 | 709 | 16 | 0.827 | 0.063 | True | Excinuclease ABC subunit C |
| PGF_02059020 | 21.96 | 560 | 16 | 0.928 | 0.000 | True | Energy-dependent translational throttle protein EttA |
| PGF_01136362 | 21.86 | 729 | 16 | 0.810 | 0.031 | True | Transketolase (EC 2.2.1.1) |
| PGF_00071558 | 21.73 | 604 | 16 | 0.884 | 0.039 | True | Bacterial proteasome-activating AAA-ATPase (PAN) |
| PGF_04569524 | 21.72 | 548 | 16 | 0.928 | 0.004 | True | ATP synthase alpha chain (EC 3.6.3.14) |
| PGF_04674543 | 21.57 | 549 | 16 | 0.921 | 0.002 | True | DNA repair helicase |
| PGF_02226715 | 21.54 | 763 | 16 | 0.780 | 0.032 | True | ATP-dependent DNA helicase RecG (EC 3.6.4.12) |
| PGF_00423472 | 21.52 | 699 | 16 | 0.814 | 0.044 | True | DinG family ATP-dependent helicase YoaA |
| PGF_00416129 | 21.40 | 573 | 16 | 0.894 | 0.015 | True | CTP synthase (EC 6.3.4.2) |
| PGF_00057270 | 21.36 | 1097 | 16 | 0.645 | 0.225 | True | DNA topoisomerase IV subunit A (EC 5.99.1.3) |
| PGF_00016824 | 21.21 | 705 | 16 | 0.799 | 0.030 | True | ATP-dependent DNA helicase UvrD/PcrA, actinomycete paralog |
| PGF_00025090 | 21.19 | 605 | 16 | 0.861 | 0.032 | True | Acetolactate synthase large subunit (EC 2.2.1.6) |
| PGF_00407244 | 21.18 | 772 | 16 | 0.762 | 0.083 | True | putative helicase regulator |
| PGF_10376398 | 21.17 | 739 | 16 | 0.779 | 0.040 | True | Peptidoglycan D,D-transpeptidase MrdA (EC 3.4.16.4) |
| PGF_05562713 | 20.99 | 755 | 16 | 0.764 | 0.020 | True | [Protein-PII] uridylyltransferase (EC 2.7.7.59) / [Protein-PII]-UMP uridylyl-removing enzyme |
| PGF_10487171 | 20.93 | 618 | 16 | 0.842 | 0.042 | True | SdrA |
| PGF_02552445 | 20.88 | 532 | 16 | 0.905 | 0.000 | True | Bis-ABC ATPase SCO1840 |
| PGF_10321811 | 20.83 | 551 | 16 | 0.887 | 0.019 | True | Acetyl-coenzyme A carboxyl transferase alpha chain (EC 6.4.1.2) / Acetyl-coenzyme A carboxyl transferase beta chain (EC 6.4.1.2); Propionyl-CoA carboxylase beta chain (EC 6.4.1.3) |
| PGF_09939762 | 20.83 | 524 | 16 | 0.910 | 0.054 | True | SSU ribosomal protein S1p |
| PGF_12694106 | 20.81 | 599 | 16 | 0.850 | 0.035 | True | 2-isopropylmalate synthase (EC 2.3.3.13) |
| PGF_00006935 | 20.77 | 530 | 16 | 0.902 | 0.009 | True | GMP synthase [glutamine-hydrolyzing], amidotransferase subunit (EC 6.3.5.2) / GMP synthase [glutamine-hydrolyzing], ATP pyrophosphatase subunit (EC 6.3.5.2) |
| PGF_00006449 | 20.76 | 632 | 16 | 0.826 | 0.088 | True | Fumarate hydratase class I (EC 4.2.1.2) |
| PGF_03144461 | 20.69 | 700 | 16 | 0.782 | 0.038 | True | Bifunctional beta-1,5/1,6-galactofuranosyltransferase GlfT2 in cell wall galactan polymerization (EC 2.4.1.288) |
| PGF_00004392 | 20.65 | 575 | 16 | 0.861 | 0.009 | True | Ferredoxin--sulfite reductase, actinobacterial type (EC 1.8.7.1) |
| PGF_00061802 | 20.65 | 640 | 16 | 0.816 | 0.089 | True | Trehalose synthase (EC 5.4.99.16) @ Alpha-amylase (EC 3.2.1.1) |
| PGF_00037588 | 20.65 | 598 | 16 | 0.844 | 0.010 | True | Prolyl-tRNA synthetase (EC 6.1.1.15), bacterial type |
| PGF_00015514 | 20.64 | 481 | 16 | 0.941 | 0.015 | True | Iron-sulfur cluster assembly protein SufB |
| PGF_07904640 | 20.57 | 512 | 16 | 0.909 | 0.009 | True | XRE-family DNA-binding domain / UDP-N-acetylglucosamine 1-carboxyvinyltransferase (EC 2.5.1.7) |
| PGF_06122750 | 20.55 | 636 | 16 | 0.815 | 0.056 | True | DNA polymerase III epsilon subunit-related protein MSMEG4261 |
| PGF_00066286 | 20.55 | 567 | 16 | 0.863 | 0.028 | True | Uroporphyrinogen-III methyltransferase (EC 2.1.1.107) / Uroporphyrinogen-III synthase (EC 4.2.1.75) |
| PGF_05195027 | 20.52 | 486 | 16 | 0.931 | 0.003 | True | ATP synthase beta chain (EC 3.6.3.14) |
| PGF_00052238 | 20.36 | 543 | 16 | 0.874 | 0.036 | True | Signal recognition particle protein Ffh |
| PGF_04333086 | 20.32 | 815 | 16 | 0.712 | 0.117 | True | DNA ligase (NAD(+)) (EC 6.5.1.2) |
| PGF_00038970 | 20.29 | 453 | 16 | 0.953 | 0.000 | True | Pup ligase PafA, possible component of postulated heterodimer PafA-PafA' |
| PGF_00067382 | 20.19 | 578 | 16 | 0.840 | 0.075 | True | (R)-citramalate synthase (EC 2.3.1.182) |
| PGF_03609651 | 20.19 | 615 | 16 | 0.814 | 0.026 | True | Methionyl-tRNA synthetase (EC 6.1.1.10) |
| PGF_00039287 | 20.17 | 767 | 16 | 0.728 | 0.039 | True | Putative DNA-binding protein |
| PGF_03029859 | 20.09 | 638 | 16 | 0.795 | 0.047 | True | Bis-ABC ATPase Uup |
| PGF_00017545 | 20.06 | 612 | 16 | 0.811 | 0.022 | True | Long-chain-fatty-acid--CoA ligase (EC 6.2.1.3) @ Long-chain fatty-acid-CoA ligase (EC 6.2.1.3), Mycobacterial subgroup FadD15 |
| PGF_00037269 | 19.96 | 693 | 16 | 0.758 | 0.080 | True | 2-Amino-2-deoxy-isochorismate synthase (EC 4.1.3.-) |
| PGF_09986522 | 19.96 | 756 | 16 | 0.726 | 0.122 | True | Putative glucanase glgE (EC 3.2.1.-) |
| PGF_00038969 | 19.89 | 506 | 16 | 0.884 | 0.002 | True | Pup ligase PafA' paralog, possible component of postulated heterodimer PafA-PafA' |
| PGF_00416702 | 19.88 | 777 | 16 | 0.713 | 0.030 | True | Catalase-peroxidase KatG (EC 1.11.1.21) |
| PGF_00067554 | 19.81 | 513 | 16 | 0.874 | 0.020 | True | Aspartyl-tRNA(Asn) amidotransferase subunit B (EC 6.3.5.6) @ Glutamyl-tRNA(Gln) amidotransferase subunit B (EC 6.3.5.7) |
| PGF_00419420 | 19.61 | 483 | 16 | 0.892 | 0.025 | True | 3-isopropylmalate dehydratase large subunit (EC 4.2.1.33) |
| PGF_00024687 | 19.45 | 457 | 16 | 0.910 | 0.021 | True | NADH-ubiquinone oxidoreductase chain D (EC 1.6.5.3) |
| PGF_00009966 | 19.42 | 472 | 16 | 0.894 | 0.011 | True | Glycyl-tRNA synthetase (EC 6.1.1.14) |
| PGF_00421796 | 19.42 | 603 | 16 | 0.791 | 0.022 | True | DNA repair protein RecN |
| PGF_02862285 | 19.40 | 484 | 16 | 0.882 | 0.012 | True | Argininosuccinate synthase (EC 6.3.4.5) |
| PGF_00016338 | 19.35 | 434 | 16 | 0.929 | 0.017 | True | ATP-dependent Clp protease ATP-binding subunit ClpX |
| PGF_10440725 | 19.33 | 501 | 16 | 0.864 | 0.024 | True | Glutamate synthase [NADPH] small chain (EC 1.4.1.13) |
| PGF_02029783 | 19.30 | 541 | 16 | 0.830 | 0.037 | True | GTP-binding protein Obg |
| PGF_00060428 | 19.29 | 397 | 16 | 0.968 | 0.000 | True | Translation elongation factor Tu |
| PGF_00013509 | 19.27 | 551 | 16 | 0.821 | 0.044 | True | IMP cyclohydrolase (EC 3.5.4.10) / Phosphoribosylaminoimidazolecarboxamide formyltransferase (EC 2.1.2.3) |
| PGF_00419707 | 19.25 | 446 | 16 | 0.911 | 0.000 | True | Crotonyl-CoA carboxylase/reductase, ethylmalonyl-CoA producing |
| PGF_00007028 | 19.24 | 545 | 16 | 0.824 | 0.059 | True | Ribosome LSU-associated GTP-binding protein HflX |
| PGF_06522349 | 19.22 | 530 | 16 | 0.835 | 0.044 | True | Aspartyl-tRNA(Asn) amidotransferase subunit A (EC 6.3.5.6) @ Glutamyl-tRNA(Gln) amidotransferase subunit A (EC 6.3.5.7) |
| PGF_06672507 | 19.16 | 782 | 16 | 0.685 | 0.086 | True | Probable serine/threonine-protein kinase pknL (EC 2.7.11.1) |
| PGF_00050995 | 19.11 | 553 | 16 | 0.813 | 0.067 | True | Amidophosphoribosyltransferase (EC 2.4.2.14) |
| PGF_00001493 | 19.09 | 631 | 16 | 0.760 | 0.061 | True | P-loop GTPase domain-containing protein Sros_8871 |
| PGF_00007024 | 19.06 | 460 | 16 | 0.889 | 0.012 | True | GTP-binding protein EngA |
| PGF_12879692 | 19.06 | 708 | 16 | 0.716 | 0.055 | True | Helicase PriA essential for oriC/DnaA-independent DNA replication |
| PGF_08562657 | 19.06 | 526 | 16 | 0.831 | 0.046 | True | tRNA-i(6)A37 methylthiotransferase (EC 2.8.4.3) |
| PGF_09658708 | 19.03 | 468 | 16 | 0.880 | 0.004 | True | Dihydrolipoamide dehydrogenase (EC 1.8.1.4) |
| PGF_07540743 | 19.02 | 565 | 16 | 0.800 | 0.031 | True | Thiamin ABC transporter, transmembrane component |
| PGF_07459509 | 18.99 | 622 | 16 | 0.762 | 0.094 | True | L-aspartate oxidase (EC 1.4.3.16) |
| PGF_04704185 | 18.98 | 466 | 16 | 0.879 | 0.008 | True | NAD(P)H quinone reductase LpdA (EC 1.6.5.2) |
| PGF_06887246 | 18.97 | 810 | 16 | 0.667 | 0.061 | True | DNA internalization-related competence protein ComEC/Rec2 |
| PGF_02064356 | 18.85 | 490 | 16 | 0.852 | 0.016 | True | Adenosylhomocysteinase (EC 3.3.1.1) |
| PGF_00045065 | 18.85 | 514 | 16 | 0.831 | 0.019 | True | Putative transmembrane transport protein |
| PGF_08398205 | 18.83 | 505 | 16 | 0.838 | 0.039 | True | CCA tRNA nucleotidyltransferase (EC 2.7.7.72) |
| PGF_00001487 | 18.74 | 556 | 16 | 0.795 | 0.009 | True | P-loop GTPase domain-containing protein Sros_8870 |
| PGF_02564336 | 18.71 | 626 | 16 | 0.748 | 0.054 | True | Multidrug efflux pump P55 |
| PGF_00018085 | 18.62 | 629 | 16 | 0.742 | 0.092 | True | Lysyl-tRNA synthetase (class I) (EC 6.1.1.6) |
| PGF_09031534 | 18.57 | 799 | 16 | 0.657 | 0.187 | True | 1,4-alpha-glucan (glycogen) branching enzyme, GH-13-type (EC 2.4.1.18) |
| PGF_02620298 | 18.53 | 495 | 16 | 0.833 | 0.030 | True | Argininosuccinate lyase (EC 4.3.2.1) |
| PGF_04425336 | 18.47 | 458 | 16 | 0.863 | 0.043 | True | NADH-ubiquinone oxidoreductase chain F (EC 1.6.5.3) |
| PGF_00051220 | 18.45 | 517 | 16 | 0.811 | 0.047 | True | Serine phosphatase RsbU, regulator of sigma subunit |
| PGF_02452671 | 18.44 | 490 | 16 | 0.833 | 0.044 | True | Cell division trigger factor (EC 5.2.1.8) |
| PGF_02516909 | 18.42 | 425 | 16 | 0.894 | 0.000 | True | Enolase (EC 4.2.1.11) |
| PGF_06051594 | 18.41 | 474 | 16 | 0.846 | 0.049 | True | Predicted ATPase related to phosphate starvation-inducible protein PhoH |
| PGF_05581732 | 18.41 | 440 | 16 | 0.878 | 0.019 | True | Protein translocase subunit SecY |
| PGF_01137124 | 18.34 | 454 | 16 | 0.861 | 0.012 | True | Phosphoglucosamine mutase (EC 5.4.2.10) |
| PGF_00054959 | 18.33 | 442 | 16 | 0.872 | 0.032 | True | Sulfate adenylyltransferase subunit 1 (EC 2.7.7.4) |
| PGF_06935032 | 18.32 | 434 | 16 | 0.879 | 0.015 | True | Adenylosuccinate synthetase (EC 6.3.4.4) |
| PGF_00294865 | 18.30 | 566 | 16 | 0.769 | 0.052 | True | Two component system sensor histidine kinase MtrB |
| PGF_00071194 | 18.28 | 487 | 16 | 0.828 | 0.070 | True | 2-keto-3-deoxy-D-arabino-heptulosonate-7-phosphate synthase II (EC 2.5.1.54) |
| PGF_00760084 | 18.26 | 508 | 16 | 0.810 | 0.080 | True | Cystathionine beta-synthase (EC 4.2.1.22) |
| PGF_07760799 | 18.26 | 497 | 16 | 0.819 | 0.027 | True | Pyruvate kinase (EC 2.7.1.40) |
| PGF_00423429 | 18.11 | 581 | 16 | 0.751 | 0.050 | True | Dihydroxyacetone kinase-like protein, phosphatase domain / Dihydroxyacetone kinase-like protein, kinase domain |
| PGF_00426236 | 18.10 | 392 | 16 | 0.914 | 0.023 | True | (E)-4-hydroxy-3-methylbut-2-enyl-diphosphate synthase (flavodoxin) (EC 1.17.7.3) |
| PGF_00417840 | 18.09 | 393 | 16 | 0.912 | 0.002 | True | Chorismate synthase (EC 4.2.3.5) |
| PGF_04560429 | 18.04 | 444 | 16 | 0.856 | 0.038 | True | Mg/Co/Ni transporter MgtE, CBS domain-containing |
| PGF_05935795 | 18.03 | 497 | 16 | 0.809 | 0.075 | True | Replication-associated recombination protein RarA |
| PGF_00064395 | 18.02 | 480 | 16 | 0.822 | 0.026 | True | UTP--glucose-1-phosphate uridylyltransferase (EC 2.7.7.9) |
| PGF_00422271 | 17.97 | 339 | 16 | 0.976 | 0.003 | True | DNA-directed RNA polymerase alpha subunit (EC 2.7.7.6) |
| PGF_00066964 | 17.94 | 496 | 16 | 0.805 | 0.010 | True | Xanthine/uracil/thiamine/ascorbate permease family protein |
| PGF_00421792 | 17.93 | 508 | 16 | 0.796 | 0.076 | True | DNA repair protein RadA |
| PGF_00006461 | 17.92 | 478 | 16 | 0.820 | 0.031 | True | Fumarate hydratase class II (EC 4.2.1.2) |
| PGF_00047078 | 17.88 | 352 | 16 | 0.953 | 0.007 | True | RecA protein |
| PGF_10461681 | 17.85 | 1550 | 16 | 0.453 | 0.290 | True | Ribonuclease E (EC 3.1.26.12) |
| PGF_00829251 | 17.83 | 609 | 16 | 0.722 | 0.082 | True | 3-hydroxyacyl-CoA dehydrogenase (EC 1.1.1.35) / 3-hydroxyacyl-CoA dehydrogenase (EC 1.1.1.35) |
| PGF_00008337 | 17.77 | 504 | 16 | 0.792 | 0.033 | True | Glutamyl-tRNA synthetase (EC 6.1.1.17) @ Glutamyl-tRNA(Gln) synthetase (EC 6.1.1.24) |
| PGF_00008075 | 17.77 | 383 | 16 | 0.908 | 0.000 | True | Glutamate N-acetyltransferase (EC 2.3.1.35) @ N-acetylglutamate synthase (EC 2.3.1.1) |
| PGF_00054528 | 17.66 | 407 | 16 | 0.875 | 0.039 | True | Succinyl-CoA ligase [ADP-forming] beta chain (EC 6.2.1.5) |
| PGF_00014381 | 17.66 | 362 | 16 | 0.928 | 0.005 | True | Inositol-1-phosphate synthase (EC 5.5.1.4) |
| PGF_10365126 | 17.63 | 955 | 16 | 0.571 | 0.241 | True | DNA polymerase III subunits gamma and tau (EC 2.7.7.7) |
| PGF_00048846 | 17.63 | 573 | 16 | 0.737 | 0.118 | True | Ribosomal protein S12p Asp88 (E. coli) methylthiotransferase (EC 2.8.4.4) |
| PGF_00062027 | 17.58 | 446 | 16 | 0.832 | 0.041 | True | Tryptophan synthase beta chain (EC 4.2.1.20) |
| PGF_02278006 | 17.55 | 398 | 16 | 0.879 | 0.014 | True | Quinolinate synthetase (EC 2.5.1.72) |
| PGF_00420081 | 17.54 | 418 | 16 | 0.858 | 0.043 | True | Cysteine synthesis adenylyltransferase/sulfurtransferase |
| PGF_00022722 | 17.52 | 389 | 16 | 0.888 | 0.008 | True | Mrp protein homolog |
| PGF_02430085 | 17.50 | 407 | 16 | 0.867 | 0.006 | True | Acyl-CoA dehydrogenase (EC 1.3.8.1), Mycobacterial subgroup FadE23 |
| PGF_00014314 | 17.46 | 384 | 16 | 0.891 | 0.037 | True | Inosine-5'-monophosphate dehydrogenase, catalytic domain (EC 1.1.1.205) |
| PGF_06377994 | 17.44 | 489 | 16 | 0.789 | 0.043 | True | FtsW-like cell division membrane protein CA_C0505 |
| PGF_02797400 | 17.43 | 512 | 16 | 0.771 | 0.083 | True | Cysteinyl-tRNA synthetase (EC 6.1.1.16) |
| PGF_03520151 | 17.41 | 440 | 16 | 0.830 | 0.070 | True | Cell division protein FtsZ |
| PGF_05622049 | 17.41 | 473 | 16 | 0.800 | 0.036 | True | UPF0053 protein Rv1842c/MT1890 |
| PGF_07583562 | 17.40 | 429 | 16 | 0.840 | 0.035 | True | Cysteine desulfurase (EC 2.8.1.7) => SufS |
| PGF_00042294 | 17.39 | 515 | 16 | 0.766 | 0.043 | True | Putative methyltransferase SCO3545 |
| PGF_00066867 | 17.36 | 521 | 16 | 0.761 | 0.101 | True | Xaa-Pro aminopeptidase (EC 3.4.11.9) |
| PGF_00064046 | 17.35 | 408 | 16 | 0.859 | 0.024 | True | UDP-galactopyranose mutase (EC 5.4.99.9) |
| PGF_00063999 | 17.35 | 474 | 16 | 0.797 | 0.011 | True | UDP-N-acetylmuramate--L-alanine ligase (EC 6.3.2.8) |
| PGF_00007012 | 17.34 | 358 | 16 | 0.916 | 0.000 | True | GTP-binding and nucleic acid-binding protein YchF |
| PGF_00416674 | 17.33 | 456 | 16 | 0.811 | 0.073 | True | 3,4-dihydroxy-2-butanone 4-phosphate synthase (EC 4.1.99.12) / GTP cyclohydrolase II (EC 3.5.4.25) |
| PGF_00024274 | 17.32 | 398 | 16 | 0.868 | 0.022 | True | N5-carboxyaminoimidazole ribonucleotide synthase (EC 6.3.4.18) |
| PGF_04456509 | 17.32 | 569 | 16 | 0.726 | 0.059 | True | 2-succinyl-5-enolpyruvyl-6-hydroxy-3-cyclohexene-1-carboxylic-acid synthase (EC 2.2.1.9) |
| PGF_00051525 | 17.29 | 435 | 16 | 0.829 | 0.027 | True | Seryl-tRNA synthetase (EC 6.1.1.11) |
| PGF_02277678 | 17.29 | 419 | 16 | 0.845 | 0.038 | True | Phosphoglycerate kinase (EC 2.7.2.3) |
| PGF_00648450 | 17.26 | 347 | 16 | 0.927 | 0.011 | True | Rod shape-determining protein MreB |
| PGF_09019918 | 17.26 | 486 | 16 | 0.783 | 0.078 | True | Glutamate-1-semialdehyde 2,1-aminomutase (EC 5.4.3.8) |
| PGF_00425738 | 17.25 | 328 | 16 | 0.952 | 0.000 | True | Sporulation transcription regulator WhiA |
| PGF_00618776 | 17.24 | 392 | 16 | 0.871 | 0.035 | True | 23S rRNA (adenine(2503)-C(2))-methyltransferase @ tRNA (adenine(37)-C(2))-methyltransferase (EC 2.1.1.192) |
| PGF_00420975 | 17.23 | 428 | 16 | 0.833 | 0.022 | True | D-inositol-3-phosphate glycosyltransferase (EC 2.4.1.250) |
| PGF_00421347 | 17.22 | 364 | 16 | 0.903 | 0.010 | True | DNA integrity scanning protein DisA |
| PGF_00015855 | 17.11 | 342 | 16 | 0.925 | 0.000 | True | Ketol-acid reductoisomerase (NADP(+)) (EC 1.1.1.86) |
| PGF_03659350 | 17.06 | 580 | 16 | 0.708 | 0.088 | True | Protein-O-mannosyltransferase |
| PGF_05760069 | 17.04 | 449 | 16 | 0.804 | 0.024 | True | Histidinol dehydrogenase (EC 1.1.1.23) |
| PGF_02019462 | 17.01 | 371 | 16 | 0.883 | 0.013 | True | Phenylalanyl-tRNA synthetase alpha chain (EC 6.1.1.20) |
| PGF_00016074 | 17.00 | 413 | 16 | 0.836 | 0.006 | True | L-cysteine:1D-myo-inosityl 2-amino-2-deoxy-alpha-D-glucopyranoside ligase MshC (EC 6.3.1.13) |
| PGF_10302926 | 17.00 | 621 | 16 | 0.682 | 0.085 | True | Cell division protein FtsI [Peptidoglycan synthetase] (EC 2.4.1.129) |
| PGF_00912265 | 16.98 | 394 | 16 | 0.855 | 0.034 | True | tRNA-dihydrouridine synthase DusB |
| PGF_05950073 | 16.95 | 413 | 16 | 0.834 | 0.056 | True | Cystathionine gamma-lyase (EC 4.4.1.1) |
| PGF_00063916 | 16.95 | 438 | 16 | 0.810 | 0.036 | True | Tyrosyl-tRNA synthetase (EC 6.1.1.1) |
| PGF_05877811 | 16.94 | 537 | 16 | 0.731 | 0.010 | True | AAA+ ATPase superfamily protein YifB/ComM, associated with DNA recombination |
| PGF_05387084 | 16.92 | 430 | 16 | 0.816 | 0.028 | True | Gamma-glutamyl phosphate reductase (EC 1.2.1.41) |
| PGF_05049118 | 16.89 | 377 | 16 | 0.870 | 0.038 | True | Holliday junction ATP-dependent DNA helicase RuvB (EC 3.6.4.12) |
| PGF_05572316 | 16.88 | 460 | 16 | 0.787 | 0.051 | True | FIG007959: peptidase, M16 family |
| PGF_10511254 | 16.87 | 498 | 16 | 0.756 | 0.022 | True | FtsI-like cell elongation transpeptidase |
| PGF_03858132 | 16.86 | 551 | 16 | 0.718 | 0.098 | True | Methylphosphotriester-DNA--protein-cysteine S-methyltransferase (EC 2.1.1.n11) / DNA-3-methyladenine glycosylase II (EC 3.2.2.21) |
| PGF_00404271 | 16.86 | 1026 | 16 | 0.526 | 0.395 | True | Biotin carboxylase of acetyl-CoA carboxylase (EC 6.3.4.14) / Biotin carboxyl carrier protein of acetyl-CoA carboxylase |
| PGF_03219041 | 16.86 | 420 | 16 | 0.822 | 0.015 | True | putative phospholipase D family protein |
| PGF_01557745 | 16.84 | 421 | 16 | 0.821 | 0.042 | True | Threonine dehydratase, catabolic (EC 4.3.1.19) @ L-serine dehydratase, (PLP)-dependent (EC 4.3.1.17) |
| PGF_10143857 | 16.80 | 344 | 16 | 0.906 | 0.001 | True | Fructose-bisphosphate aldolase class II (EC 4.1.2.13) |
| PGF_01368025 | 16.78 | 384 | 16 | 0.856 | 0.034 | True | Formaldehyde dehydrogenase MscR, NAD/mycothiol-dependent (EC 1.2.1.66) / S-nitrosomycothiol reductase MscR |
| PGF_00425024 | 16.77 | 435 | 16 | 0.804 | 0.042 | True | Exodeoxyribonuclease VII large subunit (EC 3.1.11.6) |
| PGF_03428301 | 16.71 | 486 | 16 | 0.758 | 0.055 | True | Acyl-CoA dehydrogenase (EC 1.3.8.1), Mycobacterial subgroup FadE24 |
| PGF_00030640 | 16.71 | 362 | 16 | 0.878 | 0.013 | True | Peptide chain release factor 1 |
| PGF_12771936 | 16.67 | 454 | 16 | 0.782 | 0.062 | True | dNTP triphosphohydrolase, broad substrate specificity |
| PGF_06098633 | 16.67 | 429 | 16 | 0.805 | 0.022 | True | Uncharacterized DUF349-containing protein SCO1511 |
| PGF_00218860 | 16.66 | 378 | 16 | 0.857 | 0.009 | True | Two-component system sensor histidine kinase |
| PGF_00021023 | 16.63 | 648 | 16 | 0.653 | 0.076 | True | Methylmalonyl-CoA mutase small subunit, MutA (EC 5.4.99.2) |
| PGF_00033987 | 16.60 | 384 | 16 | 0.847 | 0.026 | True | Phosphoserine aminotransferase (EC 2.6.1.52) |
| PGF_00754619 | 16.56 | 520 | 16 | 0.726 | 0.064 | True | 3'-to-5' exoribonuclease RNase R |
| PGF_08173994 | 16.56 | 451 | 16 | 0.780 | 0.060 | True | Glycosyltransferase SCO2318 |
| PGF_02881268 | 16.54 | 428 | 16 | 0.800 | 0.060 | True | RecA-superfamily ATPases implicated in signal transduction |
| PGF_00767262 | 16.54 | 655 | 16 | 0.646 | 0.098 | True | Exodeoxyribonuclease V alpha chain (EC 3.1.11.5) |
| PGF_09111052 | 16.54 | 443 | 16 | 0.786 | 0.076 | True | Phosphopantothenoylcysteine decarboxylase (EC 4.1.1.36) / Phosphopantothenoylcysteine synthetase (EC 6.3.2.5) |
| PGF_00420080 | 16.53 | 325 | 16 | 0.917 | 0.024 | True | Cysteine synthase, CysO-dependent |
| PGF_07844318 | 16.52 | 443 | 16 | 0.785 | 0.032 | True | 3-phosphoshikimate 1-carboxyvinyltransferase (EC 2.5.1.19) |
| PGF_00421679 | 16.51 | 1003 | 16 | 0.521 | 0.321 | True | DNA primase DnaG |
| PGF_00017259 | 16.51 | 586 | 16 | 0.682 | 0.143 | True | ATP-dependent RNA helicase |
| PGF_00048782 | 16.47 | 326 | 16 | 0.912 | 0.000 | True | Ribose-phosphate pyrophosphokinase (EC 2.7.6.1) |
| PGF_09188652 | 16.47 | 428 | 16 | 0.796 | 0.040 | True | Molybdopterin molybdenumtransferase (EC 2.10.1.1) |
| PGF_04139053 | 16.44 | 363 | 16 | 0.863 | 0.035 | True | Uroporphyrinogen III decarboxylase (EC 4.1.1.37) |
| PGF_00404376 | 16.41 | 434 | 16 | 0.788 | 0.022 | True | probable membrane protein STY4873 |
| PGF_07015581 | 16.38 | 421 | 16 | 0.799 | 0.083 | True | Carbamoyl-phosphate synthase small chain (EC 6.3.5.5) |
| PGF_00017265 | 16.38 | 607 | 16 | 0.665 | 0.198 | True | ATP-dependent RNA helicase |
| PGF_04991657 | 16.34 | 430 | 16 | 0.788 | 0.051 | True | Oxygen-independent coproporphyrinogen-III oxidase-like protein YggW |
| PGF_03004613 | 16.33 | 486 | 16 | 0.741 | 0.059 | True | Histidyl-tRNA synthetase (EC 6.1.1.21) |
| PGF_01062603 | 16.30 | 393 | 16 | 0.822 | 0.077 | True | Malyl-CoA lyase (EC 4.1.3.24) |
| PGF_00046352 | 16.29 | 328 | 16 | 0.900 | 0.024 | True | RNA polymerase sigma factor SigB |
| PGF_09994213 | 16.28 | 364 | 16 | 0.853 | 0.009 | True | Glycerol-3-phosphate ABC transporter, ATP-binding protein UgpC (TC 3.A.1.1.3) |
| PGF_07191648 | 16.26 | 373 | 16 | 0.842 | 0.037 | True | 3-isopropylmalate dehydrogenase (EC 1.1.1.85) |
| PGF_07063065 | 16.23 | 329 | 16 | 0.895 | 0.010 | True | Transcription termination protein NusA |
| PGF_00067188 | 16.20 | 356 | 16 | 0.858 | 0.023 | True | Aspartate-semialdehyde dehydrogenase (EC 1.2.1.11) |
| PGF_00016393 | 16.20 | 278 | 16 | 0.971 | 0.000 | True | LSU ribosomal protein L2p (L8e) |
| PGF_00033954 | 16.18 | 377 | 16 | 0.833 | 0.046 | True | Phosphoribosylformylglycinamidine cyclo-ligase (EC 6.3.3.1) |
| PGF_08376928 | 16.17 | 373 | 16 | 0.837 | 0.034 | True | Branched-chain amino acid aminotransferase (EC 2.6.1.42) |
| PGF_07024091 | 16.14 | 396 | 16 | 0.811 | 0.027 | True | coenzyme F420-dependent N5,N10-methylene tetrahydromethanopterin reductase |
| PGF_03316046 | 16.12 | 371 | 16 | 0.837 | 0.032 | True | Threonine synthase (EC 4.2.3.1) |
| PGF_04999088 | 16.11 | 494 | 16 | 0.725 | 0.094 | True | Uncharacterized protein SCO5199 |
| PGF_00048928 | 16.09 | 341 | 16 | 0.871 | 0.021 | True | Ribosome small subunit biogenesis RbfA-release protein RsgA |
| PGF_07235150 | 16.02 | 599 | 16 | 0.655 | 0.083 | True | Peptidoglycan lipid II flippase MurJ |
| PGF_00025600 | 16.02 | 481 | 16 | 0.730 | 0.090 | True | Nicotinate phosphoribosyltransferase (EC 6.3.4.21) |
| PGF_00024322 | 15.98 | 507 | 16 | 0.710 | 0.062 | True | NAD(P) transhydrogenase subunit beta (EC 1.6.1.2) |
| PGF_01022008 | 15.96 | 487 | 16 | 0.723 | 0.138 | True | Isochorismate synthase (EC 5.4.4.2) @ Menaquinone-specific isochorismate synthase (EC 5.4.4.2) |
| PGF_09129813 | 15.95 | 389 | 16 | 0.809 | 0.034 | True | Glutamate 5-kinase (EC 2.7.2.11) / RNA-binding C-terminal domain PUA |
| PGF_04797267 | 15.91 | 319 | 16 | 0.891 | 0.023 | True | Naphthoate synthase (EC 4.1.3.36) |
| PGF_03811905 | 15.91 | 398 | 16 | 0.797 | 0.061 | True | Histidinol-phosphate aminotransferase (EC 2.6.1.9) |
| PGF_01126005 | 15.90 | 422 | 16 | 0.774 | 0.062 | True | Rod shape-determining protein RodA |
| PGF_01021996 | 15.89 | 396 | 16 | 0.798 | 0.012 | True | Putative serine protease |
| PGF_08905885 | 15.87 | 412 | 16 | 0.782 | 0.073 | True | Signal recognition particle receptor FtsY |
| PGF_10244701 | 15.84 | 333 | 16 | 0.868 | 0.011 | True | GTP 3',8-cyclase (EC 4.1.99.22) |
| PGF_00008334 | 15.84 | 472 | 16 | 0.729 | 0.086 | True | Glutamyl-tRNA reductase (EC 1.2.1.70) |
| PGF_06230969 | 15.80 | 495 | 16 | 0.710 | 0.072 | True | NAD(P)H-hydrate epimerase (EC 5.1.99.6) / ADP-dependent (S)-NAD(P)H-hydrate dehydratase (EC 4.2.1.136) |
| PGF_08364774 | 15.77 | 351 | 16 | 0.842 | 0.037 | True | Fructose-1,6-bisphosphatase, GlpX type (EC 3.1.3.11) |
| PGF_08577075 | 15.76 | 336 | 16 | 0.860 | 0.001 | True | Potassium channel beta chain |
| PGF_02959749 | 15.74 | 473 | 16 | 0.724 | 0.077 | True | Ribonuclease D (EC 3.1.26.3) |
| PGF_01745391 | 15.72 | 384 | 16 | 0.802 | 0.056 | True | Cystathionine gamma-synthase (EC 2.5.1.48) |
| PGF_00017479 | 15.72 | 337 | 16 | 0.856 | 0.063 | True | Lipoyl synthase (EC 2.8.1.8) |
| PGF_00489714 | 15.70 | 343 | 16 | 0.848 | 0.032 | True | Porphobilinogen synthase (EC 4.2.1.24) |
| PGF_05053989 | 15.68 | 420 | 16 | 0.765 | 0.109 | True | ATP-dependent DNA ligase (EC 6.5.1.1) LigC |
| PGF_00008611 | 15.66 | 339 | 16 | 0.850 | 0.016 | True | Glycerol-3-phosphate dehydrogenase [NAD(P)+] (EC 1.1.1.94) |
| PGF_00024234 | 15.64 | 367 | 16 | 0.817 | 0.035 | True | N-succinyl-L,L-diaminopimelate desuccinylase (EC 3.5.1.18) |
| PGF_03439827 | 15.61 | 447 | 16 | 0.738 | 0.067 | True | Precorrin-2 oxidase (EC 1.3.1.76) @ Sirohydrochlorin ferrochelatase activity of CysG (EC 4.99.1.4) / Uroporphyrinogen-III methyltransferase (EC 2.1.1.107) |
| PGF_00853393 | 15.60 | 308 | 16 | 0.889 | 0.046 | True | Succinyl-CoA ligase [ADP-forming] alpha chain (EC 6.2.1.5) |
| PGF_00053770 | 15.59 | 395 | 16 | 0.785 | 0.060 | True | Aminomethyltransferase (glycine cleavage system T protein) (EC 2.1.2.10) |
| PGF_00024259 | 15.59 | 352 | 16 | 0.831 | 0.017 | True | N5,N10-methylenetetrahydromethanopterin reductase-related protein, MSMEG1563 family |
| PGF_01867628 | 15.57 | 348 | 16 | 0.835 | 0.023 | True | Heat-inducible transcription repressor HrcA |
| PGF_00418031 | 15.56 | 417 | 16 | 0.762 | 0.034 | True | Chromosome segregation ATPases |
| PGF_04521913 | 15.55 | 507 | 16 | 0.690 | 0.081 | True | UDP-N-acetylmuramoyl-tripeptide--D-alanyl-D-alanine ligase (EC 6.3.2.10) |
| PGF_00692117 | 15.54 | 424 | 16 | 0.755 | 0.085 | True | Anion-transporting ATPase Rv3680 |
| PGF_10569727 | 15.53 | 316 | 16 | 0.874 | 0.013 | True | LSU rRNA pseudouridine(1911/1915/1917) synthase (EC 5.4.99.23) |
| PGF_09077460 | 15.53 | 379 | 16 | 0.798 | 0.039 | True | 1-deoxy-D-xylulose 5-phosphate reductoisomerase (EC 1.1.1.267) |
| PGF_05463076 | 15.50 | 478 | 16 | 0.709 | 0.062 | True | Protoporphyrinogen IX oxidase, aerobic, HemY (EC 1.3.3.4) |
| PGF_08127242 | 15.49 | 423 | 16 | 0.753 | 0.069 | True | Glutamine-dependent 2-keto-4-methylthiobutyrate transaminase |
| PGF_10125426 | 15.46 | 483 | 16 | 0.704 | 0.108 | True | Peptidoglycan glycosyltransferase FtsW (EC 2.4.1.129) |
| PGF_00060021 | 15.42 | 347 | 16 | 0.828 | 0.012 | True | Transcriptional regulator |
| PGF_00963385 | 15.41 | 597 | 16 | 0.631 | 0.195 | True | putative membrane protein |
| PGF_00006510 | 15.38 | 441 | 16 | 0.733 | 0.079 | True | Fumarylacetoacetase (EC 3.7.1.2) |
| PGF_03295331 | 15.37 | 373 | 16 | 0.796 | 0.033 | True | UDP-N-acetylglucosamine--N-acetylmuramyl-(pentapeptide) pyrophosphoryl-undecaprenol N-acetylglucosamine transferase (EC 2.4.1.227) |
| PGF_10555225 | 15.35 | 449 | 16 | 0.724 | 0.072 | True | Ornithine aminotransferase (EC 2.6.1.13) |
| PGF_03062930 | 15.34 | 434 | 16 | 0.737 | 0.074 | True | Decaprenyl-phosphate N-acetylglucosaminephosphotransferase (EC 2.7.8.35) |
| PGF_02567701 | 15.34 | 508 | 16 | 0.681 | 0.081 | True | Two component system sensor histidine kinase MprB |
| PGF_00413208 | 15.34 | 407 | 16 | 0.760 | 0.039 | True | tRNA (guanine(37)-N(1))-methyltransferase (EC 2.1.1.228) |
| PGF_00038897 | 15.33 | 332 | 16 | 0.842 | 0.048 | True | Aldo/keto reductase, SCO1670 family |
| PGF_00057399 | 15.33 | 372 | 16 | 0.795 | 0.005 | True | Transaldolase (EC 2.2.1.2) |
| PGF_10505717 | 15.32 | 731 | 16 | 0.567 | 0.243 | True | Glucose-6-phosphate isomerase (EC 5.3.1.9) |
| PGF_04449783 | 15.25 | 347 | 16 | 0.819 | 0.059 | True | Pantothenate kinase (EC 2.7.1.33) |
| PGF_04149954 | 15.20 | 412 | 16 | 0.749 | 0.052 | True | putative membrane protein |
| PGF_00024232 | 15.15 | 403 | 16 | 0.755 | 0.074 | True | N-succinyl-L,L-diaminopimelate aminotransferase (EC 2.6.1.17), type 2 |
| PGF_00015259 | 15.13 | 320 | 16 | 0.846 | 0.046 | True | ATP synthase gamma chain (EC 3.6.3.14) |
| PGF_00007027 | 15.09 | 389 | 16 | 0.765 | 0.104 | True | GTP-binding protein Era |
| PGF_02866866 | 15.09 | 394 | 16 | 0.760 | 0.059 | True | Coproporphyrin ferrochelatase (EC 4.99.1.9) |
| PGF_00033237 | 15.08 | 384 | 16 | 0.770 | 0.093 | True | Phosphate starvation-inducible protein PhoH, predicted ATPase |
| PGF_00027514 | 15.06 | 422 | 16 | 0.733 | 0.053 | True | N-acetylornithine aminotransferase (EC 2.6.1.11) |
| PGF_00172869 | 15.06 | 400 | 16 | 0.753 | 0.054 | True | Alkaline phosphodiesterase I (EC 3.1.4.1) / Nucleotide pyrophosphatase (EC 3.6.1.9) |
| PGF_02390924 | 15.05 | 334 | 16 | 0.824 | 0.021 | True | 16S rRNA (cytosine(1402)-N(4))-methyltransferase (EC 2.1.1.199) |
| PGF_08151051 | 15.02 | 310 | 16 | 0.853 | 0.048 | True | Histidinol-phosphatase [alternative form] (EC 3.1.3.15) |
| PGF_00053513 | 14.99 | 340 | 16 | 0.813 | 0.020 | True | Solanesyl diphosphate synthase (EC 2.5.1.11) |
| PGF_00417658 | 14.96 | 412 | 16 | 0.737 | 0.099 | True | 3-dehydroquinate synthase (EC 4.2.3.4) |
| PGF_00026615 | 14.96 | 336 | 16 | 0.816 | 0.057 | True | N-acetylglutamate kinase (EC 2.7.2.8) |
| PGF_00054961 | 14.95 | 317 | 16 | 0.840 | 0.038 | True | Sulfate adenylyltransferase subunit 2 (EC 2.7.7.4) |
| PGF_00405726 | 14.95 | 350 | 16 | 0.799 | 0.049 | True | putative Adenosine kinase (EC 2.7.1.20) |
| PGF_00019068 | 14.93 | 516 | 16 | 0.657 | 0.093 | True | Maltokinase (EC 2.7.1.175) |
| PGF_00015517 | 14.91 | 442 | 16 | 0.709 | 0.093 | True | Iron-sulfur cluster assembly protein SufD |
| PGF_00049889 | 14.90 | 291 | 16 | 0.874 | 0.064 | True | SSU ribosomal protein S3p (S3e) |
| PGF_00052943 | 14.89 | 317 | 16 | 0.836 | 0.039 | True | Site-specific tyrosine recombinase XerD |
| PGF_00422259 | 14.89 | 375 | 16 | 0.769 | 0.071 | True | DNA-dependent DNA polymerase beta chain |
| PGF_01147190 | 14.88 | 492 | 16 | 0.671 | 0.101 | True | Deoxyribodipyrimidine photolyase (EC 4.1.99.3) |
| PGF_08582746 | 14.84 | 332 | 16 | 0.814 | 0.039 | True | 23S rRNA (guanosine(2251)-2'-O)-methyltransferase (EC 2.1.1.185) |
| PGF_09201380 | 14.83 | 332 | 16 | 0.814 | 0.012 | True | DNA polymerase III delta subunit (EC 2.7.7.7) |
| PGF_10125211 | 14.81 | 456 | 16 | 0.694 | 0.110 | True | Phosphate regulon sensor protein PhoR (SphS) (EC 2.7.13.3) |
| PGF_00121830 | 14.77 | 519 | 16 | 0.648 | 0.112 | True | FIG01121868: Possible membrane protein, Rv0205 |
| PGF_00413192 | 14.77 | 364 | 16 | 0.774 | 0.100 | True | tRNA (adenine(58)-N(1))-methyltransferase (EC 2.1.1.220) |
| PGF_00037521 | 14.77 | 316 | 16 | 0.831 | 0.015 | True | Proline dehydrogenase (EC 1.5.5.2) |
| PGF_03069128 | 14.76 | 714 | 16 | 0.553 | 0.207 | True | putative membrane protein |
| PGF_00057483 | 14.76 | 819 | 16 | 0.516 | 0.229 | True | Transcription termination factor Rho |
| PGF_07889681 | 14.76 | 392 | 16 | 0.746 | 0.114 | True | N-acetyl-gamma-glutamyl-phosphate reductase (EC 1.2.1.38) |
| PGF_07157721 | 14.67 | 332 | 16 | 0.805 | 0.043 | True | Heme O synthase, protoheme IX farnesyltransferase, COX10-CtaB |
| PGF_00056897 | 14.64 | 269 | 16 | 0.893 | 0.010 | True | Thymidylate synthase (EC 2.1.1.45) |
| PGF_00016850 | 14.63 | 415 | 16 | 0.718 | 0.116 | True | Branched-chain amino acid dehydrogenase [deaminating] (EC 1.4.1.9)(EC 1.4.1.23) |
| PGF_04310056 | 14.61 | 438 | 16 | 0.698 | 0.107 | True | Acyltransferase family protein |
| PGF_00049827 | 14.61 | 303 | 16 | 0.839 | 0.035 | True | SSU rRNA (adenine(1518)-N(6)/adenine(1519)-N(6))-dimethyltransferase (EC 2.1.1.182) |
| PGF_06671647 | 14.58 | 441 | 16 | 0.694 | 0.114 | True | N-formylglutamate deformylase (EC 3.5.1.68) [alternative form] |
| PGF_10387199 | 14.56 | 426 | 16 | 0.706 | 0.122 | True | DNA recombination and repair protein RecF |
| PGF_06755829 | 14.56 | 400 | 16 | 0.728 | 0.025 | True | DNA polymerase III delta prime subunit (EC 2.7.7.7) |
| PGF_00016357 | 14.55 | 240 | 16 | 0.939 | 0.001 | True | LSU ribosomal protein L1p (L10Ae) |
| PGF_00021125 | 14.54 | 453 | 16 | 0.683 | 0.100 | True | SAM-dependent methyltransferase |
| PGF_02191019 | 14.53 | 348 | 16 | 0.779 | 0.070 | True | Thiamine-monophosphate kinase (EC 2.7.4.16) |
| PGF_00006100 | 14.51 | 348 | 16 | 0.778 | 0.094 | True | tRNA-modifying protein YgfZ |
| PGF_00023532 | 14.49 | 309 | 16 | 0.824 | 0.034 | True | Mycothiol S-conjugate amidase Mca |
| PGF_00063974 | 14.47 | 360 | 16 | 0.763 | 0.027 | True | UDP-N-acetylenolpyruvoylglucosamine reductase (EC 1.3.1.98) |
| PGF_00033936 | 14.44 | 308 | 16 | 0.823 | 0.031 | True | Phosphoribosylaminoimidazole-succinocarboxamide synthase (EC 6.3.2.6) |
| PGF_06936561 | 14.41 | 271 | 16 | 0.875 | 0.004 | True | Translation elongation factor Ts |
| PGF_00423089 | 14.41 | 383 | 16 | 0.736 | 0.084 | True | Diaminohydroxyphosphoribosylaminopyrimidine deaminase (EC 3.5.4.26) / 5-amino-6-(5-phosphoribosylamino)uracil reductase (EC 1.1.1.193) |
| PGF_04835795 | 14.40 | 337 | 16 | 0.785 | 0.058 | True | Site-specific tyrosine recombinase XerC |
| PGF_04150742 | 14.38 | 344 | 16 | 0.776 | 0.027 | True | Zinc ABC transporter, permease protein ZnuB |
| PGF_03021654 | 14.38 | 480 | 16 | 0.656 | 0.133 | True | Anthranilate synthase, aminase component (EC 4.1.3.27) @ Para-aminobenzoate synthase, aminase component (EC 2.6.1.85) |
| PGF_02623406 | 14.36 | 326 | 16 | 0.795 | 0.039 | True | FMN adenylyltransferase (EC 2.7.7.2) / Riboflavin kinase (EC 2.7.1.26) |
| PGF_00066906 | 14.35 | 335 | 16 | 0.784 | 0.066 | True | Aspartate carbamoyltransferase (EC 2.1.3.2) |
| PGF_02011760 | 14.35 | 263 | 16 | 0.885 | 0.029 | True | Imidazole glycerol phosphate synthase cyclase subunit |
| PGF_03591205 | 14.33 | 528 | 16 | 0.624 | 0.190 | True | Serine/threonine phosphatase PPP (EC 3.1.3.16) |
| PGF_07609122 | 14.32 | 276 | 16 | 0.862 | 0.032 | True | Pyrroline-5-carboxylate reductase (EC 1.5.1.2) |
| PGF_03374975 | 14.30 | 339 | 16 | 0.777 | 0.062 | True | O-succinylbenzoate synthase (EC 4.2.1.113) |
| PGF_06684654 | 14.30 | 359 | 16 | 0.755 | 0.074 | True | UDP-glucose 4-epimerase (EC 5.1.3.2) |
| PGF_05165078 | 14.29 | 332 | 16 | 0.784 | 0.058 | True | Methionyl-tRNA formyltransferase (EC 2.1.2.9) |
| PGF_00020173 | 14.29 | 341 | 16 | 0.774 | 0.080 | True | 2,3,4,5-tetrahydropyridine-2,6-dicarboxylate N-succinyltransferase (EC 2.3.1.117) |
| PGF_04505269 | 14.27 | 395 | 16 | 0.718 | 0.154 | True | SSU ribosomal protein S2p (SAe) |
| PGF_00907900 | 14.25 | 225 | 16 | 0.950 | 0.000 | True | cAMP-binding proteins - catabolite gene activator and regulatory subunit of cAMP-dependent protein kinases |
| PGF_10097367 | 14.24 | 339 | 16 | 0.773 | 0.059 | True | Porphobilinogen deaminase (EC 2.5.1.61) |
| PGF_00001910 | 14.23 | 298 | 16 | 0.824 | 0.052 | True | Uncharacterized protien SCO2557 |
| PGF_00415308 | 14.19 | 279 | 16 | 0.850 | 0.001 | True | 23S rRNA (cytidine(1920)-2'-O)-methyltransferase (EC 2.1.1.226) @ 16S rRNA (cytidine(1409)-2'-O)-methyltransferase (EC 2.1.1.227) |
| PGF_00014855 | 14.17 | 289 | 16 | 0.834 | 0.026 | True | ATP phosphoribosyltransferase (EC 2.4.2.17) => HisGl |
| PGF_00037867 | 14.16 | 302 | 16 | 0.815 | 0.077 | True | Proteasome subunit beta (EC 3.4.25.1), bacterial |
| PGF_08125688 | 14.16 | 378 | 16 | 0.728 | 0.094 | True | Anion-transporting ATPase Rv3679 |
| PGF_02450432 | 14.15 | 306 | 16 | 0.809 | 0.029 | True | Phosphatidate cytidylyltransferase (EC 2.7.7.41) |
| PGF_09242109 | 14.15 | 262 | 16 | 0.874 | 0.013 | True | 3-hydroxyacyl-CoA dehydrogenase |
| PGF_01033770 | 14.15 | 394 | 16 | 0.713 | 0.115 | True | Dihydroorotate dehydrogenase (quinone) (EC 1.3.5.2) |
| PGF_05767868 | 14.14 | 262 | 16 | 0.874 | 0.012 | True | Probable transcriptional regulatory protein YebC |
| PGF_03152833 | 14.14 | 369 | 16 | 0.736 | 0.143 | True | Ku domain protein |
| PGF_01053539 | 14.13 | 337 | 16 | 0.770 | 0.051 | True | Inner membrane protein YihY, formerly thought to be RNase BN |
| PGF_04574228 | 14.10 | 274 | 16 | 0.852 | 0.029 | True | Triosephosphate isomerase (EC 5.3.1.1) |
| PGF_03788368 | 14.05 | 310 | 16 | 0.798 | 0.060 | True | RNase adapter protein RapZ |
| PGF_08421732 | 14.05 | 347 | 16 | 0.754 | 0.080 | True | NAD kinase (EC 2.7.1.23) |
| PGF_12840005 | 14.02 | 374 | 16 | 0.725 | 0.050 | True | Two-component system sensor histidine kinase |
| PGF_00048586 | 14.01 | 260 | 16 | 0.869 | 0.056 | True | Ribonuclease PH (EC 2.7.7.56) |
| PGF_00033950 | 14.00 | 260 | 16 | 0.868 | 0.044 | True | Phosphoribosylformimino-5-aminoimidazole carboxamide ribotide isomerase (EC 5.3.1.16) @ Acting phosphoribosylanthranilate isomerase (EC 5.3.1.24) |
| PGF_01534078 | 13.99 | 430 | 16 | 0.675 | 0.083 | True | Murein endolytic transglycosylase MltG |
| PGF_00014051 | 13.95 | 268 | 16 | 0.852 | 0.014 | True | Indole-3-glycerol phosphate synthase (EC 4.1.1.48) |
| PGF_00048829 | 13.90 | 267 | 16 | 0.851 | 0.009 | True | LSU rRNA pseudouridine(2605) synthase (EC 5.4.99.22) |
| PGF_00769755 | 13.89 | 497 | 16 | 0.623 | 0.146 | True | 16S rRNA (cytosine(967)-C(5))-methyltransferase (EC 2.1.1.176) |
| PGF_00033363 | 13.85 | 248 | 16 | 0.879 | 0.003 | True | Phosphoadenylyl-sulfate reductase [thioredoxin] (EC 1.8.4.8) |
| PGF_02930420 | 13.81 | 410 | 16 | 0.682 | 0.116 | True | Ubiquinol-cytochrome C reductase iron-sulfur subunit (EC 1.10.2.2) |
| PGF_02703381 | 13.78 | 247 | 16 | 0.877 | 0.004 | True | Phosphoglycerate mutase (EC 5.4.2.11) |
| PGF_03215471 | 13.77 | 317 | 16 | 0.774 | 0.109 | True | 16S rRNA (cytidine(1402)-2'-O)-methyltransferase (EC 2.1.1.198) |
| PGF_00041181 | 13.71 | 358 | 16 | 0.725 | 0.123 | True | Putative glycosyltransferase |
| PGF_08005402 | 13.71 | 349 | 16 | 0.734 | 0.072 | True | FIG002813: LPPG:FO 2-phospho-L-lactate transferase like, CofD-like |
| PGF_02517283 | 13.71 | 318 | 16 | 0.769 | 0.070 | True | Segregation and condensation protein A |
| PGF_01996161 | 13.70 | 666 | 16 | 0.531 | 0.080 | True | Flp pilus assembly protein TadB |
| PGF_02472178 | 13.70 | 319 | 16 | 0.767 | 0.071 | True | Uncharacterized metal-dependent hydrolase YcfH |
| PGF_00687197 | 13.68 | 316 | 16 | 0.770 | 0.042 | True | 1D-myo-inositol 2-acetamido-2-deoxy-alpha-D-glucopyranoside deacetylase (EC 3.5.1.103) |
| PGF_04409877 | 13.66 | 364 | 16 | 0.716 | 0.078 | True | Thiamin ABC transporter, ATPase component |
| PGF_00012356 | 13.66 | 313 | 16 | 0.772 | 0.042 | True | Homoserine kinase (EC 2.7.1.39) |
| PGF_00064454 | 13.65 | 302 | 16 | 0.786 | 0.042 | True | Ubiquinol-cytochrome C reductase, diheme cytochrome cc subunit |
| PGF_00415545 | 13.61 | 416 | 16 | 0.667 | 0.090 | True | 23S rRNA (uracil(747)-C(5))-methyltransferase (EC 2.1.1.189) |
| PGF_12831525 | 13.60 | 261 | 16 | 0.842 | 0.047 | True | SAM-dependent methyltransferase SCO2317, type 11 |
| PGF_06594113 | 13.59 | 541 | 16 | 0.584 | 0.107 | True | D-alanyl-D-alanine carboxypeptidase (EC 3.4.16.4) |
| PGF_02893929 | 13.57 | 271 | 16 | 0.824 | 0.050 | True | 3'(2'),5'-bisphosphate nucleotidase (EC 3.1.3.7) |
| PGF_02895544 | 13.55 | 223 | 16 | 0.908 | 0.003 | True | TrkA-like protein |
| PGF_04094270 | 13.54 | 220 | 16 | 0.913 | 0.001 | True | TrkA-like protein |
| PGF_02617708 | 13.53 | 299 | 16 | 0.782 | 0.023 | True | A/G-specific adenine glycosylase (EC 3.2.2.-) |
| PGF_00038986 | 13.49 | 293 | 16 | 0.788 | 0.081 | True | Purine nucleoside phosphorylase (EC 2.4.2.1) |
| PGF_03952305 | 13.48 | 372 | 16 | 0.699 | 0.051 | True | Arogenate dehydrogenase (EC 1.3.1.43) |
| PGF_12734335 | 13.47 | 266 | 16 | 0.826 | 0.028 | True | hypothetical protein |
| PGF_07544077 | 13.46 | 326 | 16 | 0.746 | 0.128 | True | Prephenate dehydratase (EC 4.2.1.51) |
| PGF_03701810 | 13.44 | 303 | 16 | 0.772 | 0.001 | True | Cell-division-associated, ABC-transporter-like signaling protein FtsX |
| PGF_07420523 | 13.44 | 267 | 16 | 0.823 | 0.030 | True | Exodeoxyribonuclease III (EC 3.1.11.2) |
| PGF_05636894 | 13.41 | 327 | 16 | 0.742 | 0.064 | True | Rhamnosyl transferase |
| PGF_00059672 | 13.40 | 284 | 16 | 0.795 | 0.083 | True | (2E,6Z)-farnesyl diphosphate synthase (EC 2.5.1.68) |
| PGF_00033959 | 13.39 | 224 | 16 | 0.894 | 0.008 | True | Phosphoribosylformylglycinamidine synthase, glutamine amidotransferase subunit (EC 6.3.5.3) |
| PGF_07479808 | 13.38 | 333 | 16 | 0.733 | 0.082 | True | Ketopantoate reductase PanG (EC 1.1.1.169) |
| PGF_00549380 | 13.36 | 313 | 16 | 0.755 | 0.044 | True | 4-hydroxy-tetrahydrodipicolinate synthase (EC 4.3.3.7) |
| PGF_00407669 | 13.34 | 375 | 16 | 0.689 | 0.048 | True | Integral membrane protein |
| PGF_00876943 | 13.33 | 348 | 16 | 0.715 | 0.154 | True | 1,4-dihydroxy-2-naphthoate polyprenyltransferase (EC 2.5.1.74) |
| PGF_00013341 | 13.33 | 345 | 16 | 0.718 | 0.069 | True | Hypothetical protein of L-Asparaginase type 2-like superfamily |
| PGF_00023514 | 13.33 | 239 | 16 | 0.862 | 0.031 | True | Mycobacterial persistence response regulator MprA |
| PGF_00523822 | 13.32 | 327 | 16 | 0.736 | 0.057 | True | Pantoate--beta-alanine ligase (EC 6.3.2.1) |
| PGF_06461498 | 13.31 | 281 | 16 | 0.794 | 0.023 | True | Orotidine 5'-phosphate decarboxylase (EC 4.1.1.23) |
| PGF_04010787 | 13.30 | 344 | 16 | 0.717 | 0.094 | True | Uncharacterized metalohydrolase SCO3582 |
| PGF_00413295 | 13.29 | 303 | 16 | 0.764 | 0.054 | True | tRNA pseudouridine(55) synthase (EC 5.4.99.25) |
| PGF_05026985 | 13.28 | 339 | 16 | 0.721 | 0.097 | True | Deoxyribose-phosphate aldolase (EC 4.1.2.4) |
| PGF_03863088 | 13.27 | 331 | 16 | 0.729 | 0.056 | True | FIG000875: Thioredoxin domain-containing protein EC-YbbN |
| PGF_10464361 | 13.27 | 250 | 16 | 0.839 | 0.066 | True | Two component system response regulator MtrA |
| PGF_00049893 | 13.25 | 202 | 16 | 0.932 | 0.000 | True | SSU ribosomal protein S4p (S9e) @ SSU ribosomal protein S4p (S9e), zinc-independent |
| PGF_10401812 | 13.25 | 220 | 16 | 0.893 | 0.014 | True | Thymidine kinase (EC 2.7.1.21) |
| PGF_06216244 | 13.24 | 260 | 16 | 0.821 | 0.058 | True | DNA recombination and repair protein RecO |
| PGF_02463284 | 13.24 | 199 | 16 | 0.938 | 0.004 | True | Recombination protein RecR |
| PGF_00065324 | 13.24 | 276 | 16 | 0.797 | 0.036 | True | Uncharacterized protein SCO3347 |
| PGF_00420383 | 13.23 | 357 | 16 | 0.700 | 0.122 | True | Cytochrome c oxidase caa3-type assembly factor CtaG_BS (unrelated to Cox11-CtaG family) |
| PGF_04213876 | 13.23 | 264 | 16 | 0.814 | 0.028 | True | Pantothenate kinase type III, CoaX-like (EC 2.7.1.33) |
| PGF_00403095 | 13.20 | 213 | 16 | 0.905 | 0.000 | True | Uracil phosphoribosyltransferase (EC 2.4.2.9) |
| PGF_05580933 | 13.17 | 301 | 16 | 0.759 | 0.071 | True | Peptide chain release factor N(5)-glutamine methyltransferase (EC 2.1.1.297) |
| PGF_00057381 | 13.16 | 305 | 16 | 0.753 | 0.135 | True | Trans,polycis-decaprenyl diphosphate synthase (EC 2.5.1.86) |
| PGF_10245672 | 13.11 | 225 | 16 | 0.874 | 0.021 | True | Ribulose-phosphate 3-epimerase (EC 5.1.3.1) |
| PGF_00062023 | 13.06 | 281 | 16 | 0.779 | 0.047 | True | Tryptophan synthase alpha chain (EC 4.2.1.20) |
| PGF_00016431 | 13.06 | 229 | 16 | 0.863 | 0.035 | True | LSU ribosomal protein L3p (L3e) |
| PGF_02897933 | 13.04 | 255 | 16 | 0.817 | 0.046 | True | Nitrogen metabolism regulator GlnR, OmpR family |
| PGF_05091456 | 13.03 | 313 | 16 | 0.736 | 0.034 | True | Uncharacterized inner membrane protein RarD |
| PGF_00049896 | 13.02 | 213 | 16 | 0.892 | 0.033 | True | SSU ribosomal protein S5p (S2e) |
| PGF_04807486 | 13.02 | 337 | 16 | 0.709 | 0.085 | True | tRNA dimethylallyltransferase (EC 2.5.1.75) |
| PGF_00413290 | 12.99 | 285 | 16 | 0.770 | 0.027 | True | tRNA pseudouridine(38-40) synthase (EC 5.4.99.12) |
| PGF_00015467 | 12.99 | 241 | 16 | 0.837 | 0.044 | True | Iron-dependent repressor IdeR/DtxR |
| PGF_01213071 | 12.98 | 223 | 16 | 0.869 | 0.023 | True | LSU ribosomal protein L10p (P0) |
| PGF_09087715 | 12.97 | 353 | 16 | 0.690 | 0.074 | True | tRNA(Ile)-lysidine synthetase (EC 6.3.4.19) |
| PGF_00424619 | 12.91 | 349 | 16 | 0.691 | 0.029 | True | 4-nitrophenylphosphatase (EC 3.1.3.41) |
| PGF_10348836 | 12.90 | 307 | 16 | 0.736 | 0.103 | True | NADH-ubiquinone oxidoreductase chain J (EC 1.6.5.3) |
| PGF_00845945 | 12.89 | 304 | 16 | 0.740 | 0.057 | True | DegV family protein |
| PGF_04007946 | 12.87 | 263 | 16 | 0.794 | 0.058 | True | Octanoate-[acyl-carrier-protein]-protein-N-octanoyltransferase (EC 2.3.1.181) |
| PGF_03810679 | 12.86 | 274 | 16 | 0.777 | 0.117 | True | SOS-response repressor and protease LexA (EC 3.4.21.88) |
| PGF_00042696 | 12.83 | 474 | 16 | 0.589 | 0.227 | True | Putative oxidoreductase |
| PGF_00422922 | 12.81 | 191 | 16 | 0.927 | 0.000 | True | Deoxycytidine triphosphate deaminase (EC 3.5.4.30) (dUMP-forming) |
| PGF_04380075 | 12.80 | 186 | 16 | 0.938 | 0.010 | True | NADH-ubiquinone oxidoreductase chain B (EC 1.6.5.3) |
| PGF_00006724 | 12.76 | 296 | 16 | 0.742 | 0.043 | True | GCN5-related N-acetyltransferase, FIGfam019367 |
| PGF_03003628 | 12.75 | 410 | 16 | 0.630 | 0.138 | True | putative membrane protein |
| PGF_00037865 | 12.73 | 362 | 16 | 0.669 | 0.161 | True | Proteasome subunit alpha (EC 3.4.25.1), bacterial |
| PGF_00016284 | 12.71 | 263 | 16 | 0.784 | 0.042 | True | LOG family protein |
| PGF_03574598 | 12.70 | 371 | 16 | 0.660 | 0.054 | True | Lon-like protease with PDZ domain |
| PGF_05949825 | 12.67 | 324 | 16 | 0.704 | 0.055 | True | Heme A synthase, cytochrome oxidase biogenesis protein Cox15-CtaA |
| PGF_00001331 | 12.64 | 266 | 16 | 0.775 | 0.127 | True | Endonuclease NucS |
| PGF_00016443 | 12.63 | 196 | 16 | 0.902 | 0.038 | True | LSU ribosomal protein L5p (L11e) |
| PGF_05344975 | 12.58 | 260 | 16 | 0.780 | 0.036 | True | YdfG-like protein |
| PGF_00956915 | 12.58 | 368 | 16 | 0.656 | 0.078 | True | O-succinylbenzoic acid--CoA ligase (EC 6.2.1.26) |
| PGF_00426616 | 12.58 | 360 | 16 | 0.663 | 0.086 | True | FIG005453: Putative DeoR-family transcriptional regulator |
| PGF_03799365 | 12.53 | 188 | 16 | 0.914 | 0.005 | True | Translation elongation factor P |
| PGF_00024615 | 12.52 | 327 | 16 | 0.693 | 0.062 | True | NADH pyrophosphatase (EC 3.6.1.22), decaps 5'-NAD modified RNA |
| PGF_00064272 | 12.48 | 276 | 16 | 0.751 | 0.058 | True | UPF0246 protein YaaA |
| PGF_01366870 | 12.48 | 320 | 16 | 0.698 | 0.101 | True | NUDIX hydrolase-like protein SCO3573 |
| PGF_00054323 | 12.46 | 207 | 16 | 0.866 | 0.000 | True | YciO protein, TsaC/YrdC paralog |
| PGF_00029310 | 12.46 | 254 | 16 | 0.782 | 0.089 | True | POSSIBLE RNA METHYLTRANSFERASE (RNA METHYLASE) |
| PGF_00016444 | 12.45 | 180 | 16 | 0.928 | 0.001 | True | LSU ribosomal protein L6p (L9e) |
| PGF_06648032 | 12.44 | 329 | 16 | 0.686 | 0.090 | True | RecB family exonuclease |
| PGF_03072132 | 12.42 | 161 | 16 | 0.979 | 0.000 | True | CarD-like transcriptional regulator |
| PGF_10351306 | 12.41 | 502 | 16 | 0.554 | 0.230 | True | DNA recombination protein RmuC |
| PGF_00419621 | 12.40 | 269 | 16 | 0.756 | 0.100 | True | Coproheme decarboxylase HemQ (no EC) |
| PGF_00054956 | 12.38 | 726 | 16 | 0.460 | 0.379 | True | Sulfate adenylyltransferase (EC 2.7.7.4) / Adenylylsulfate kinase (EC 2.7.1.25) |
| PGF_06660812 | 12.36 | 202 | 16 | 0.870 | 0.014 | True | 3-isopropylmalate dehydratase small subunit (EC 4.2.1.33) |
| PGF_08169626 | 12.35 | 247 | 16 | 0.786 | 0.036 | True | FIG00824290: FHA domain protein |
| PGF_10233208 | 12.35 | 253 | 16 | 0.777 | 0.026 | True | 16S rRNA (uracil(1498)-N(3))-methyltransferase (EC 2.1.1.193) |
| PGF_00013490 | 12.32 | 183 | 16 | 0.910 | 0.000 | True | Hypoxanthine-guanine phosphoribosyltransferase (EC 2.4.2.8) |
| PGF_01962206 | 12.32 | 301 | 16 | 0.710 | 0.084 | True | Uracil-DNA glycosylase, family 5 (EC 3.2.2.27) |
| PGF_00408775 | 12.29 | 535 | 16 | 0.531 | 0.310 | True | Potassium channel protein |
| PGF_08394843 | 12.27 | 278 | 16 | 0.736 | 0.129 | True | Endonuclease III (EC 4.2.99.18) |
| PGF_00047479 | 12.26 | 400 | 16 | 0.613 | 0.070 | True | Related to HTH domain of SpoOJ/ParA/ParB/repB family, involved in chromosome partitioning |
| PGF_00055665 | 12.23 | 415 | 16 | 0.601 | 0.201 | True | TPR-repeat-containing protein |
| PGF_02923127 | 12.21 | 245 | 16 | 0.780 | 0.029 | True | Uridylate kinase (EC 2.7.4.22) |
| PGF_01176589 | 12.20 | 270 | 16 | 0.742 | 0.072 | True | 4-hydroxy-tetrahydrodipicolinate reductase (EC 1.17.1.8) |
| PGF_10340618 | 12.19 | 411 | 16 | 0.601 | 0.129 | True | Potassium efflux system KefA protein / Small-conductance mechanosensitive channel |
| PGF_04828114 | 12.18 | 342 | 16 | 0.659 | 0.112 | True | Quinolinate phosphoribosyltransferase [decarboxylating] (EC 2.4.2.19) |
| PGF_03174068 | 12.10 | 339 | 16 | 0.657 | 0.171 | True | Transcription antitermination protein NusG |
| PGF_04867467 | 12.09 | 236 | 16 | 0.787 | 0.096 | True | Imidazole glycerol phosphate synthase amidotransferase subunit HisH |
| PGF_03753407 | 12.05 | 339 | 16 | 0.654 | 0.115 | True | tRNA threonylcarbamoyladenosine biosynthesis protein TsaE |
| PGF_04211832 | 12.02 | 262 | 16 | 0.743 | 0.041 | True | Polyphosphate glucokinase (EC 2.7.1.63) |
| PGF_10480638 | 11.98 | 225 | 16 | 0.799 | 0.086 | True | Cytochrome c oxidase polypeptide III (EC 1.9.3.1) |
| PGF_02787494 | 11.92 | 218 | 16 | 0.807 | 0.048 | True | Phosphate transport regulator (distant homolog of PhoU) |
| PGF_02516666 | 11.91 | 365 | 16 | 0.623 | 0.159 | True | Cytochrome c oxidase polypeptide II (EC 1.9.3.1) |
| PGF_03790040 | 11.90 | 292 | 16 | 0.696 | 0.139 | True | Ribonuclease III (EC 3.1.26.3) |
| PGF_00013803 | 11.90 | 211 | 16 | 0.819 | 0.041 | True | Imidazoleglycerol-phosphate dehydratase (EC 4.2.1.19) |
| PGF_06530721 | 11.87 | 286 | 16 | 0.702 | 0.146 | True | tRNA (guanine(46)-N(7))-methyltransferase (EC 2.1.1.33) |
| PGF_05880970 | 11.87 | 204 | 16 | 0.831 | 0.022 | True | Cardiolipin synthase (CMP-forming), eukaryotic type Cls-II (EC 2.7.8.41) |
| PGF_05843198 | 11.85 | 256 | 16 | 0.741 | 0.063 | True | FIG017108: hypothetical protein |
| PGF_01898991 | 11.84 | 239 | 16 | 0.766 | 0.070 | True | Threonylcarbamoyl-AMP synthase (EC 2.7.7.87) |
| PGF_03189552 | 11.82 | 200 | 16 | 0.836 | 0.006 | True | Holliday junction ATP-dependent DNA helicase RuvA (EC 3.6.4.12) |
| PGF_03647550 | 11.78 | 163 | 16 | 0.923 | 0.000 | True | UPF0234 protein Yitk |
| PGF_00047155 | 11.78 | 269 | 16 | 0.718 | 0.165 | True | Redox-sensing transcriptional repressor Rex |
| PGF_00405499 | 11.77 | 340 | 16 | 0.638 | 0.086 | True | YpfJ protein, zinc metalloprotease superfamily |
| PGF_02349208 | 11.76 | 230 | 16 | 0.776 | 0.020 | True | Hypothetical, related to broad specificity phosphatases COG0406 |
| PGF_00336478 | 11.76 | 222 | 16 | 0.789 | 0.064 | True | Two-component transcriptional response regulator PdtaR, LuxR family |
| PGF_01382519 | 11.75 | 215 | 16 | 0.802 | 0.043 | True | Ribosome hibernation promoting factor Hpf |
| PGF_06975895 | 11.72 | 280 | 16 | 0.700 | 0.082 | True | Metal-dependent hydrolases of the beta-lactamase superfamily III |
| PGF_08432396 | 11.69 | 230 | 16 | 0.771 | 0.088 | True | Nicotinate-nucleotide adenylyltransferase (EC 2.7.7.18) |
| PGF_00689961 | 11.68 | 211 | 16 | 0.804 | 0.056 | True | Guanylate kinase (EC 2.7.4.8) |
| PGF_00006023 | 11.68 | 313 | 16 | 0.660 | 0.013 | True | Type II/IV secretion system protein TadC, associated with Flp pilus assembly |
| PGF_04244475 | 11.65 | 195 | 16 | 0.835 | 0.020 | True | Adenylate kinase (EC 2.7.4.3) |
| PGF_00007085 | 11.64 | 416 | 16 | 0.571 | 0.111 | True | Galactokinase (EC 2.7.1.6) |
| PGF_00853991 | 11.63 | 321 | 16 | 0.649 | 0.168 | True | Diaminopimelate epimerase (EC 5.1.1.7) |
| PGF_02455692 | 11.62 | 245 | 16 | 0.742 | 0.056 | True | Cytidylate kinase (EC 2.7.4.25) |
| PGF_00048926 | 11.61 | 186 | 16 | 0.851 | 0.015 | True | Ribosome recycling factor |
| PGF_00000584 | 11.60 | 267 | 16 | 0.710 | 0.031 | True | Uncharacterized protien SCO1664 |
| PGF_00423533 | 11.55 | 323 | 16 | 0.643 | 0.066 | True | 4-diphosphocytidyl-2-C-methyl-D-erythritol kinase (EC 2.7.1.148) |
| PGF_00406805 | 11.54 | 306 | 16 | 0.660 | 0.113 | True | putative esterase/lipase |
| PGF_00025215 | 11.54 | 189 | 16 | 0.840 | 0.036 | True | Acetolactate synthase small subunit (EC 2.2.1.6) |
| PGF_00060585 | 11.54 | 256 | 16 | 0.721 | 0.080 | True | Transmembrane protein MT2276, clustered with lipoate gene |
| PGF_08228852 | 11.54 | 285 | 16 | 0.684 | 0.057 | True | Zinc ABC transporter, ATP-binding protein ZnuC |
| PGF_00060478 | 11.53 | 281 | 16 | 0.688 | 0.172 | True | Translation initiation factor 3 |
| PGF_00065286 | 11.52 | 213 | 16 | 0.789 | 0.045 | True | Uncharacterized protein Q1 colocalized with Q |
| PGF_00410606 | 11.52 | 234 | 16 | 0.753 | 0.067 | True | Uncharacterized protein SCO2312 |
| PGF_00002592 | 11.48 | 292 | 16 | 0.672 | 0.154 | True | FIG01964566: Predicted membrane protein, hemolysin III homolog |
| PGF_06111020 | 11.45 | 236 | 16 | 0.745 | 0.097 | True | Thymidylate kinase (EC 2.7.4.9) |
| PGF_06766299 | 11.44 | 492 | 16 | 0.516 | 0.192 | True | hypothetical protein |
| PGF_00984073 | 11.43 | 241 | 16 | 0.736 | 0.098 | True | Phosphate transport system regulatory protein PhoU |
| PGF_01447139 | 11.43 | 220 | 16 | 0.771 | 0.118 | True | ATP:Cob(I)alamin adenosyltransferase (EC 2.5.1.17) |
| PGF_07695531 | 11.42 | 143 | 16 | 0.955 | 0.006 | True | LSU ribosomal protein L11p (L12e) |
| PGF_10123167 | 11.42 | 189 | 16 | 0.831 | 0.000 | True | 2-amino-4-hydroxy-6-hydroxymethyldihydropteridine pyrophosphokinase (EC 2.7.6.3) |
| PGF_04788810 | 11.40 | 220 | 16 | 0.769 | 0.092 | True | Peptidyl-tRNA hydrolase (EC 3.1.1.29) |
| PGF_00049904 | 11.40 | 171 | 16 | 0.872 | 0.082 | True | SSU ribosomal protein S7p (S5e) |
| PGF_02334240 | 11.38 | 303 | 16 | 0.654 | 0.122 | True | Biotin--protein ligase (EC 6.3.4.9)(EC 6.3.4.10)(EC 6.3.4.11)(EC 6.3.4.15) |
| PGF_00016343 | 11.37 | 139 | 16 | 0.965 | 0.000 | True | LSU ribosomal protein L16p (L10e) |
| PGF_00033968 | 11.33 | 219 | 16 | 0.766 | 0.054 | True | Phosphoribosylglycinamide formyltransferase (EC 2.1.2.2) |
| PGF_00504549 | 11.25 | 279 | 16 | 0.674 | 0.198 | True | O-methyltransferase Rv1220c |
| PGF_00003506 | 11.23 | 250 | 16 | 0.710 | 0.044 | True | FIG137478: Hypothetical protein |
| PGF_09288314 | 11.22 | 331 | 16 | 0.617 | 0.228 | True | RNA methyltransferase, TrmH family |
| PGF_02792560 | 11.22 | 229 | 16 | 0.741 | 0.063 | True | tRNA threonylcarbamoyladenosine biosynthesis protein TsaB |
| PGF_00419702 | 11.20 | 191 | 16 | 0.811 | 0.053 | True | Crossover junction endodeoxyribonuclease RuvC (EC 3.1.22.4) |
| PGF_01187824 | 11.20 | 257 | 16 | 0.699 | 0.142 | True | Demethylmenaquinone methyltransferase (EC 2.1.1.163) |
| PGF_08181546 | 11.20 | 301 | 16 | 0.645 | 0.090 | True | Shikimate 5-dehydrogenase I alpha (EC 1.1.1.25) |
| PGF_01922063 | 11.19 | 202 | 16 | 0.787 | 0.121 | True | Inorganic pyrophosphatase (EC 3.6.1.1) |
| PGF_02150552 | 11.17 | 229 | 16 | 0.738 | 0.091 | True | Putative hydrolase in cluster with formaldehyde/S-nitrosomycothiol reductase MscR |
| PGF_00049909 | 11.14 | 181 | 16 | 0.828 | 0.075 | True | SSU ribosomal protein S9p (S16e) |
| PGF_00015543 | 11.12 | 250 | 16 | 0.704 | 0.101 | True | Iron-sulfur cluster regulator SufR |
| PGF_00049837 | 11.12 | 135 | 16 | 0.957 | 0.024 | True | SSU ribosomal protein S11p (S14e) |
| PGF_01959504 | 11.12 | 199 | 16 | 0.788 | 0.024 | True | DUF2017 domain-containing protein |
| PGF_00024692 | 11.11 | 290 | 16 | 0.652 | 0.224 | True | NADH-ubiquinone oxidoreductase chain E (EC 1.6.5.3) |
| PGF_06180597 | 11.09 | 147 | 16 | 0.915 | 0.000 | True | LSU ribosomal protein L13p (L13Ae) |
| PGF_00016342 | 11.08 | 147 | 16 | 0.914 | 0.006 | True | LSU ribosomal protein L15p (L27Ae) |
| PGF_06649360 | 11.07 | 289 | 16 | 0.651 | 0.140 | True | Segregation and condensation protein B |
| PGF_00807568 | 11.07 | 324 | 16 | 0.615 | 0.029 | True | FIG056164: rhomboid family serine protease |
| PGF_04232510 | 11.05 | 177 | 16 | 0.830 | 0.018 | True | Phospholipid-binding protein |
| PGF_00267378 | 11.04 | 153 | 16 | 0.892 | 0.016 | True | AIG2-like domain protein |
| PGF_00049906 | 10.99 | 139 | 16 | 0.932 | 0.005 | True | SSU ribosomal protein S8p (S15Ae) |
| PGF_03751076 | 10.98 | 245 | 16 | 0.701 | 0.141 | True | Riboflavin synthase eubacterial/eukaryotic (EC 2.5.1.9) |
| PGF_00045787 | 10.97 | 221 | 16 | 0.738 | 0.094 | True | Pyridoxal 5'-phosphate synthase (glutamine hydrolyzing), glutaminase subunit (EC 4.3.3.6) |
| PGF_01668012 | 10.96 | 297 | 16 | 0.636 | 0.097 | True | Sulfur carrier protein FdhD |
| PGF_00066124 | 10.93 | 218 | 16 | 0.740 | 0.128 | True | Pyrimidine operon regulatory protein PyrR |
| PGF_02031353 | 10.92 | 206 | 16 | 0.761 | 0.084 | True | FIG01269488: protein, clustered with ribosomal protein L32p |
| PGF_00049398 | 10.91 | 230 | 16 | 0.720 | 0.125 | True | Alternative RNA polymerase sigma factor SigE |
| PGF_07402119 | 10.90 | 255 | 16 | 0.682 | 0.052 | True | 6-phosphogluconolactonase (EC 3.1.1.31), eukaryotic type |
| PGF_08518355 | 10.90 | 283 | 16 | 0.648 | 0.062 | True | ATP synthase F0 sector subunit a (EC 3.6.3.14) |
| PGF_01761390 | 10.89 | 231 | 16 | 0.717 | 0.177 | True | NADH-ubiquinone oxidoreductase chain I (EC 1.6.5.3) |
| PGF_02976460 | 10.89 | 219 | 16 | 0.736 | 0.053 | True | Phosphatidylinositol phosphate synthase @ Archaetidylinositol phosphate synthase (EC 2.7.8.39) |
| PGF_00016340 | 10.88 | 122 | 16 | 0.985 | 0.000 | True | LSU ribosomal protein L14p (L23e) |
| PGF_05172785 | 10.86 | 183 | 16 | 0.803 | 0.029 | True | N5-carboxyaminoimidazole ribonucleotide mutase (EC 5.4.99.18) |
| PGF_02361937 | 10.86 | 230 | 16 | 0.716 | 0.065 | True | Hydrolase SCO5215, alpha/beta fold family |
| PGF_08843714 | 10.83 | 224 | 16 | 0.723 | 0.094 | True | Septum formation protein Maf |
| PGF_01722966 | 10.83 | 244 | 16 | 0.693 | 0.093 | True | Methyltransferase type 11 |
| PGF_07133621 | 10.82 | 202 | 16 | 0.762 | 0.056 | True | 16S rRNA (guanine(966)-N(2))-methyltransferase (EC 2.1.1.171) |
| PGF_03295678 | 10.79 | 232 | 16 | 0.708 | 0.103 | True | CDP-diacylglycerol--glycerol-3-phosphate 3-phosphatidyltransferase (EC 2.7.8.5) |
| PGF_12734426 | 10.78 | 348 | 16 | 0.578 | 0.136 | True | ATP/GTP-binding protein |
| PGF_00049840 | 10.78 | 125 | 16 | 0.964 | 0.008 | True | SSU ribosomal protein S13p (S18e) |
| PGF_00016358 | 10.73 | 130 | 16 | 0.941 | 0.015 | True | LSU ribosomal protein L20p |
| PGF_05296102 | 10.70 | 319 | 16 | 0.599 | 0.152 | True | Acetyl-CoA:Cys-GlcN-Ins acetyltransferase, mycothiol synthase MshD (EC 2.3.1.189) |
| PGF_00001849 | 10.69 | 205 | 16 | 0.747 | 0.160 | True | FIG01122152: hypothetical protein |
| PGF_00034972 | 10.69 | 189 | 16 | 0.777 | 0.032 | True | Adenine phosphoribosyltransferase (EC 2.4.2.7) |
| PGF_00422463 | 10.65 | 174 | 16 | 0.808 | 0.054 | True | DUF1794 |
| PGF_10546429 | 10.64 | 198 | 16 | 0.756 | 0.055 | True | ATP synthase F0 sector subunit b (EC 3.6.3.14) |
| PGF_12753784 | 10.63 | 264 | 16 | 0.654 | 0.211 | True | Transcriptional regulator SCO5170, AcrR family |
| PGF_07240617 | 10.62 | 278 | 16 | 0.637 | 0.146 | True | Uncharacterized AMP-finding protein SCO3041 |
| PGF_00048556 | 10.61 | 302 | 16 | 0.610 | 0.185 | True | Ribonuclease HII (EC 3.1.26.4) |
| PGF_00016919 | 10.59 | 277 | 16 | 0.637 | 0.150 | True | Leucyl/phenylalanyl-tRNA--protein transferase (EC 2.3.2.6) |
| PGF_00026362 | 10.57 | 247 | 16 | 0.673 | 0.152 | True | Nucleoside 5-triphosphatase RdgB (dHAPTP, dITP, XTP-specific) (EC 3.6.1.66) |
| PGF_10334950 | 10.57 | 234 | 16 | 0.691 | 0.073 | True | NADPH-dependent FMN reductase family protein |
| PGF_00419913 | 10.54 | 168 | 16 | 0.813 | 0.044 | True | Cyclic pyranopterin monophosphate synthase (EC 4.6.1.17) |
| PGF_06332624 | 10.54 | 258 | 16 | 0.656 | 0.056 | True | FIG215594: Membrane spanning protein |
| PGF_06941403 | 10.51 | 134 | 16 | 0.908 | 0.072 | True | SSU ribosomal protein S12p (S23e) |
| PGF_00985834 | 10.48 | 139 | 16 | 0.889 | 0.006 | True | Aspartate 1-decarboxylase (EC 4.1.1.11) |
| PGF_05732449 | 10.47 | 241 | 16 | 0.674 | 0.162 | True | Protein disulfide oxidoreductase |
| PGF_00000568 | 10.43 | 129 | 16 | 0.918 | 0.018 | True | FIG00820327: hypothetical protein |
| PGF_07668761 | 10.43 | 389 | 16 | 0.529 | 0.077 | True | Phosphate ABC transporter, substrate-binding protein PstS (TC 3.A.1.7.1) |
| PGF_00016368 | 10.43 | 152 | 16 | 0.846 | 0.053 | True | LSU ribosomal protein L22p (L17e) |
| PGF_10381714 | 10.40 | 208 | 16 | 0.721 | 0.138 | True | Orotate phosphoribosyltransferase (EC 2.4.2.10) |
| PGF_00048643 | 10.40 | 170 | 16 | 0.798 | 0.057 | True | Ribonucleotide reductase transcriptional regulator NrdR |
| PGF_01135236 | 10.35 | 200 | 16 | 0.732 | 0.012 | True | Dephospho-CoA kinase (EC 2.7.1.24) |
| PGF_00220548 | 10.34 | 166 | 16 | 0.802 | 0.056 | True | Thiol peroxidase, Bcp-type (EC 1.11.1.15) |
| PGF_00045833 | 10.33 | 236 | 16 | 0.673 | 0.089 | True | Pyridoxamine 5'-phosphate oxidase (EC 1.4.3.5) |
| PGF_02156631 | 10.32 | 195 | 16 | 0.739 | 0.075 | True | Arginine pathway regulatory protein ArgR, repressor of arg regulon |
| PGF_01420802 | 10.31 | 143 | 16 | 0.863 | 0.007 | True | Transcriptional regulator MraZ |
| PGF_03021263 | 10.26 | 185 | 16 | 0.754 | 0.122 | True | SepF, FtsZ-interacting protein related to cell division |
| PGF_09402727 | 10.22 | 272 | 16 | 0.620 | 0.039 | True | Molybdenum ABC transporter permease protein ModB |
| PGF_00016452 | 10.22 | 149 | 16 | 0.837 | 0.006 | True | LSU ribosomal protein L9p |
| PGF_05952248 | 10.22 | 404 | 16 | 0.508 | 0.183 | True | putative secreted hydrolase |
| PGF_04618589 | 10.21 | 177 | 16 | 0.767 | 0.075 | True | Glycogen accumulation regulator GarA |
| PGF_02565697 | 10.18 | 206 | 16 | 0.709 | 0.068 | True | hypothetical protein |
| PGF_00662997 | 10.12 | 287 | 16 | 0.598 | 0.108 | True | Cobalamin synthase (EC 2.7.8.26) |
| PGF_00416525 | 10.11 | 176 | 16 | 0.762 | 0.056 | True | Carbonic anhydrase, beta class (EC 4.2.1.1) |
| PGF_06626131 | 10.11 | 182 | 16 | 0.749 | 0.101 | True | 2-C-methyl-D-erythritol 2,4-cyclodiphosphate synthase (EC 4.6.1.12) |
| PGF_01724713 | 10.10 | 203 | 16 | 0.709 | 0.154 | True | Ribose-5-phosphate isomerase B (EC 5.3.1.6) |
| PGF_06000821 | 10.08 | 220 | 16 | 0.679 | 0.109 | True | 16S rRNA processing protein RimM |
| PGF_00049828 | 10.06 | 102 | 16 | 0.997 | 0.000 | True | SSU ribosomal protein S10p (S20e) |
| PGF_04762552 | 10.03 | 166 | 16 | 0.779 | 0.026 | True | Phosphopantetheine adenylyltransferase (EC 2.7.7.3) |
| PGF_00016445 | 10.00 | 137 | 16 | 0.854 | 0.052 | True | LSU ribosomal protein L7p/L12p (P1/P2) |
| PGF_00041788 | 9.99 | 168 | 16 | 0.771 | 0.083 | True | Putative iron-sulfur cluster assembly scaffold protein for SUF system, SufE2 |
| PGF_00020361 | 9.97 | 175 | 16 | 0.754 | 0.111 | True | Metal-dependent hydrolase YbeY, involved in rRNA and/or ribosome maturation and assembly |
| PGF_00426932 | 9.97 | 166 | 16 | 0.774 | 0.041 | True | 6,7-dimethyl-8-ribityllumazine synthase (EC 2.5.1.78) |
| PGF_00016353 | 9.94 | 129 | 16 | 0.875 | 0.015 | True | LSU ribosomal protein L18p (L5e) |
| PGF_10488938 | 9.90 | 165 | 16 | 0.771 | 0.061 | True | Putative pre-16S rRNA nuclease YqgF |
| PGF_00178044 | 9.89 | 209 | 16 | 0.684 | 0.168 | True | SSU ribosomal protein S16p |
| PGF_02569889 | 9.88 | 249 | 16 | 0.626 | 0.178 | True | hypothetical protein |
| PGF_02359675 | 9.84 | 135 | 16 | 0.847 | 0.007 | True | Uncharacterized protein SCO1141 |
| PGF_00016346 | 9.82 | 260 | 16 | 0.609 | 0.221 | True | LSU ribosomal protein L17p |
| PGF_00022507 | 9.77 | 191 | 16 | 0.707 | 0.146 | True | Molybdopterin adenylyltransferase (EC 2.7.7.75) |
| PGF_04710902 | 9.74 | 263 | 16 | 0.600 | 0.065 | True | Lactam utilization protein LamB |
| PGF_00426285 | 9.74 | 200 | 16 | 0.689 | 0.123 | True | FIG004853: possible toxin to DivIC |
| PGF_05126349 | 9.72 | 214 | 16 | 0.664 | 0.089 | True | CblZ, a non-orthologous displasment for Alpha-ribazole-5'-phosphate phosphatase |
| PGF_00000494 | 9.71 | 209 | 16 | 0.672 | 0.148 | True | FIG00816212: Putative membrane protein |
| PGF_04845029 | 9.71 | 117 | 16 | 0.898 | 0.008 | True | LSU ribosomal protein L19p |
| PGF_00038579 | 9.66 | 102 | 16 | 0.956 | 0.001 | True | Protein often found in Actinomycetes clustered with signal peptidase and/or RNaseHII |
| PGF_09008105 | 9.65 | 135 | 16 | 0.831 | 0.065 | True | Chorismate mutase II (EC 5.4.99.5) |
| PGF_00065292 | 9.61 | 208 | 16 | 0.666 | 0.146 | True | Uncharacterized protein Rv0487/MT0505 clustered with mycothiol biosynthesis gene |
| PGF_02898874 | 9.60 | 158 | 16 | 0.764 | 0.117 | True | Putative response regulator |
| PGF_00420001 | 9.59 | 176 | 16 | 0.723 | 0.151 | True | CysO-cysteine peptidase |
| PGF_04198961 | 9.59 | 217 | 16 | 0.651 | 0.118 | True | DNA-3-methyladenine glycosylase II (EC 3.2.2.21) |
| PGF_00021018 | 9.55 | 172 | 16 | 0.728 | 0.116 | True | Methylmalonyl-CoA epimerase (EC 5.1.99.1) @ Ethylmalonyl-CoA epimerase |
| PGF_01674551 | 9.55 | 129 | 16 | 0.840 | 0.008 | True | Glycine cleavage system H protein |
| PGF_05608844 | 9.53 | 110 | 16 | 0.909 | 0.000 | True | Small basic protein Sbp |
| PGF_02838109 | 9.53 | 191 | 16 | 0.689 | 0.118 | True | Shikimate kinase I (EC 2.7.1.71) |
| PGF_03518570 | 9.51 | 143 | 16 | 0.796 | 0.016 | True | Nucleoside diphosphate kinase (EC 2.7.4.6) |
| PGF_04457297 | 9.50 | 208 | 16 | 0.659 | 0.042 | True | 5-formyltetrahydrofolate cyclo-ligase (EC 6.3.3.2) |
| PGF_00016377 | 9.50 | 124 | 16 | 0.853 | 0.052 | True | LSU ribosomal protein L24p (L26e) |
| PGF_03990071 | 9.49 | 101 | 16 | 0.944 | 0.001 | True | LSU ribosomal protein L23p (L23Ae) |
| PGF_08197987 | 9.45 | 147 | 16 | 0.780 | 0.173 | True | Iron-sulfur cluster insertion protein SCO2161 |
| PGF_07919746 | 9.45 | 135 | 16 | 0.813 | 0.013 | True | ATP synthase epsilon chain (EC 3.6.3.14) |
| PGF_00400287 | 9.44 | 108 | 16 | 0.908 | 0.011 | True | integration host factor |
| PGF_10450086 | 9.36 | 143 | 16 | 0.783 | 0.049 | True | Transcription termination protein NusB |
| PGF_05357708 | 9.34 | 221 | 16 | 0.628 | 0.153 | True | DNA-3-methyladenine glycosylase (EC 3.2.2.20) |
| PGF_01737397 | 9.34 | 491 | 16 | 0.422 | 0.398 | True | FIG00031715: Predicted metal-dependent phosphoesterases (PHP family) |
| PGF_00036456 | 9.33 | 200 | 16 | 0.660 | 0.216 | True | Predicted transcriptional regulator of sulfate adenylyltransferase, Rrf2 family |
| PGF_00418586 | 9.33 | 142 | 16 | 0.783 | 0.072 | True | Cold shock protein of CSP family => SCO4325 |
| PGF_00049860 | 9.33 | 93 | 16 | 0.967 | 0.000 | True | SSU ribosomal protein S19p (S15e) |
| PGF_06661068 | 9.29 | 135 | 16 | 0.800 | 0.076 | True | PaaD-like protein (DUF59) involved in Fe-S cluster assembly |
| PGF_07275226 | 9.25 | 313 | 16 | 0.523 | 0.211 | True | Competence protein F homolog, phosphoribosyltransferase domain; protein YhgH required for utilization of DNA as sole source of carbon and energy |
| PGF_00049847 | 9.24 | 91 | 16 | 0.968 | 0.000 | True | SSU ribosomal protein S15p (S13e) |
| PGF_02790700 | 9.21 | 189 | 16 | 0.670 | 0.072 | True | Adenosylcobinamide kinase (EC 2.7.1.156) / Adenosylcobinamide-phosphate guanylyltransferase (EC 2.7.7.62) |
| PGF_00417577 | 9.21 | 156 | 16 | 0.737 | 0.073 | True | 3-dehydroquinate dehydratase II (EC 4.2.1.10) |
| PGF_07687979 | 9.20 | 226 | 16 | 0.612 | 0.160 | True | Zn-ribbon-containing, possibly RNA-binding protein and truncated derivatives |
| PGF_03642879 | 9.19 | 138 | 16 | 0.783 | 0.038 | True | Cytochrome c oxidase polypeptide IV (EC 1.9.3.1) |
| PGF_07824136 | 9.19 | 437 | 16 | 0.439 | 0.316 | True | Late competence protein ComEA, DNA receptor |
| PGF_01430214 | 9.18 | 197 | 16 | 0.654 | 0.189 | True | Deoxyuridine 5'-triphosphate nucleotidohydrolase (EC 3.6.1.23) |
| PGF_05906360 | 9.15 | 112 | 16 | 0.864 | 0.066 | True | DNA-directed RNA polymerase omega subunit (EC 2.7.7.6) |
| PGF_09452281 | 9.12 | 201 | 16 | 0.643 | 0.184 | True | Ribosome-binding factor A |
| PGF_01682834 | 9.07 | 202 | 16 | 0.638 | 0.176 | True | Bacterial ribosome SSU maturation protein RimP |
| PGF_06368851 | 9.07 | 327 | 16 | 0.502 | 0.285 | True | DNA-binding protein HU / low-complexity, AKP-rich domain |
| PGF_03681316 | 9.05 | 136 | 16 | 0.776 | 0.074 | True | Hemoglobin-like protein HbO |
| PGF_07836795 | 9.02 | 99 | 16 | 0.906 | 0.000 | True | Aspartyl-tRNA(Asn) amidotransferase subunit C (EC 6.3.5.6) @ Glutamyl-tRNA(Gln) amidotransferase subunit C (EC 6.3.5.7) |
| PGF_09581668 | 8.99 | 98 | 16 | 0.909 | 0.009 | True | Heat shock protein 10 kDa family chaperone GroES |
| PGF_00066839 | 8.99 | 83 | 16 | 0.987 | 0.000 | True | WhiB-like transcription regulator |
| PGF_00257320 | 8.99 | 275 | 16 | 0.542 | 0.355 | True | hypothetical protein |
| PGF_00022550 | 8.96 | 170 | 16 | 0.687 | 0.159 | True | Molybdopterin synthase catalytic subunit MoaE (EC 2.8.1.12) |
| PGF_08749001 | 8.96 | 104 | 16 | 0.878 | 0.058 | True | NADH-ubiquinone oxidoreductase chain K (EC 1.6.5.3) |
| PGF_00003179 | 8.96 | 195 | 16 | 0.641 | 0.117 | True | FIG049476: HIT family protein |
| PGF_04978890 | 8.94 | 102 | 16 | 0.885 | 0.025 | True | LSU ribosomal protein L21p |
| PGF_00049901 | 8.81 | 100 | 16 | 0.881 | 0.038 | True | SSU ribosomal protein S6p |
| PGF_02776739 | 8.73 | 127 | 16 | 0.774 | 0.052 | True | Phosphoribosyl-AMP cyclohydrolase (EC 3.5.4.19) |
| PGF_00012430 | 8.69 | 155 | 16 | 0.698 | 0.094 | True | HspR, transcriptional repressor of DnaK operon |
| PGF_00016385 | 8.68 | 84 | 16 | 0.947 | 0.000 | True | LSU ribosomal protein L27p |
| PGF_03277153 | 8.65 | 104 | 16 | 0.848 | 0.037 | True | Sulfur metabolism protein SseC |
| PGF_10548652 | 8.60 | 121 | 16 | 0.782 | 0.101 | True | GroES-like protein in Actinomycetes |
| PGF_01979294 | 8.50 | 147 | 16 | 0.701 | 0.146 | True | Dihydroneopterin aldolase (EC 4.1.2.25) |
| PGF_00053952 | 8.49 | 86 | 16 | 0.915 | 0.015 | True | Sporulation regulatory protein WhiB |
| PGF_00049854 | 8.47 | 94 | 16 | 0.874 | 0.028 | True | SSU ribosomal protein S17p (S11e) |
| PGF_00057353 | 8.46 | 129 | 16 | 0.745 | 0.144 | True | Anti-sigma factor antagonist BldG |
| PGF_00435430 | 8.46 | 147 | 16 | 0.698 | 0.171 | True | Ribosomal silencing factor RsfA |
| PGF_04388239 | 8.35 | 80 | 16 | 0.934 | 0.001 | True | KH domain RNA binding protein YlqC |
| PGF_00960330 | 8.33 | 105 | 16 | 0.813 | 0.066 | True | ATP-dependent Clp protease adaptor protein ClpS |
| PGF_08837849 | 8.31 | 167 | 16 | 0.643 | 0.148 | True | Cytidine deaminase (EC 3.5.4.5) |
| PGF_00013272 | 8.23 | 173 | 16 | 0.626 | 0.166 | True | Peptidyl-tRNA hydrolase ArfB (EC 3.1.1.29) |
| PGF_02454577 | 8.17 | 86 | 16 | 0.881 | 0.000 | True | SSU ribosomal protein S20p |
| PGF_00059852 | 8.14 | 127 | 16 | 0.722 | 0.109 | True | Transcriptional regulator, WhiB family |
| PGF_10556954 | 8.13 | 125 | 16 | 0.727 | 0.040 | True | hypothetical protein |
| PGF_01172431 | 8.07 | 96 | 16 | 0.823 | 0.088 | True | Phosphoribosyl-ATP pyrophosphatase (EC 3.6.1.31) |
| PGF_04289125 | 8.00 | 149 | 16 | 0.656 | 0.094 | True | Molybdate-binding domain of ModE |
| PGF_00012152 | 8.00 | 117 | 16 | 0.740 | 0.071 | True | Histone protein Lsr2 |
| PGF_04097953 | 7.99 | 131 | 16 | 0.699 | 0.125 | True | Putative oxidoreductase |
| PGF_04972523 | 7.96 | 125 | 16 | 0.712 | 0.089 | True | Ferredoxin, 2Fe-2S |
| PGF_03751823 | 7.95 | 321 | 16 | 0.443 | 0.314 | True | Heat shock protein GrpE |
| PGF_00055165 | 7.88 | 102 | 16 | 0.780 | 0.098 | True | Sulfur/cysteine carrier protein CysO |
| PGF_00060472 | 7.86 | 111 | 16 | 0.746 | 0.082 | True | Antitoxin HigA |
| PGF_02899131 | 7.81 | 83 | 16 | 0.857 | 0.035 | True | LSU ribosomal protein L29p (L35e) |
| PGF_00934722 | 7.75 | 79 | 16 | 0.872 | 0.009 | True | Protein translocase membrane subunit SecG |
| PGF_00049842 | 7.71 | 61 | 16 | 0.987 | 0.000 | True | SSU ribosomal protein S14p (S29e) @ SSU ribosomal protein S14p (S29e), zinc-dependent |
| PGF_00016387 | 7.65 | 61 | 16 | 0.979 | 0.000 | True | LSU ribosomal protein L28p @ LSU ribosomal protein L28p, zinc-dependent |
| PGF_00003233 | 7.56 | 133 | 16 | 0.655 | 0.133 | True | FIG059443: hypothetical protein |
| PGF_05032476 | 7.55 | 127 | 16 | 0.670 | 0.106 | True | Uncharacterized protein MSMEG_5817 |
| PGF_03654614 | 7.43 | 135 | 16 | 0.640 | 0.235 | True | hypothetical protein |
| PGF_02501663 | 7.38 | 154 | 16 | 0.594 | 0.140 | True | UPF0102 protein YraN |
| PGF_00016395 | 7.29 | 60 | 16 | 0.941 | 0.000 | True | LSU ribosomal protein L30p (L7e) |
| PGF_07957805 | 7.26 | 94 | 16 | 0.748 | 0.056 | True | Phosphoribosylformylglycinamidine synthase, PurS subunit (EC 6.3.5.3) |
| PGF_03538342 | 7.24 | 124 | 16 | 0.651 | 0.222 | True | Transcriptional regulator, WhiB family |
| PGF_09669766 | 7.22 | 86 | 16 | 0.778 | 0.025 | True | Protein translocase subunit SecE |
| PGF_03038172 | 7.16 | 86 | 16 | 0.772 | 0.052 | True | Mycoredoxin (EC 1.20.4.3) |
| PGF_07092827 | 7.13 | 89 | 16 | 0.756 | 0.077 | True | Excisionase/Xis, DNA-binding |
| PGF_06609275 | 7.09 | 56 | 16 | 0.948 | 0.000 | True | LSU ribosomal protein L33p @ LSU ribosomal protein L33p, zinc-dependent |
| PGF_04946791 | 7.06 | 85 | 16 | 0.766 | 0.022 | True | hypothetical protein |
| PGF_03739865 | 7.04 | 81 | 16 | 0.783 | 0.127 | True | Dodecin, a flavin storage/sequestration protein |
| PGF_05859499 | 7.01 | 134 | 16 | 0.606 | 0.137 | True | Uncharacterized protein SCO2538 |
| PGF_06060005 | 7.00 | 71 | 16 | 0.831 | 0.041 | True | Excisionase-like protein SCO3328 |
| PGF_02572755 | 6.96 | 309 | 16 | 0.396 | 0.383 | True | hypothetical protein |
| PGF_08210457 | 6.90 | 67 | 16 | 0.843 | 0.028 | True | Prokaryotic ubiquitin-like protein Pup |
| PGF_00046192 | 6.89 | 233 | 16 | 0.452 | 0.339 | True | RNA 3'-terminal phosphate cyclase (EC 6.5.1.4) |
| PGF_10282774 | 6.50 | 60 | 16 | 0.839 | 0.125 | True | Uncharacterized protein Noca_1520 |
| PGF_05770273 | 6.36 | 108 | 16 | 0.612 | 0.299 | True | LSU ribosomal protein L31p @ LSU ribosomal protein L31p, zinc-dependent |
| PGF_07550057 | 6.04 | 64 | 16 | 0.755 | 0.107 | True | FIG002473: Protein YcaR in KDO2-Lipid A biosynthesis cluster |
| PGF_10315700 | 6.03 | 69 | 16 | 0.727 | 0.097 | True | Uncharacterized protein SCO4088 |
| PGF_01787293 | 6.03 | 113 | 16 | 0.567 | 0.298 | True | Uncharacterized protein Sros_7085 |
| PGF_00016424 | 5.94 | 37 | 16 | 0.977 | 0.000 | True | LSU ribosomal protein L36p @ LSU ribosomal protein L36p, zinc-dependent |
| PGF_12924000 | 5.94 | 80 | 16 | 0.664 | 0.291 | True | Uncharacterized protein MSMEG_5081 |
| PGF_05076104 | 5.85 | 89 | 16 | 0.620 | 0.195 | True | Exodeoxyribonuclease VII small subunit (EC 3.1.11.6) |
| PGF_00980320 | 5.66 | 88 | 16 | 0.604 | 0.212 | True | hypothetical protein |
| PGF_10083078 | 4.14 | 63 | 16 | 0.521 | 0.428 | True | hypothetical protein |
| PGF_01205865 | 3.44 | 317 | 16 | 0.193 | 0.727 | False | hypothetical protein |
