## Supplementary material for "Biotechnological potential of aromatic compounds–utilizing bacteria from Brazilian caves, including a novel cave *Nocardioides sp*. SF1": Table S2

**Table S2 – Single-copy genes used to phylogenomic of 60 *Nocardioides* species**

| **PGFam** | **Align. Score** | **Align. Length** | **Num Seqs** | **Mean Sqr Freq** | **Prop Gaps** | **Used In Analysis** | **Product** |
| --- | --- | --- | --- | --- | --- | --- | --- |
| PGF_08854233 | 25.96 | 1050 | 60 | 0.801 | 0.080 | True | Ribonucleotide reductase of class II (coenzyme B12-dependent) (EC 1.17.4.1) |
| PGF_00950554 | 23.41 | 723 | 60 | 0.871 | 0.020 | True | Excinuclease ABC subunit B |
| PGF_05500127 | 23.14 | 935 | 60 | 0.757 | 0.066 | True | Valyl-tRNA synthetase (EC 6.1.1.9) |
| PGF_03272313 | 23.07 | 1030 | 60 | 0.719 | 0.121 | True | DNA gyrase subunit A (EC 5.99.1.3) |
| PGF_10049811 | 22.75 | 1108 | 60 | 0.683 | 0.148 | True | Protein translocase subunit SecA |
| PGF_00426726 | 22.66 | 1010 | 60 | 0.713 | 0.078 | True | FIG005666: putative helicase |
| PGF_01053024 | 22.28 | 1065 | 60 | 0.683 | 0.071 | True | Glutamine synthetase adenylyl-L-tyrosine phosphorylase (EC 2.7.7.89) / Glutamate-ammonia-ligase adenylyltransferase (EC 2.7.7.42) |
| PGF_03752158 | 21.52 | 710 | 60 | 0.807 | 0.076 | True | FIG092679: Fe-S oxidoreductase |
| PGF_00071558 | 20.73 | 614 | 60 | 0.837 | 0.052 | True | Bacterial proteasome-activating AAA-ATPase (PAN) |
| PGF_00426115 | 20.17 | 675 | 60 | 0.777 | 0.060 | True | 1-deoxy-D-xylulose 5-phosphate synthase (EC 2.2.1.7) |
| PGF_06703483 | 20.15 | 835 | 60 | 0.697 | 0.143 | True | DNA gyrase subunit B (EC 5.99.1.3) |
| PGF_00015514 | 20.15 | 487 | 60 | 0.913 | 0.029 | True | Iron-sulfur cluster assembly protein SufB |
| PGF_00038970 | 19.74 | 453 | 60 | 0.927 | 0.000 | True | Pup ligase PafA, possible component of postulated heterodimer PafA-PafA' |
| PGF_05195470 | 19.71 | 834 | 60 | 0.683 | 0.130 | True | Polyphosphate kinase (EC 2.7.4.1) |
| PGF_07904640 | 19.32 | 535 | 60 | 0.835 | 0.049 | True | XRE-family DNA-binding domain / UDP-N-acetylglucosamine 1-carboxyvinyltransferase (EC 2.5.1.7) |
| PGF_00423472 | 18.97 | 758 | 60 | 0.689 | 0.119 | True | DinG family ATP-dependent helicase YoaA |
| PGF_00038969 | 18.85 | 507 | 60 | 0.837 | 0.007 | True | Pup ligase PafA' paralog, possible component of postulated heterodimer PafA-PafA' |
| PGF_00052238 | 18.62 | 585 | 60 | 0.770 | 0.103 | True | Signal recognition particle protein Ffh |
| PGF_00016338 | 18.52 | 433 | 60 | 0.890 | 0.015 | True | ATP-dependent Clp protease ATP-binding subunit ClpX |
| PGF_00066286 | 18.50 | 618 | 60 | 0.744 | 0.104 | True | Uroporphyrinogen-III methyltransferase (EC 2.1.1.107) / Uroporphyrinogen-III synthase (EC 4.2.1.75) |
| PGF_00419420 | 18.42 | 503 | 60 | 0.821 | 0.064 | True | 3-isopropylmalate dehydratase large subunit (EC 4.2.1.33) |
| PGF_00067554 | 18.37 | 524 | 60 | 0.803 | 0.048 | True | Aspartyl-tRNA(Asn) amidotransferase subunit B (EC 6.3.5.6) @ Glutamyl-tRNA(Gln) amidotransferase subunit B (EC 6.3.5.7) |
| PGF_06525160 | 18.36 | 787 | 60 | 0.655 | 0.137 | True | NAD synthetase (EC 6.3.1.5) / Glutamine amidotransferase chain of NAD synthetase |
| PGF_00033095 | 18.31 | 1080 | 60 | 0.557 | 0.230 | True | Phenylalanyl-tRNA synthetase beta chain (EC 6.1.1.20) |
| PGF_10440725 | 18.27 | 513 | 60 | 0.806 | 0.048 | True | Glutamate synthase [NADPH] small chain (EC 1.4.1.13) |
| PGF_00016824 | 18.23 | 746 | 60 | 0.667 | 0.084 | True | ATP-dependent DNA helicase UvrD/PcrA, actinomycete paralog |
| PGF_02226715 | 18.17 | 807 | 60 | 0.640 | 0.083 | True | ATP-dependent DNA helicase RecG (EC 3.6.4.12) |
| PGF_00425021 | 18.08 | 1490 | 60 | 0.468 | 0.252 | True | Exodeoxyribonuclease V gamma chain (EC 3.1.11.5) |
| PGF_00004392 | 18.08 | 634 | 60 | 0.718 | 0.098 | True | Ferredoxin--sulfite reductase, actinobacterial type (EC 1.8.7.1) |
| PGF_00060414 | 18.03 | 954 | 60 | 0.584 | 0.339 | True | Translation elongation factor LepA |
| PGF_03609651 | 18.00 | 646 | 60 | 0.708 | 0.074 | True | Methionyl-tRNA synthetase (EC 6.1.1.10) |
| PGF_00017545 | 17.79 | 659 | 60 | 0.693 | 0.093 | True | Long-chain-fatty-acid--CoA ligase (EC 6.2.1.3) @ Long-chain fatty-acid-CoA ligase (EC 6.2.1.3), Mycobacterial subgroup FadD15 |
| PGF_12694106 | 17.77 | 654 | 60 | 0.695 | 0.114 | True | 2-isopropylmalate synthase (EC 2.3.3.13) |
| PGF_00007024 | 17.69 | 479 | 60 | 0.808 | 0.055 | True | GTP-binding protein EngA |
| PGF_04425336 | 17.52 | 473 | 60 | 0.806 | 0.075 | True | NADH-ubiquinone oxidoreductase chain F (EC 1.6.5.3) |
| PGF_05581732 | 17.49 | 450 | 60 | 0.824 | 0.040 | True | Protein translocase subunit SecY |
| PGF_00422271 | 17.43 | 342 | 60 | 0.942 | 0.012 | True | DNA-directed RNA polymerase alpha subunit (EC 2.7.7.6) |
| PGF_02516909 | 17.40 | 434 | 60 | 0.835 | 0.020 | True | Enolase (EC 4.2.1.11) |
| PGF_00037269 | 17.16 | 776 | 60 | 0.616 | 0.181 | True | 2-Amino-2-deoxy-isochorismate synthase (EC 4.1.3.-) |
| PGF_00417840 | 17.12 | 405 | 60 | 0.851 | 0.031 | True | Chorismate synthase (EC 4.2.3.5) |
| PGF_02029783 | 17.06 | 577 | 60 | 0.710 | 0.100 | True | GTP-binding protein Obg |
| PGF_00007028 | 16.83 | 603 | 60 | 0.686 | 0.157 | True | Ribosome LSU-associated GTP-binding protein HflX |
| PGF_00416129 | 16.79 | 826 | 60 | 0.584 | 0.312 | True | CTP synthase (EC 6.3.4.2) |
| PGF_00420081 | 16.79 | 420 | 60 | 0.819 | 0.053 | True | Cysteine synthesis adenylyltransferase/sulfurtransferase |
| PGF_00425738 | 16.76 | 329 | 60 | 0.924 | 0.003 | True | Sporulation transcription regulator WhiA |
| PGF_00426236 | 16.69 | 439 | 60 | 0.796 | 0.125 | True | (E)-4-hydroxy-3-methylbut-2-enyl-diphosphate synthase (flavodoxin) (EC 1.17.7.3) |
| PGF_00071194 | 16.65 | 517 | 60 | 0.732 | 0.128 | True | 2-keto-3-deoxy-D-arabino-heptulosonate-7-phosphate synthase II (EC 2.5.1.54) |
| PGF_00047078 | 16.61 | 380 | 60 | 0.852 | 0.077 | True | RecA protein |
| PGF_00006461 | 16.56 | 494 | 60 | 0.745 | 0.062 | True | Fumarate hydratase class II (EC 4.2.1.2) |
| PGF_00008075 | 16.53 | 397 | 60 | 0.830 | 0.035 | True | Glutamate N-acetyltransferase (EC 2.3.1.35) @ N-acetylglutamate synthase (EC 2.3.1.1) |
| PGF_05935795 | 16.29 | 513 | 60 | 0.719 | 0.109 | True | Replication-associated recombination protein RarA |
| PGF_03202156 | 16.27 | 427 | 60 | 0.787 | 0.064 | True | S-adenosylmethionine synthetase (EC 2.5.1.6) |
| PGF_02430085 | 16.16 | 418 | 60 | 0.790 | 0.041 | True | Acyl-CoA dehydrogenase (EC 1.3.8.1), Mycobacterial subgroup FadE23 |
| PGF_07583562 | 16.11 | 443 | 60 | 0.765 | 0.060 | True | Cysteine desulfurase (EC 2.8.1.7) => SufS |
| PGF_00407244 | 16.10 | 1052 | 60 | 0.496 | 0.324 | True | putative helicase regulator |
| PGF_00421347 | 16.10 | 373 | 60 | 0.833 | 0.035 | True | DNA integrity scanning protein DisA |
| PGF_00420080 | 16.05 | 325 | 60 | 0.890 | 0.025 | True | Cysteine synthase, CysO-dependent |
| PGF_00008337 | 16.03 | 543 | 60 | 0.688 | 0.098 | True | Glutamyl-tRNA synthetase (EC 6.1.1.17) @ Glutamyl-tRNA(Gln) synthetase (EC 6.1.1.24) |
| PGF_06935032 | 15.98 | 446 | 60 | 0.757 | 0.041 | True | Adenylosuccinate synthetase (EC 6.3.4.4) |
| PGF_10143857 | 15.95 | 345 | 60 | 0.858 | 0.005 | True | Fructose-bisphosphate aldolase class II (EC 4.1.2.13) |
| PGF_00421792 | 15.82 | 536 | 60 | 0.683 | 0.124 | True | DNA repair protein RadA |
| PGF_00016393 | 15.81 | 278 | 60 | 0.948 | 0.000 | True | LSU ribosomal protein L2p (L8e) |
| PGF_00876106 | 15.78 | 629 | 60 | 0.629 | 0.220 | True | Chromosomal replication initiator protein DnaA |
| PGF_02278006 | 15.67 | 433 | 60 | 0.753 | 0.091 | True | Quinolinate synthetase (EC 2.5.1.72) |
| PGF_08398205 | 15.67 | 639 | 60 | 0.620 | 0.236 | True | CCA tRNA nucleotidyltransferase (EC 2.7.7.72) |
| PGF_05950073 | 15.61 | 430 | 60 | 0.753 | 0.091 | True | Cystathionine gamma-lyase (EC 4.4.1.1) |
| PGF_05387084 | 15.54 | 461 | 60 | 0.724 | 0.091 | True | Gamma-glutamyl phosphate reductase (EC 1.2.1.41) |
| PGF_02797400 | 15.51 | 555 | 60 | 0.658 | 0.149 | True | Cysteinyl-tRNA synthetase (EC 6.1.1.16) |
| PGF_00030640 | 15.49 | 363 | 60 | 0.813 | 0.021 | True | Peptide chain release factor 1 |
| PGF_00062027 | 15.46 | 492 | 60 | 0.697 | 0.140 | True | Tryptophan synthase beta chain (EC 4.2.1.20) |
| PGF_00618776 | 15.36 | 451 | 60 | 0.723 | 0.154 | True | 23S rRNA (adenine(2503)-C(2))-methyltransferase @ tRNA (adenine(37)-C(2))-methyltransferase (EC 2.1.1.192) |
| PGF_00024274 | 15.35 | 438 | 60 | 0.734 | 0.114 | True | N5-carboxyaminoimidazole ribonucleotide synthase (EC 6.3.4.18) |
| PGF_00016074 | 15.32 | 441 | 60 | 0.730 | 0.066 | True | L-cysteine:1D-myo-inosityl 2-amino-2-deoxy-alpha-D-glucopyranoside ligase MshC (EC 6.3.1.13) |
| PGF_00033987 | 15.31 | 393 | 60 | 0.772 | 0.048 | True | Phosphoserine aminotransferase (EC 2.6.1.52) |
| PGF_06473395 | 15.29 | 420 | 60 | 0.746 | 0.097 | True | DNA polymerase III beta subunit (EC 2.7.7.7) |
| PGF_05049118 | 15.23 | 409 | 60 | 0.753 | 0.114 | True | Holliday junction ATP-dependent DNA helicase RuvB (EC 3.6.4.12) |
| PGF_05760069 | 15.18 | 492 | 60 | 0.684 | 0.110 | True | Histidinol dehydrogenase (EC 1.1.1.23) |
| PGF_00423429 | 15.05 | 676 | 60 | 0.579 | 0.182 | True | Dihydroxyacetone kinase-like protein, phosphatase domain / Dihydroxyacetone kinase-like protein, kinase domain |
| PGF_00294865 | 15.00 | 631 | 60 | 0.597 | 0.157 | True | Two component system sensor histidine kinase MtrB |
| PGF_03219041 | 14.82 | 442 | 60 | 0.705 | 0.065 | True | putative phospholipase D family protein |
| PGF_00046352 | 14.77 | 359 | 60 | 0.780 | 0.103 | True | RNA polymerase sigma factor SigB |
| PGF_08364774 | 14.63 | 363 | 60 | 0.768 | 0.070 | True | Fructose-1,6-bisphosphatase, GlpX type (EC 3.1.3.11) |
| PGF_03316046 | 14.61 | 401 | 60 | 0.729 | 0.103 | True | Threonine synthase (EC 4.2.3.1) |
| PGF_00033954 | 14.41 | 395 | 60 | 0.725 | 0.090 | True | Phosphoribosylformylglycinamidine cyclo-ligase (EC 6.3.3.1) |
| PGF_00067188 | 14.31 | 385 | 60 | 0.729 | 0.097 | True | Aspartate-semialdehyde dehydrogenase (EC 1.2.1.11) |
| PGF_00489714 | 14.31 | 359 | 60 | 0.755 | 0.075 | True | Porphobilinogen synthase (EC 4.2.1.24) |
| PGF_00042294 | 14.26 | 592 | 60 | 0.586 | 0.169 | True | Putative methyltransferase SCO3545 |
| PGF_08173994 | 14.26 | 514 | 60 | 0.629 | 0.174 | True | Glycosyltransferase SCO2318 |
| PGF_04991657 | 14.19 | 454 | 60 | 0.666 | 0.106 | True | Oxygen-independent coproporphyrinogen-III oxidase-like protein YggW |
| PGF_00015259 | 14.18 | 333 | 60 | 0.777 | 0.081 | True | ATP synthase gamma chain (EC 3.6.3.14) |
| PGF_05026985 | 14.08 | 358 | 60 | 0.744 | 0.104 | True | Deoxyribose-phosphate aldolase (EC 4.1.2.4) |
| PGF_00049889 | 14.04 | 312 | 60 | 0.795 | 0.120 | True | SSU ribosomal protein S3p (S3e) |
| PGF_00912265 | 14.03 | 505 | 60 | 0.624 | 0.242 | True | tRNA-dihydrouridine synthase DusB |
| PGF_01867628 | 14.00 | 364 | 60 | 0.734 | 0.065 | True | Heat-inducible transcription repressor HrcA |
| PGF_08905885 | 13.96 | 453 | 60 | 0.656 | 0.155 | True | Signal recognition particle receptor FtsY |
| PGF_00016357 | 13.86 | 241 | 60 | 0.893 | 0.010 | True | LSU ribosomal protein L1p (L10Ae) |
| PGF_00056897 | 13.80 | 274 | 60 | 0.834 | 0.028 | True | Thymidylate synthase (EC 2.1.1.45) |
| PGF_05053989 | 13.79 | 449 | 60 | 0.651 | 0.172 | True | ATP-dependent DNA ligase (EC 6.5.1.1) LigC |
| PGF_10461681 | 13.74 | 2050 | 60 | 0.303 | 0.472 | True | Ribonuclease E (EC 3.1.26.12) |
| PGF_04449783 | 13.70 | 362 | 60 | 0.720 | 0.099 | True | Pantothenate kinase (EC 2.7.1.33) |
| PGF_00008334 | 13.62 | 498 | 60 | 0.610 | 0.140 | True | Glutamyl-tRNA reductase (EC 1.2.1.70) |
| PGF_00692117 | 13.55 | 502 | 60 | 0.605 | 0.221 | True | Anion-transporting ATPase Rv3680 |
| PGF_00053513 | 13.51 | 360 | 60 | 0.712 | 0.071 | True | Solanesyl diphosphate synthase (EC 2.5.1.11) |
| PGF_07889681 | 13.43 | 415 | 60 | 0.659 | 0.166 | True | N-acetyl-gamma-glutamyl-phosphate reductase (EC 1.2.1.38) |
| PGF_02011760 | 13.37 | 279 | 60 | 0.800 | 0.085 | True | Imidazole glycerol phosphate synthase cyclase subunit |
| PGF_04574228 | 13.32 | 279 | 60 | 0.797 | 0.048 | True | Triosephosphate isomerase (EC 5.3.1.1) |
| PGF_00413192 | 13.30 | 382 | 60 | 0.680 | 0.149 | True | tRNA (adenine(58)-N(1))-methyltransferase (EC 2.1.1.220) |
| PGF_00420975 | 13.22 | 610 | 60 | 0.535 | 0.310 | True | D-inositol-3-phosphate glycosyltransferase (EC 2.4.1.250) |
| PGF_10387199 | 13.17 | 514 | 60 | 0.581 | 0.245 | True | DNA recombination and repair protein RecF |
| PGF_00048586 | 13.15 | 265 | 60 | 0.808 | 0.082 | True | Ribonuclease PH (EC 2.7.7.56) |
| PGF_00405726 | 13.14 | 363 | 60 | 0.690 | 0.087 | True | putative Adenosine kinase (EC 2.7.1.20) |
| PGF_00020173 | 13.10 | 350 | 60 | 0.700 | 0.104 | True | 2,3,4,5-tetrahydropyridine-2,6-dicarboxylate N-succinyltransferase (EC 2.3.1.117) |
| PGF_02959749 | 13.06 | 538 | 60 | 0.563 | 0.180 | True | Ribonuclease D (EC 3.1.26.3) |
| PGF_03062930 | 13.05 | 460 | 60 | 0.608 | 0.142 | True | Decaprenyl-phosphate N-acetylglucosaminephosphotransferase (EC 2.7.8.35) |
| PGF_05165078 | 13.04 | 341 | 60 | 0.706 | 0.089 | True | Methionyl-tRNA formyltransferase (EC 2.1.2.9) |
| PGF_00052943 | 13.01 | 375 | 60 | 0.672 | 0.173 | True | Site-specific tyrosine recombinase XerD |
| PGF_01745391 | 13.00 | 442 | 60 | 0.618 | 0.182 | True | Cystathionine gamma-synthase (EC 2.5.1.48) |
| PGF_00037867 | 12.98 | 305 | 60 | 0.743 | 0.087 | True | Proteasome subunit beta (EC 3.4.25.1), bacterial |
| PGF_04999088 | 12.92 | 598 | 60 | 0.528 | 0.245 | True | Uncharacterized protein SCO5199 |
| PGF_00001910 | 12.90 | 310 | 60 | 0.733 | 0.092 | True | Uncharacterized protien SCO2557 |
| PGF_02191019 | 12.87 | 359 | 60 | 0.679 | 0.111 | True | Thiamine-monophosphate kinase (EC 2.7.4.16) |
| PGF_00413333 | 12.85 | 491 | 60 | 0.580 | 0.222 | True | Lipid II:glycine glycyltransferase (EC 2.3.2.16) |
| PGF_00063974 | 12.84 | 393 | 60 | 0.648 | 0.113 | True | UDP-N-acetylenolpyruvoylglucosamine reductase (EC 1.3.1.98) |
| PGF_00014855 | 12.74 | 306 | 60 | 0.728 | 0.074 | True | ATP phosphoribosyltransferase (EC 2.4.2.17) => HisGl |
| PGF_00033950 | 12.74 | 274 | 60 | 0.770 | 0.096 | True | Phosphoribosylformimino-5-aminoimidazole carboxamide ribotide isomerase (EC 5.3.1.16) @ Acting phosphoribosylanthranilate isomerase (EC 5.3.1.24) |
| PGF_02406951 | 12.73 | 679 | 60 | 0.488 | 0.284 | True | Wax ester synthase/acyl-CoA:diacylglycerol acyltransferase; Diacyglycerol O-acyltransferase (EC 2.3.1.20) |
| PGF_02390924 | 12.72 | 397 | 60 | 0.639 | 0.176 | True | 16S rRNA (cytosine(1402)-N(4))-methyltransferase (EC 2.1.1.199) |
| PGF_00015517 | 12.72 | 475 | 60 | 0.584 | 0.162 | True | Iron-sulfur cluster assembly protein SufD |
| PGF_00066906 | 12.67 | 357 | 60 | 0.671 | 0.122 | True | Aspartate carbamoyltransferase (EC 2.1.3.2) |
| PGF_03788368 | 12.66 | 306 | 60 | 0.724 | 0.049 | True | RNase adapter protein RapZ |
| PGF_06755829 | 12.64 | 429 | 60 | 0.610 | 0.093 | True | DNA polymerase III delta prime subunit (EC 2.7.7.7) |
| PGF_02895544 | 12.63 | 232 | 60 | 0.829 | 0.039 | True | TrkA-like protein |
| PGF_00403095 | 12.61 | 214 | 60 | 0.862 | 0.004 | True | Uracil phosphoribosyltransferase (EC 2.4.2.9) |
| PGF_00027514 | 12.59 | 478 | 60 | 0.576 | 0.171 | True | N-acetylornithine aminotransferase (EC 2.6.1.11) |
| PGF_02463284 | 12.59 | 202 | 60 | 0.886 | 0.018 | True | Recombination protein RecR |
| PGF_06684654 | 12.57 | 389 | 60 | 0.637 | 0.132 | True | UDP-glucose 4-epimerase (EC 5.1.3.2) |
| PGF_00057381 | 12.55 | 299 | 60 | 0.726 | 0.132 | True | Trans,polycis-decaprenyl diphosphate synthase (EC 2.5.1.86) |
| PGF_05767868 | 12.54 | 265 | 60 | 0.770 | 0.035 | True | Probable transcriptional regulatory protein YebC |
| PGF_00049827 | 12.53 | 348 | 60 | 0.671 | 0.161 | True | SSU rRNA (adenine(1518)-N(6)/adenine(1519)-N(6))-dimethyltransferase (EC 2.1.1.182) |
| PGF_04094270 | 12.51 | 231 | 60 | 0.823 | 0.045 | True | TrkA-like protein |
| PGF_02450432 | 12.44 | 333 | 60 | 0.682 | 0.111 | True | Phosphatidate cytidylyltransferase (EC 2.7.7.41) |
| PGF_00016431 | 12.42 | 230 | 60 | 0.819 | 0.042 | True | LSU ribosomal protein L3p (L3e) |
| PGF_02866866 | 12.41 | 449 | 60 | 0.586 | 0.182 | True | Coproporphyrin ferrochelatase (EC 4.99.1.9) |
| PGF_00049893 | 12.35 | 202 | 60 | 0.869 | 0.010 | True | SSU ribosomal protein S4p (S9e) @ SSU ribosomal protein S4p (S9e), zinc-independent |
| PGF_00016613 | 12.32 | 415 | 60 | 0.605 | 0.181 | True | Lactyl (2) diphospho-(5')guanosine:7,8-didemethyl-8-hydroxy-5-deazariboflavin 2-phospho-L-lactate transferase (EC 2.7.8.28) |
| PGF_12831525 | 12.28 | 266 | 60 | 0.753 | 0.070 | True | SAM-dependent methyltransferase SCO2317, type 11 |
| PGF_01033770 | 12.26 | 417 | 60 | 0.600 | 0.164 | True | Dihydroorotate dehydrogenase (quinone) (EC 1.3.5.2) |
| PGF_00422922 | 12.26 | 203 | 60 | 0.860 | 0.057 | True | Deoxycytidine triphosphate deaminase (EC 3.5.4.30) (dUMP-forming) |
| PGF_00038986 | 12.23 | 326 | 60 | 0.678 | 0.169 | True | Purine nucleoside phosphorylase (EC 2.4.2.1) |
| PGF_00014051 | 12.21 | 293 | 60 | 0.713 | 0.101 | True | Indole-3-glycerol phosphate synthase (EC 4.1.1.48) |
| PGF_00415308 | 12.18 | 326 | 60 | 0.675 | 0.144 | True | 23S rRNA (cytidine(1920)-2'-O)-methyltransferase (EC 2.1.1.226) @ 16S rRNA (cytidine(1409)-2'-O)-methyltransferase (EC 2.1.1.227) |
| PGF_08151051 | 12.13 | 400 | 60 | 0.607 | 0.253 | True | Histidinol-phosphatase [alternative form] (EC 3.1.3.15) |
| PGF_10464361 | 12.11 | 259 | 60 | 0.753 | 0.096 | True | Two component system response regulator MtrA |
| PGF_00001331 | 12.02 | 267 | 60 | 0.736 | 0.139 | True | Endonuclease NucS |
| PGF_02617708 | 11.99 | 322 | 60 | 0.668 | 0.093 | True | A/G-specific adenine glycosylase (EC 3.2.2.-) |
| PGF_00049896 | 11.98 | 225 | 60 | 0.799 | 0.082 | True | SSU ribosomal protein S5p (S2e) |
| PGF_00769755 | 11.96 | 597 | 60 | 0.490 | 0.257 | False | 16S rRNA (cytosine(967)-C(5))-methyltransferase (EC 2.1.1.176) |
| PGF_08005402 | 11.96 | 372 | 60 | 0.620 | 0.130 | True | FIG002813: LPPG:FO 2-phospho-L-lactate transferase like, CofD-like |
| PGF_08125688 | 11.95 | 412 | 60 | 0.589 | 0.187 | True | Anion-transporting ATPase Rv3679 |
| PGF_00016443 | 11.88 | 206 | 60 | 0.828 | 0.076 | True | LSU ribosomal protein L5p (L11e) |
| PGF_00006100 | 11.87 | 427 | 60 | 0.574 | 0.259 | True | tRNA-modifying protein YgfZ |
| PGF_08421732 | 11.86 | 406 | 60 | 0.589 | 0.210 | True | NAD kinase (EC 2.7.1.23) |
| PGF_03374975 | 11.78 | 420 | 60 | 0.575 | 0.243 | True | O-succinylbenzoate synthase (EC 4.2.1.113) |
| PGF_04807486 | 11.77 | 345 | 60 | 0.634 | 0.103 | True | tRNA dimethylallyltransferase (EC 2.5.1.75) |
| PGF_00024232 | 11.77 | 517 | 60 | 0.518 | 0.291 | True | N-succinyl-L,L-diaminopimelate aminotransferase (EC 2.6.1.17), type 2 |
| PGF_00549380 | 11.75 | 333 | 60 | 0.644 | 0.101 | True | 4-hydroxy-tetrahydrodipicolinate synthase (EC 4.3.3.7) |
| PGF_03072132 | 11.70 | 173 | 60 | 0.889 | 0.068 | True | CarD-like transcriptional regulator |
| PGF_00016444 | 11.70 | 183 | 60 | 0.865 | 0.017 | True | LSU ribosomal protein L6p (L9e) |
| PGF_08394843 | 11.65 | 268 | 60 | 0.712 | 0.111 | True | Endonuclease III (EC 4.2.99.18) |
| PGF_03799365 | 11.64 | 188 | 60 | 0.849 | 0.005 | True | Translation elongation factor P |
| PGF_00015467 | 11.62 | 268 | 60 | 0.710 | 0.143 | True | Iron-dependent repressor IdeR/DtxR |
| PGF_05091456 | 11.62 | 318 | 60 | 0.651 | 0.052 | True | Uncharacterized inner membrane protein RarD |
| PGF_02897933 | 11.60 | 290 | 60 | 0.681 | 0.158 | True | Nitrogen metabolism regulator GlnR, OmpR family |
| PGF_00049904 | 11.60 | 156 | 60 | 0.929 | 0.000 | True | SSU ribosomal protein S7p (S5e) |
| PGF_05580933 | 11.59 | 320 | 60 | 0.648 | 0.126 | True | Peptide chain release factor N(5)-glutamine methyltransferase (EC 2.1.1.297) |
| PGF_00057483 | 11.58 | 997 | 60 | 0.367 | 0.362 | True | Transcription termination factor Rho |
| PGF_00417658 | 11.51 | 571 | 60 | 0.482 | 0.347 | True | 3-dehydroquinate synthase (EC 4.2.3.4) |
| PGF_00419621 | 11.47 | 253 | 60 | 0.721 | 0.072 | True | Coproheme decarboxylase HemQ (no EC) |
| PGF_00413295 | 11.37 | 330 | 60 | 0.626 | 0.132 | True | tRNA pseudouridine(55) synthase (EC 5.4.99.25) |
| PGF_00047155 | 11.34 | 262 | 60 | 0.700 | 0.138 | True | Redox-sensing transcriptional repressor Rex |
| PGF_07479808 | 11.33 | 394 | 60 | 0.571 | 0.220 | True | Ketopantoate reductase PanG (EC 1.1.1.169) |
| PGF_03475877 | 11.30 | 475 | 60 | 0.519 | 0.290 | True | Chromosome (plasmid) partitioning protein ParB |
| PGF_07695531 | 11.29 | 144 | 60 | 0.941 | 0.013 | True | LSU ribosomal protein L11p (L12e) |
| PGF_10348836 | 11.28 | 336 | 60 | 0.615 | 0.178 | True | NADH-ubiquinone oxidoreductase chain J (EC 1.6.5.3) |
| PGF_02923127 | 11.28 | 265 | 60 | 0.693 | 0.102 | True | Uridylate kinase (EC 2.7.4.22) |
| PGF_04213876 | 11.26 | 298 | 60 | 0.652 | 0.136 | True | Pantothenate kinase type III, CoaX-like (EC 2.7.1.33) |
| PGF_02930420 | 11.25 | 480 | 60 | 0.513 | 0.243 | True | Ubiquinol-cytochrome C reductase iron-sulfur subunit (EC 1.10.2.2) |
| PGF_03647550 | 11.23 | 163 | 60 | 0.880 | 0.000 | True | UPF0234 protein Yitk |
| PGF_00423089 | 11.20 | 490 | 60 | 0.506 | 0.283 | True | Diaminohydroxyphosphoribosylaminopyrimidine deaminase (EC 3.5.4.26) / 5-amino-6-(5-phosphoribosylamino)uracil reductase (EC 1.1.1.193) |
| PGF_06461498 | 11.20 | 333 | 60 | 0.614 | 0.170 | True | Orotidine 5'-phosphate decarboxylase (EC 4.1.1.23) |
| PGF_00016343 | 11.19 | 139 | 60 | 0.949 | 0.000 | True | LSU ribosomal protein L16p (L10e) |
| PGF_04370656 | 11.18 | 446 | 60 | 0.530 | 0.298 | True | Chromosome (plasmid) partitioning protein ParA |
| PGF_00013803 | 11.16 | 231 | 60 | 0.734 | 0.120 | True | Imidazoleglycerol-phosphate dehydratase (EC 4.2.1.19) |
| PGF_00062023 | 11.16 | 350 | 60 | 0.596 | 0.228 | True | Tryptophan synthase alpha chain (EC 4.2.1.20) |
| PGF_00048829 | 11.15 | 327 | 60 | 0.617 | 0.200 | True | LSU rRNA pseudouridine(2605) synthase (EC 5.4.99.22) |
| PGF_00054323 | 11.13 | 222 | 60 | 0.747 | 0.074 | True | YciO protein, TsaC/YrdC paralog |
| PGF_01213071 | 11.12 | 264 | 60 | 0.685 | 0.162 | True | LSU ribosomal protein L10p (P0) |
| PGF_00026615 | 11.12 | 533 | 60 | 0.482 | 0.403 | True | N-acetylglutamate kinase (EC 2.7.2.8) |
| PGF_06216244 | 11.11 | 306 | 60 | 0.635 | 0.200 | True | DNA recombination and repair protein RecO |
| PGF_03439827 | 11.10 | 716 | 60 | 0.415 | 0.413 | True | Precorrin-2 oxidase (EC 1.3.1.76) @ Sirohydrochlorin ferrochelatase activity of CysG (EC 4.99.1.4) / Uroporphyrinogen-III methyltransferase (EC 2.1.1.107) |
| PGF_00006724 | 11.07 | 294 | 60 | 0.646 | 0.045 | True | GCN5-related N-acetyltransferase, FIGfam019367 |
| PGF_03591205 | 11.05 | 660 | 60 | 0.430 | 0.338 | True | Serine/threonine phosphatase PPP (EC 3.1.3.16) |
| PGF_00048926 | 11.03 | 185 | 60 | 0.811 | 0.007 | True | Ribosome recycling factor |
| PGF_09087715 | 10.99 | 382 | 60 | 0.562 | 0.145 | True | tRNA(Ile)-lysidine synthetase (EC 6.3.4.19) |
| PGF_00049837 | 10.98 | 135 | 60 | 0.945 | 0.021 | True | SSU ribosomal protein S11p (S14e) |
| PGF_00013490 | 10.88 | 210 | 60 | 0.751 | 0.125 | True | Hypoxanthine-guanine phosphoribosyltransferase (EC 2.4.2.8) |
| PGF_01382519 | 10.87 | 222 | 60 | 0.730 | 0.069 | True | Ribosome hibernation promoting factor Hpf |
| PGF_02787494 | 10.77 | 221 | 60 | 0.725 | 0.064 | True | Phosphate transport regulator (distant homolog of PhoU) |
| PGF_00016340 | 10.74 | 122 | 60 | 0.973 | 0.000 | True | LSU ribosomal protein L14p (L23e) |
| PGF_00413290 | 10.68 | 327 | 60 | 0.590 | 0.151 | True | tRNA pseudouridine(38-40) synthase (EC 5.4.99.12) |
| PGF_00025215 | 10.62 | 197 | 60 | 0.757 | 0.080 | True | Acetolactate synthase small subunit (EC 2.2.1.6) |
| PGF_00001849 | 10.53 | 180 | 60 | 0.785 | 0.056 | True | FIG01122152: hypothetical protein |
| PGF_00049906 | 10.50 | 140 | 60 | 0.887 | 0.025 | True | SSU ribosomal protein S8p (S15Ae) |
| PGF_00984073 | 10.49 | 243 | 60 | 0.673 | 0.103 | True | Phosphate transport system regulatory protein PhoU |
| PGF_00016342 | 10.45 | 147 | 60 | 0.862 | 0.006 | True | LSU ribosomal protein L15p (L27Ae) |
| PGF_10123167 | 10.44 | 193 | 60 | 0.752 | 0.020 | True | 2-amino-4-hydroxy-6-hydroxymethyldihydropteridine pyrophosphokinase (EC 2.7.6.3) |
| PGF_00049840 | 10.41 | 125 | 60 | 0.931 | 0.008 | True | SSU ribosomal protein S13p (S18e) |
| PGF_03189552 | 10.36 | 222 | 60 | 0.696 | 0.102 | True | Holliday junction ATP-dependent DNA helicase RuvA (EC 3.6.4.12) |
| PGF_04828114 | 10.36 | 389 | 60 | 0.525 | 0.218 | True | Quinolinate phosphoribosyltransferase [decarboxylating] (EC 2.4.2.19) |
| PGF_05843198 | 10.34 | 274 | 60 | 0.625 | 0.125 | True | FIG017108: hypothetical protein |
| PGF_08432396 | 10.31 | 261 | 60 | 0.638 | 0.201 | True | Nicotinate-nucleotide adenylyltransferase (EC 2.7.7.18) |
| PGF_06180597 | 10.29 | 147 | 60 | 0.848 | 0.000 | True | LSU ribosomal protein L13p (L13Ae) |
| PGF_00336478 | 10.27 | 251 | 60 | 0.648 | 0.176 | True | Two-component transcriptional response regulator PdtaR, LuxR family |
| PGF_06941403 | 10.19 | 137 | 60 | 0.870 | 0.100 | True | SSU ribosomal protein S12p (S23e) |
| PGF_00016358 | 10.18 | 132 | 60 | 0.886 | 0.025 | True | LSU ribosomal protein L20p |
| PGF_01447139 | 10.16 | 244 | 60 | 0.650 | 0.203 | True | ATP:Cob(I)alamin adenosyltransferase (EC 2.5.1.17) |
| PGF_00523822 | 10.12 | 418 | 60 | 0.495 | 0.270 | True | Pantoate--beta-alanine ligase (EC 6.3.2.1) |
| PGF_00853991 | 10.11 | 357 | 60 | 0.535 | 0.244 | True | Diaminopimelate epimerase (EC 5.1.1.7) |
| PGF_00504549 | 10.06 | 279 | 60 | 0.602 | 0.219 | True | O-methyltransferase Rv1220c |
| PGF_00049909 | 10.06 | 194 | 60 | 0.722 | 0.133 | True | SSU ribosomal protein S9p (S16e) |
| PGF_00049828 | 10.04 | 102 | 60 | 0.994 | 0.000 | True | SSU ribosomal protein S10p (S20e) |
| PGF_00033237 | 9.93 | 777 | 60 | 0.356 | 0.547 | True | Phosphate starvation-inducible protein PhoH, predicted ATPase |
| PGF_00065286 | 9.92 | 227 | 60 | 0.658 | 0.106 | True | Uncharacterized protein Q1 colocalized with Q |
| PGF_00033968 | 9.91 | 228 | 60 | 0.656 | 0.111 | True | Phosphoribosylglycinamide formyltransferase (EC 2.1.2.2) |
| PGF_00689961 | 9.88 | 237 | 60 | 0.642 | 0.165 | True | Guanylate kinase (EC 2.7.4.8) |
| PGF_03790040 | 9.85 | 360 | 60 | 0.519 | 0.293 | True | Ribonuclease III (EC 3.1.26.3) |
| PGF_00956915 | 9.80 | 481 | 60 | 0.447 | 0.292 | True | O-succinylbenzoic acid--CoA ligase (EC 6.2.1.26) |
| PGF_09288314 | 9.74 | 352 | 60 | 0.519 | 0.278 | True | RNA methyltransferase, TrmH family |
| PGF_01187824 | 9.74 | 303 | 60 | 0.560 | 0.258 | True | Demethylmenaquinone methyltransferase (EC 2.1.1.163) |
| PGF_00045787 | 9.67 | 238 | 60 | 0.627 | 0.157 | True | Pyridoxal 5'-phosphate synthase (glutamine hydrolyzing), glutaminase subunit (EC 4.3.3.6) |
| PGF_04788810 | 9.65 | 267 | 60 | 0.591 | 0.250 | True | Peptidyl-tRNA hydrolase (EC 3.1.1.29) |
| PGF_05172785 | 9.63 | 192 | 60 | 0.695 | 0.083 | True | N5-carboxyaminoimidazole ribonucleotide mutase (EC 5.4.99.18) |
| PGF_00423533 | 9.63 | 358 | 60 | 0.509 | 0.155 | True | 4-diphosphocytidyl-2-C-methyl-D-erythritol kinase (EC 2.7.1.148) |
| PGF_03751076 | 9.63 | 262 | 60 | 0.595 | 0.204 | True | Riboflavin synthase eubacterial/eukaryotic (EC 2.5.1.9) |
| PGF_00015543 | 9.61 | 291 | 60 | 0.563 | 0.212 | True | Iron-sulfur cluster regulator SufR |
| PGF_12753784 | 9.60 | 283 | 60 | 0.571 | 0.260 | True | Transcriptional regulator SCO5170, AcrR family |
| PGF_06366599 | 9.59 | 356 | 60 | 0.508 | 0.178 | True | Probable (3R)-hydroxyacyl-CoA dehydratase HtdX |
| PGF_07051332 | 9.53 | 286 | 60 | 0.563 | 0.207 | True | LSU ribosomal protein L25p |
| PGF_00026362 | 9.51 | 261 | 60 | 0.589 | 0.200 | True | Nucleoside 5-triphosphatase RdgB (dHAPTP, dITP, XTP-specific) (EC 3.6.1.66) |
| PGF_08843714 | 9.50 | 257 | 60 | 0.592 | 0.211 | True | Septum formation protein Maf |
| PGF_02455692 | 9.48 | 291 | 60 | 0.556 | 0.210 | True | Cytidylate kinase (EC 2.7.4.25) |
| PGF_03174068 | 9.47 | 434 | 60 | 0.454 | 0.352 | True | Transcription antitermination protein NusG |
| PGF_00000568 | 9.46 | 135 | 60 | 0.814 | 0.056 | True | FIG00820327: hypothetical protein |
| PGF_00024692 | 9.37 | 366 | 60 | 0.490 | 0.326 | True | NADH-ubiquinone oxidoreductase chain E (EC 1.6.5.3) |
| PGF_00016445 | 9.35 | 141 | 60 | 0.788 | 0.084 | True | LSU ribosomal protein L7p/L12p (P1/P2) |
| PGF_00016452 | 9.32 | 151 | 60 | 0.759 | 0.020 | True | LSU ribosomal protein L9p |
| PGF_02792560 | 9.24 | 256 | 60 | 0.577 | 0.179 | True | tRNA threonylcarbamoyladenosine biosynthesis protein TsaB |
| PGF_03295678 | 9.23 | 244 | 60 | 0.591 | 0.147 | True | CDP-diacylglycerol--glycerol-3-phosphate 3-phosphatidyltransferase (EC 2.7.8.5) |
| PGF_00220548 | 9.22 | 178 | 60 | 0.691 | 0.114 | True | Thiol peroxidase, Bcp-type (EC 1.11.1.15) |
| PGF_10401812 | 9.21 | 403 | 60 | 0.459 | 0.451 | True | Thymidine kinase (EC 2.7.1.21) |
| PGF_02517283 | 9.20 | 581 | 60 | 0.382 | 0.487 | True | Segregation and condensation protein A |
| PGF_00016353 | 9.14 | 129 | 60 | 0.805 | 0.014 | True | LSU ribosomal protein L18p (L5e) |
| PGF_00876943 | 9.09 | 635 | 60 | 0.361 | 0.528 | True | 1,4-dihydroxy-2-naphthoate polyprenyltransferase (EC 2.5.1.74) |
| PGF_05608844 | 9.07 | 110 | 60 | 0.865 | 0.000 | True | Small basic protein Sbp |
| PGF_00020361 | 9.07 | 182 | 60 | 0.672 | 0.149 | True | Metal-dependent hydrolase YbeY, involved in rRNA and/or ribosome maturation and assembly |
| PGF_04618589 | 9.06 | 201 | 60 | 0.639 | 0.148 | True | Glycogen accumulation regulator GarA |
| PGF_00426932 | 8.99 | 179 | 60 | 0.672 | 0.106 | True | 6,7-dimethyl-8-ribityllumazine synthase (EC 2.5.1.78) |
| PGF_00041788 | 8.97 | 189 | 60 | 0.652 | 0.187 | True | Putative iron-sulfur cluster assembly scaffold protein for SUF system, SufE2 |
| PGF_02569889 | 8.95 | 238 | 60 | 0.580 | 0.149 | True | hypothetical protein |
| PGF_00066124 | 8.86 | 258 | 60 | 0.552 | 0.265 | True | Pyrimidine operon regulatory protein PyrR |
| PGF_07133621 | 8.80 | 224 | 60 | 0.588 | 0.149 | True | 16S rRNA (guanine(966)-N(2))-methyltransferase (EC 2.1.1.171) |
| PGF_06626131 | 8.80 | 207 | 60 | 0.611 | 0.209 | True | 2-C-methyl-D-erythritol 2,4-cyclodiphosphate synthase (EC 4.6.1.12) |
| PGF_00048643 | 8.78 | 201 | 60 | 0.620 | 0.193 | True | Ribonucleotide reductase transcriptional regulator NrdR |
| PGF_00016368 | 8.76 | 183 | 60 | 0.647 | 0.208 | True | LSU ribosomal protein L22p (L17e) |
| PGF_05296102 | 8.75 | 363 | 60 | 0.459 | 0.245 | True | Acetyl-CoA:Cys-GlcN-Ins acetyltransferase, mycothiol synthase MshD (EC 2.3.1.189) |
| PGF_00066839 | 8.73 | 84 | 60 | 0.952 | 0.011 | True | WhiB-like transcription regulator |
| PGF_05880970 | 8.72 | 318 | 60 | 0.489 | 0.353 | True | Cardiolipin synthase (CMP-forming), eukaryotic type Cls-II (EC 2.7.8.41) |
| PGF_00420001 | 8.69 | 187 | 60 | 0.635 | 0.202 | True | CysO-cysteine peptidase |
| PGF_08197987 | 8.62 | 159 | 60 | 0.684 | 0.244 | True | Iron-sulfur cluster insertion protein SCO2161 |
| PGF_08518355 | 8.61 | 355 | 60 | 0.457 | 0.244 | True | ATP synthase F0 sector subunit a (EC 3.6.3.14) |
| PGF_00049847 | 8.60 | 96 | 60 | 0.878 | 0.052 | True | SSU ribosomal protein S15p (S13e) |
| PGF_00021018 | 8.52 | 188 | 60 | 0.621 | 0.187 | True | Methylmalonyl-CoA epimerase (EC 5.1.99.1) @ Ethylmalonyl-CoA epimerase |
| PGF_02031353 | 8.46 | 263 | 60 | 0.522 | 0.294 | True | FIG01269488: protein, clustered with ribosomal protein L32p |
| PGF_10483430 | 8.46 | 203 | 60 | 0.594 | 0.212 | True | Methylated-DNA--protein-cysteine methyltransferase (EC 2.1.1.63) |
| PGF_01430214 | 8.39 | 207 | 60 | 0.583 | 0.225 | True | Deoxyuridine 5'-triphosphate nucleotidohydrolase (EC 3.6.1.23) |
| PGF_00008838 | 8.36 | 300 | 60 | 0.483 | 0.221 | True | 16S rRNA (guanine(527)-N(7))-methyltransferase (EC 2.1.1.170) |
| PGF_05357708 | 8.34 | 231 | 60 | 0.549 | 0.172 | True | DNA-3-methyladenine glycosylase (EC 3.2.2.20) |
| PGF_03518570 | 8.29 | 147 | 60 | 0.684 | 0.052 | True | Nucleoside diphosphate kinase (EC 2.7.4.6) |
| PGF_00016385 | 8.29 | 87 | 60 | 0.888 | 0.033 | True | LSU ribosomal protein L27p |
| PGF_01420802 | 8.27 | 150 | 60 | 0.675 | 0.054 | True | Transcriptional regulator MraZ |
| PGF_00016377 | 8.23 | 141 | 60 | 0.693 | 0.159 | True | LSU ribosomal protein L24p (L26e) |
| PGF_03642879 | 8.13 | 146 | 60 | 0.673 | 0.084 | True | Cytochrome c oxidase polypeptide IV (EC 1.9.3.1) |
| PGF_06701488 | 8.10 | 256 | 60 | 0.506 | 0.274 | True | RNA-binding protein Jag |
| PGF_00418586 | 8.05 | 165 | 60 | 0.626 | 0.201 | True | Cold shock protein of CSP family => SCO4325 |
| PGF_03753407 | 8.03 | 542 | 60 | 0.345 | 0.432 | True | tRNA threonylcarbamoyladenosine biosynthesis protein TsaE |
| PGF_00036456 | 8.02 | 230 | 60 | 0.529 | 0.330 | True | Predicted transcriptional regulator of sulfate adenylyltransferase, Rrf2 family |
| PGF_05906360 | 7.88 | 131 | 60 | 0.688 | 0.187 | True | DNA-directed RNA polymerase omega subunit (EC 2.7.7.6) |
| PGF_00000494 | 7.78 | 281 | 60 | 0.464 | 0.361 | True | FIG00816212: Putative membrane protein |
| PGF_00049860 | 7.77 | 127 | 60 | 0.689 | 0.263 | True | SSU ribosomal protein S19p (S15e) |
| PGF_00049854 | 7.77 | 99 | 60 | 0.781 | 0.068 | True | SSU ribosomal protein S17p (S11e) |
| PGF_00650730 | 7.74 | 107 | 60 | 0.748 | 0.135 | True | Transcriptional regulator, AsnC family |
| PGF_03277153 | 7.73 | 114 | 60 | 0.724 | 0.127 | True | Sulfur metabolism protein SseC |
| PGF_06661068 | 7.68 | 189 | 60 | 0.559 | 0.318 | True | PaaD-like protein (DUF59) involved in Fe-S cluster assembly |
| PGF_00049842 | 7.65 | 61 | 60 | 0.980 | 0.000 | True | SSU ribosomal protein S14p (S29e) @ SSU ribosomal protein S14p (S29e), zinc-dependent |
| PGF_09581668 | 7.64 | 126 | 60 | 0.681 | 0.223 | True | Heat shock protein 10 kDa family chaperone GroES |
| PGF_01172431 | 7.61 | 96 | 60 | 0.776 | 0.092 | True | Phosphoribosyl-ATP pyrophosphatase (EC 3.6.1.31) |
| PGF_00057353 | 7.55 | 145 | 60 | 0.627 | 0.222 | True | Anti-sigma factor antagonist BldG |
| PGF_00016346 | 7.55 | 334 | 60 | 0.413 | 0.397 | True | LSU ribosomal protein L17p |
| PGF_00435430 | 7.53 | 163 | 60 | 0.590 | 0.232 | True | Ribosomal silencing factor RsfA |
| PGF_04978890 | 7.44 | 129 | 60 | 0.655 | 0.214 | True | LSU ribosomal protein L21p |
| PGF_00016387 | 7.37 | 61 | 60 | 0.943 | 0.000 | True | LSU ribosomal protein L28p @ LSU ribosomal protein L28p, zinc-dependent |
| PGF_01682834 | 7.32 | 243 | 60 | 0.470 | 0.318 | True | Bacterial ribosome SSU maturation protein RimP |
| PGF_00049901 | 7.20 | 126 | 60 | 0.642 | 0.229 | True | SSU ribosomal protein S6p |
| PGF_00934722 | 7.13 | 81 | 60 | 0.792 | 0.037 | True | Protein translocase membrane subunit SecG |
| PGF_09402727 | 7.10 | 530 | 60 | 0.309 | 0.494 | True | Molybdenum ABC transporter permease protein ModB |
| PGF_00960330 | 7.04 | 123 | 60 | 0.635 | 0.184 | True | ATP-dependent Clp protease adaptor protein ClpS |
| PGF_00055165 | 7.03 | 116 | 60 | 0.652 | 0.195 | True | Sulfur/cysteine carrier protein CysO |
| PGF_02899131 | 6.94 | 91 | 60 | 0.727 | 0.118 | True | LSU ribosomal protein L29p (L35e) |
| PGF_06609275 | 6.87 | 56 | 60 | 0.918 | 0.000 | True | LSU ribosomal protein L33p @ LSU ribosomal protein L33p, zinc-dependent |
| PGF_02776739 | 6.68 | 179 | 60 | 0.499 | 0.335 | True | Phosphoribosyl-AMP cyclohydrolase (EC 3.5.4.19) |
| PGF_03538342 | 6.59 | 131 | 60 | 0.576 | 0.260 | True | Transcriptional regulator, WhiB family |
| PGF_08210457 | 6.57 | 65 | 60 | 0.815 | 0.001 | True | Prokaryotic ubiquitin-like protein Pup |
| PGF_05032476 | 6.25 | 154 | 60 | 0.504 | 0.253 | True | Uncharacterized protein MSMEG_5817 |
| PGF_06060005 | 6.05 | 78 | 60 | 0.685 | 0.133 | True | Excisionase-like protein SCO3328 |
| PGF_00419702 | 5.91 | 597 | 60 | 0.242 | 0.685 | False | Crossover junction endodeoxyribonuclease RuvC (EC 3.1.22.4) |
| PGF_04972523 | 5.91 | 180 | 60 | 0.440 | 0.367 | True | Ferredoxin, 2Fe-2S |
| PGF_05770273 | 5.89 | 109 | 60 | 0.565 | 0.307 | True | LSU ribosomal protein L31p @ LSU ribosomal protein L31p, zinc-dependent |
| PGF_00016424 | 5.78 | 37 | 60 | 0.951 | 0.000 | True | LSU ribosomal protein L36p @ LSU ribosomal protein L36p, zinc-dependent |
| PGF_12924000 | 5.66 | 84 | 60 | 0.617 | 0.335 | True | Uncharacterized protein MSMEG_5081 |
| PGF_03038172 | 5.62 | 124 | 60 | 0.504 | 0.330 | True | Mycoredoxin (EC 1.20.4.3) |
| PGF_00426285 | 5.59 | 535 | 60 | 0.242 | 0.663 | True | FIG004853: possible toxin to DivIC |
| PGF_03751658 | 5.37 | 125 | 60 | 0.480 | 0.300 | True | Membrane protein insertion efficiency factor YidD |
| PGF_05076104 | 4.95 | 101 | 60 | 0.493 | 0.291 | True | Exodeoxyribonuclease VII small subunit (EC 3.1.11.6) |
| PGF_00016418 | 4.54 | 86 | 60 | 0.490 | 0.468 | True | LSU ribosomal protein L34p |
