## Supplementary figures and images for "Biotechnological potential of aromatic compounds–utilizing bacteria from Brazilian caves, including a novel cave *Nocardioides sp*. SF1"

### Figure S1

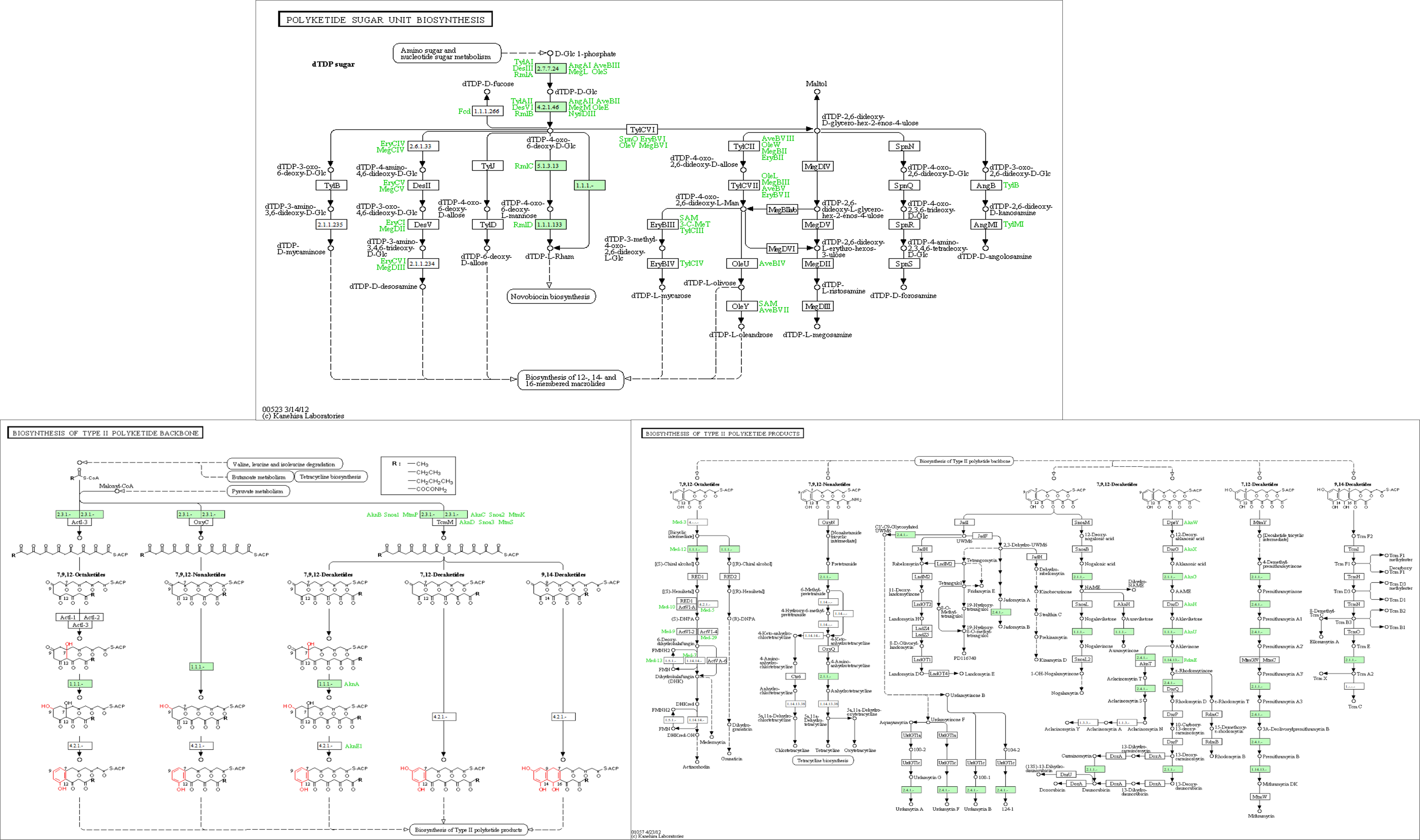
