## Supplementary material for "Biotechnological potential of aromatic compounds–utilizing bacteria from Brazilian caves, including a novel cave *Nocardioides sp*. SF1": Figure S1

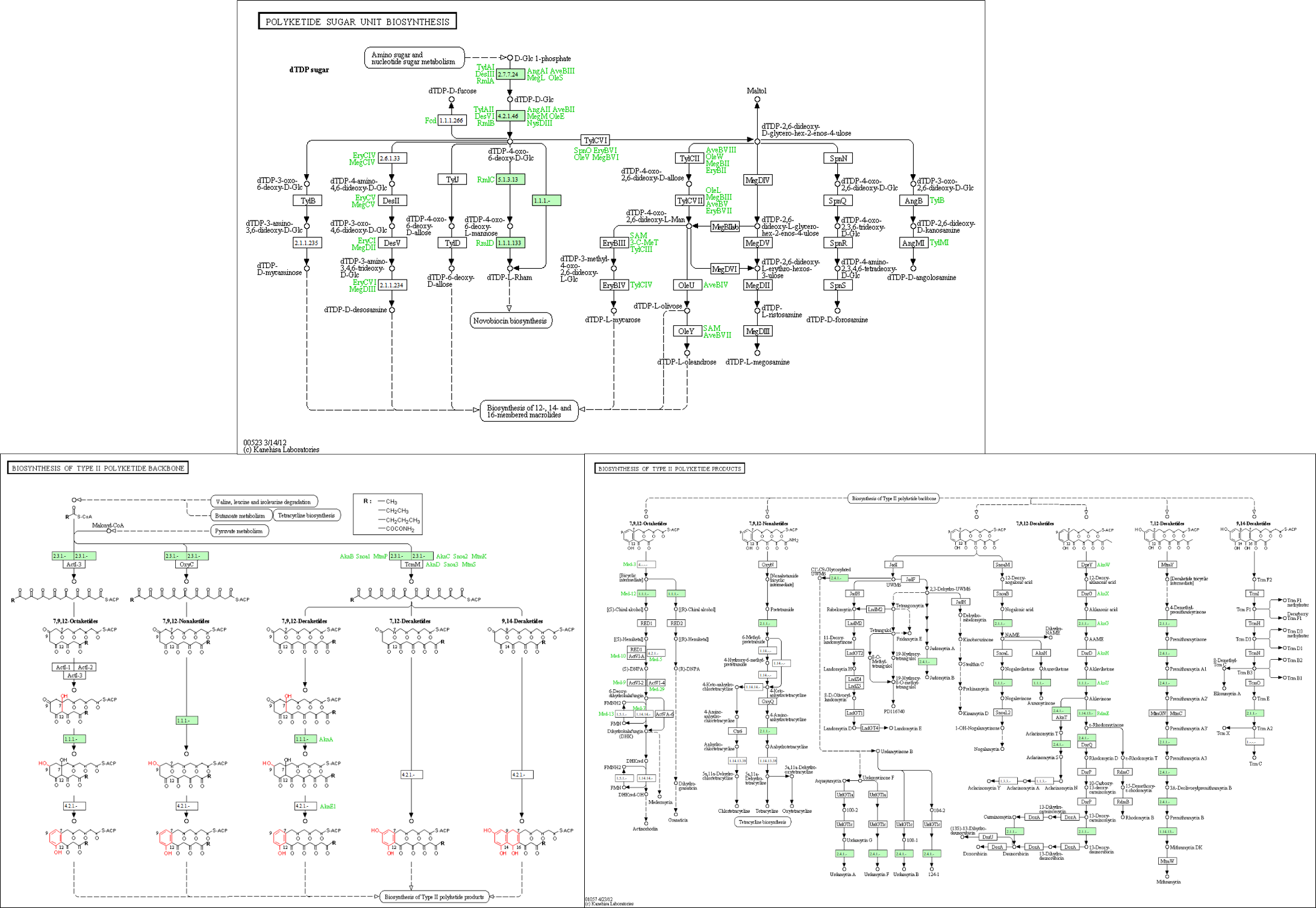

Figure S1 - Predicted type II polyketide biosynthetic pathways in the genome of *Nocardioides* sp. SF1. The top panel shows the “polyketide sugar unit biosynthesis” map, highlighting the enzymes present (in green) that are involved in the formation of activated dTDP‑sugars, precursors of rare glycosyl units. The bottom panels represent, on the left, the “biosynthesis of type II polyketide backbone”, indicating the type II PKS complex enzymes detected in the genome, and on the right, the “biosynthesis of type II polyketide products”, linking these enzymes to putative final aromatic structures decorated with sugars
